# The effect of statistical normalisation on network propagation scores

**DOI:** 10.1101/2020.01.20.911842

**Authors:** Sergio Picart-Armada, Wesley K. Thompson, Alfonso Buil, Alexandre Perera-Lluna

## Abstract

**Motivation:** Network diffusion and label propagation are fundamental tools in computational biology, with applications like gene-disease association, protein function prediction and module discovery. More recently, several publications have introduced a permutation analysis after the propagation process, due to concerns that network topology can bias diffusion scores. This opens the question of the statistical properties and the presence of bias of such diffusion processes in each of its applications. In this work, we characterised some common null models behind the permutation analysis and the statistical properties of the diffusion scores. We benchmarked seven diffusion scores on three case studies: synthetic signals on a yeast interactome, simulated differential gene expression on a protein-protein interaction network and prospective gene set prediction on another interaction network. For clarity, all the datasets were based on binary labels, but we also present theoretical results for quantitative labels.

**Results:** Diffusion scores starting from binary labels were affected by the label codification, and exhibited a problem-dependent topological bias that could be removed by the statistical normalisation. Parametric and non-parametric normalisation addressed both points by being codification-independent and by equalising the bias. We identified and quantified two sources of bias -mean value and variance- that yielded performance differences when normalising the scores. We provided closed formulae for both and showed how the null covariance is related to the spectral properties of the graph. Despite none of the proposed scores systematically outperformed the others, normalisation was preferred when the sought positive labels were not aligned with the bias. We conclude that the decision on bias removal should be problem and data-driven, i.e. based on a quantitative analysis of the bias and its relation to the positive entities.

**Availability:** The code is publicly available at https://github.com/b2slab/diffuBench

## 1 Introduction

The guilt by association principle states that two proteins that interact with one another are prone to participate in the same, or related, cellular functions (Oliver, 2000). This cornerstone fact has motivated the exploration of network algorithms on interaction networks for protein function prediction (Sharan *et al.*, 2007). Network analysis has further proven its usefulness in other computational biology problems, such as prioritising candidate disease genes (Barabási *et al.*, 2011), finding modular structures (Mitra *et al.*, 2013) and modelling organisms (Aderem, 2005).

Network propagation is a fundamental formalism to leverage network data in computational biology. Its theoretical basis revolves around graph spectral theory, graph kernels and random walks (Smola and Kondor, 2003). The central concept is that nodes carry abstract labels that, following the guilt by association principle, are propagated to the neighbouring nodes (Zoidi *et al.*, 2015). Unlabelled nodes can therefore be inferred a label based on the available data of their neighbours. Label propagation can be defined in several ways, such as the heat diffusion, the electrical model or random walks with restarts (RWR), some of which lead to equivalent formulations (Cowen *et al.*, 2017).

One of the most common diffusion formulations relies on the regularised Laplacian graph kernel (Smola and Kondor, 2003) – examples are provided throughout this paragraph. HotNet (Vandin *et al.*, 2010) is a tool for finding modules with a statistically high number of mutated genes in cancer, after propagating the labels of mutated genes. The authors in (Bersanelli *et al.*, 2016) have found relevant modules from gene expression and mutation data, based on a diffusion process followed by an automatic subgraph mining. GeneMANIA (Mostafavi *et al.*, 2008) is a web server that predicts gene function by optimising a combination of knowledge networks and running a diffusion process on the resulting network. TieDIE (Paull *et al.*, 2013) defines two diffusion processes in order to connect two sets of genes, applied to link perturbation in the genome with changes in the transcriptome. More generally, the predictive power of label propagation using graph kernels has been benchmarked in gene-disease association (Valentini *et al.*, 2014; Lee *et al.*, 2011; Guala and Sonnhammer, 2017).

Some studies have pointed out biases in diffusion scores and explored the effect of their removal. The authors of DADA (Erten *et al.*, 2011) have found that prioritisation using RWR favours highly connected genes and suggest several normalisation strategies. One of them computes a z-score that adjusts for the mean value and standard deviation estimated from propagation scores from random degree-preserving inputs. Another possibility is to normalise diffusion scores into empirical p-values, as used in the diffusion of *t*-statistics derived from gene expression (Cun and Fröhlich, 2013). The aim was to quantify robust biomarkers, whose diffusion score is unlikely to arise from a permuted input. In the discovery of enriched modules (Bersanelli *et al.*, 2016), the effect of the topology has been mitigated by combining diffusion scores with their empirical p-values. Similarly, exact z-scores and empirical p-values have been used for pathway analysis of metabolomics data (Picart-Armada *et al.*, 2017b). A recent study (Biran *et al.*, 2019) has normalised RWR into an empirical p-value, obtained from edge rewiring. Specifically, random degree-preserving networks have been built to re-run the propagation and draw values from the null distributions of scores. Another recent manuscript (Hill *et al.*, 2019) highlights biases in certain network propagation algorithms, related to the node degree.

Overall, a variety of measures to address the bias have emerged, but a systematic quantification and evaluation of the biases is missing. The normalisation can potentially backfire, for instance by missing highly connected nodes that are associated with the property under study (Erten *et al.*, 2011). The goal of this manuscript is to provide a quantitative way to assess the presence of the bias and its alignment with the node labels, in order to understand the impact and adequateness of the normalisation.

## 2 Approach

Here, we address basic statistical properties of the normalisation of single-network diffusion scores to remove topology-related biases. We define and quantify two sources of bias. Both are derived from a statistical standpoint, based on the exact means and variances of the null distributions of the diffusion scores under input permutation. Differences in mean values between nodes should be the first indicator of systematic advantages: nodes with highest means will often be prioritised over those with lowest means. In their absence, differences in variances should be examined instead, as nodes with highest spread can be more likely to reach extreme scores. We compare classical and normalised propagation, as implemented in diffuStats (Picart-Armada *et al.*, 2017a), in data with and without bias. The main results are derived for the commonly used regularised Laplacian kernel, although most of them apply to other graph kernels and, to a lesser extent, to random walks with restarts. Special emphasis is placed on identifying scenarios under which normalisation is beneficial or detrimental and on understanding the underlying reasons why.

## 3 Methods

### Diffusion scores

We include seven diffusion scores that are part of the diffuStats package (Picart-Armada *et al.*, 2017a): 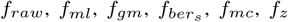 and 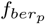. These scores are variations of the original diffusion model with a regularised unnormalised Laplacian kernel (Smola and Kondor, 2003). Labelled nodes are referred to as positives if they have the property of interest, and negatives otherwise.

#### Unnormalised scores

The starting point is the *f*_*raw*_ score, which requires a graph kernel *K* (Smola and Kondor, 2003) and input vector *y*_*raw*_ and is computed as:

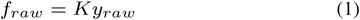

This work focuses on the unnormalised, regularised Laplacian kernel for *K*, for being a widespread choice in the computational biology literature (electrical model, heat or fluid propagation). The values in *y*_*raw*_ reflect the weights of each type of node: 1 for positives and 0 for negative and unlabelled entities.

*f*_*ml*_ and *f*_*gm*_ differ from *f*_*raw*_ by setting a weight of −1 on negative nodes. *f*_*gm*_ also weighs unlabelled nodes with a bias term adapted from GeneMANIA (not to be confused with the diffusion bias). On the other hand, 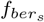 measures the relative change between *f*_*raw*_ and *y*_*raw*_, with a moderating parameter *ϵ*:

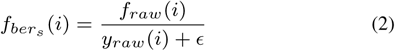

#### Normalised scores

Normalised scores attempt to equalise nodes that systematically show low or high scores, regardless of the input and due to the specific topology of the network. The lynchpin of normalisation is the null distribution of the diffusion scores under a random permutation *π* of the labelled nodes. The null scores arise from applying *f*_*raw*_ to a randomised input *X*_*y*_ = *π*(*y*_*raw*_) and comparing, for the *i*-th node, *f*_*raw*_ (*i*) to its null distribution *X*_*f*_ (*i*), where *X*_*f*_ = *KX*_*y*_. An empirical p-value can be computed throught Monte Carlo trials for the *i*-th node on *N* trials:

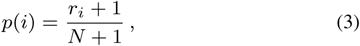

where *r*_*i*_ is the number of randomised trials having an equal or higher diffusion score in node *i*. In order to assign high scores to relevant nodes, the score is defined as *f*_*mc*_(*i*) = 1 − *p*(*i*). We also include a parametric alternative to *f*_*mc*_ by computing z-scores for each node *i*:

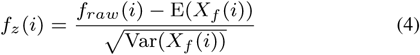

The expected value and variance of the null distributions are analytically determined (see Supplement 1). Thus, *f*_*z*_ has a computational advantage over Monte Carlo trials.

Finally, a hybrid combining an unnormalised and a normalised score is provided, inspired by how (Bersanelli *et al.*, 2016) moderated the effect of hubs: 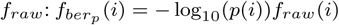.

#### Metrics and baselines

Two baseline methods were used. First `pagerank`, regarded as an input-naïve centrality measure (default damping factor of 0.85), to measure the predictive power of a basic network property. Second, a `random` predictor, to set an absolute baseline. Performances were quantified with two metrics: the area under the Receiver Operating Characteristic curve (AUROC) and the area under the Precision-Recall curve (AUPRC), as implemented in the precrec package (Saito and Rehmsmeier, 2017). For clarity, the ranking (ordering) of the nodes for any given score and instance was normalised to lie in [0, 1] by dividing it by the number of ranked nodes, so that top suggestions corresponded to ranks close to 0.

#### Bias quantification

The reference expected value of the *i*-th node 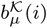 (eq. 7) was defined as proportional to the expected value of its null distribution *X*_*f*_ (*i*) (eq. 5). Reference expected values that vary across nodes can indicate systematic differences in the diffusion scores of such nodes.

In the absence of differences in the reference expected value, variance-related bias was analysed instead. The reference variance of the *i*-th gene 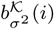 (eq. 8) was defined as, up to an additive constant, the base 10 logarithm of the variance of *X*_*f*_ (*i*), straightforward to obtain from the covariance matrix (eq. 6). The rationale is that the scores of nodes with varying dispersion measures should not be compared directly.

#### Performance explanatory models

Explanatory models have found use in the formal description of differences in performance as a function of design factors (Lopez-del Rio *et al.*, 2019; Picart-Armada *et al.*, 2019). Following (Picart-Armada *et al.*, 2019), the trends in AUROC and AUPRC were described through logistic-like quasibinomial models with a logit link function, as a generalisation of logistic models to prevent over and under-dispersion issues.

Table 1 presents the main model for each case study. The categorical regressors were: `method`, `metric` (AUROC or AUPRC), `biased` (refers to the signal, true or false), `strat` (labelled, unlabelled or overall), `array` (ALL or Lym), and the parameters `k`, `r` and `p_max` for the second case study. `path_var_ref` was quantitative, equal to the reference pathway variance 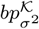 (eq. 9). The responses were either AUROC, AUPRC, or both mixed, the latter denoted by `Performance`.

**Table 1.** Case studies for characterising biases and benchmarking diffusion scores. Interactions in explanatory models are denoted by a colon.

| Case | Network | Positive nodes | Signal | Bias type | Purpose | Explanatory model for hypothesis testing |
| --- | --- | --- | --- | --- | --- | --- |
| (1) | Yeast | Synthetic | Synthetic, bias-based | Mean value | Proof of concept | Performance ∼ method + method : biased + metric |
| (2) | HPRD | KEGG pathways | Pathway sub-sampling | Mean value | Background influence in bias | AUPRC ∼ method + method : strat + array + $k + r + p_{max}$ |
| (3) | BioGRID | KEGG pathways | Prospective pathway prediction | Variance | Bias in a common scenario | AUROC ∼ method + method : path_var_ref |

## Materials

The evaluation of the diffusion scores was performed on three datasets of different nature, as described in Table 1: (1) synthetic signals on a yeast interactome, (2) pathway-based synthetic signals on a human network and (3) real signals on another human network.

### Networks

#### Yeast network

A small yeast network was used to demonstrate the casuistic of diffusion scores properties. Medium and high confidence interactions from several sources were provided by the original study (Von Mering *et al.*, 2002), as found in the igraphdata R package (Csardi, 2015). It contains 2,617 proteins and 11,855 unweighted edges, but we worked only with its largest connected component (2,375 proteins, 11,693 edges).

#### HPRD network

The diffuse large B-cell lymphoma study, available in the R package DLBCL (Dittrich and Beisser, 2010), contains a differential expression dataset accompanied by a human interactome network extracted from the Human Protein Reference Database, HPRD (Mishra *et al.*, 2006). The original network encompasses 9,385 proteins with 36,504 interactions, whose largest connected component (8,989 nodes, 34,325 interactions) was extracted to compute the diffusion scores.

We derived two gene backgrounds based on expression arrays. The first background (Lym) was taken from the expression data from 2,557 genes (2,482 in the network) in the lymphoma study (Rosenwald *et al.*, 2002). The second background (ALL) was based on the acute lymphocytic leukemia array (Chiaretti *et al.*, 2004), available in the ALL R package (Li, 2009), encompassing 6,133 genes (5,921 in the network).

#### BioGRID network

The Biological General Repository for Interaction Datasets (BioGRID) (Chatr-aryamontri *et al.*, 2017) is a public database with curated genetic and protein interaction from Homo sapiens and other organisms. BioGRID was retrieved in January 2017, but only keeping interactions dating from 2010 or older. The interactions were weighted according to (Cao *et al.*, 2014), under the assumptions that more publications about an interaction boost its confidence and that low-throughput technologies are more reliable that high-throughput ones. The network encompassed 11,394 nodes and 67,573 edges and was connected.

### Datasets

#### Synthetic bias-based dataset

100 biased and 100 unbiased instances of positive, negative and unlabelled nodes were generated in dataset (1) from table 1, by sampling positive nodes with probabilities proportional to biased and unbiased scores. By construction, the frequencies of the positives drawn for biased signals were positively correlated with the reference expected value, whereas those of the unbiased signals were uncorrelated with it.

Nodes were partitioned into three equally sized pools, from which positive nodes were drawn: (a) labelled nodes that were fed to the diffusion methods, (b) target nodes, the ones to be ranked and whose ground truth was known, and (c) filler nodes that were neither target nor labelled.

For each instance, a fixed fraction of labelled nodes *x*_*e*_ were uniformly sampled as positives, the rest of labelled nodes were deemed negatives and the target and filler nodes were left unlabelled. This input served two purposes: generate the ground truth in target nodes, and be the input for all the diffusion scores.

To generate the ground truth in target nodes of biased signals, the `raw` diffusion scores were computed from the input above. A fixed fraction of target nodes *x*_*s*_ was sampled with probabilities proportional to their raw scores, i.e. *p*(*i*) ∝ *f*_*raw*_ (*i*), to become positives. The remaining target nodes would remain negatives, completing the ground truth. The regularised unnormalised Laplacian kernel is endorsed by physical models that ensure *f*_*raw*_ (*i*) > 0 provided that inputs have one or more positives and the graph is connected. Analogously, unbiased signals were generated by sampling a fraction of target nodes *x*_*s*_, but with probabilities roughly proportional to the unbiased diffusion scores `mc`: 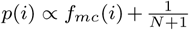. By definition, the frequency of appearance of the target nodes was independent of the bias, and the small offset ensured *p*(*i*) > 0.

In both cases, after sampling the ground truth, the same input was used again for all the diffusion scores, in order to rank the target nodes and compute the corresponding AUROC and AUPRC.

#### Pathway sub-sampling dataset

Synthetic gene expression statistics were generated, based on pathways in the Kyoto Encyclopedia of Genes and Genomes (KEGG) (Kanehisa *et al.*, 2017), and on two array-based gene backgrounds described within the HPRD network. Genes outside the background were hidden (unlabelled), and genes inside were given p-values for differential expression.

Each signal derived from *k* random KEGG pathways. The pathways were assumed to be affected as a whole, but only a sampled portion of *r* genes showed differential expression patterns. The p-values of the differential expressed genes were uniformly sampled from [0, *p*_*max*_], whereas the rest of genes were uniform in [0, 1], following a previously study (Rajagopalan and Agarwal, 2004).

For both expression arrays, genes with an FDR < 10% within their background were used as positives, the remaining background genes as negatives and the hidden nodes were deemed unlabelled. Notice that, by definition, this procedure generated false positives and false negatives among the input genes.

The target genes were those belonging to the *k* affected pathways, including those with no apparent differential expression and those among the unlabelled nodes. Methods were compared using the AUROC and AUPRC, computed separately on labelled, unlabelled genes, and overall, on a grid of parameters: *k* ∈ {1, 3, 5}, *r* ∈ {0.3, 0.5, 0.7} and *p*_*max*_ ∈ {10^−2^, 10^−3^, 10^−4^}. For each combination of parameters, *N* = 50 instances were simulated.

#### Prospective pathway dataset

The input lists consisted of the genes in 139 KEGG pathways from 14th March, 2011. The target genes were the newly added genes in the same KEGG pathways in 18th August, 2018 release. The 139 pathways had new genes in the latter release after mapping to the network.

AUROC and AUPRC were computed on each pathway, always excluding the input positive genes. The bias was examined at the pathway level, assessing whether the properties of their new genes differed from those of the rest of network genes. It was defined as the median reference variance of its new genes minus the median reference variance of all the genes besides old and new pathway genes (eq. 9).

## 4 Results

### Properties of diffusion scores

Some of the diffusion scores are equivalent in certain scenarios. In the absence of unlabelled nodes and using kernels based on the unnormalised graph Laplacian, *f*_*raw*_, *f*_*ml*_ and *f*_*gm*_ lead to an identical node prioritisation. More generally, the results using only two classes (and therefore two real values *y*^+^ > *y*^−^ as weights) always lead to the same ranking as *f*_*raw*_. An analogous result holds for the weights of the positives and the unlabelled, *y*^+^ > *y*^*u*^, in the absence of negative nodes.

The normalised scores *f*_*mc*_ and *f*_*z*_ are invariant to changes in the weights of the positive and negative examples, regardless of the presence of unlabelled nodes and the graph kernel. This property simplifies the diffusion setup and leads to weight-independent results. Along with eqs. 5 and 6, this holds even if the matrix *K* in eq. 1 is not a kernel, like the random walk similarity matrices in (Cowen *et al.*, 2017).

We also provide the closed form of the null expected value and covariance matrix of the `raw` scores, governed by the identifiers of the *n*_*l*_ labelled nodes (out of *n*). If 𝒦 contains only their corresponding columns from *K*, and 𝒴 is the input vector *y*_*raw*_ restricted to them, then:

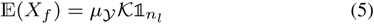

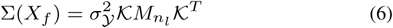

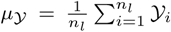 and 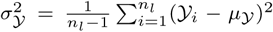 are the mean and variance of the labels. 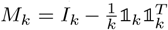, being *I*_*k*_ the *k* × *k* identity matrix and 𝟙_*k*_ the column vector with *k* ones.

If a graph kernel based on the unnormalised Laplacian is used, the covariance of the null distribution (eq. 6) is closely related to the spectral properties of the labelled nodes. In particular, in the absence of unlabelled nodes, the leading eigenvector of the null covariance is, up to a sign change, the Fiedler-vector, commonly used for graph clustering (Smola and Kondor, 2003). The statistical normalisation is therefore endowed with a topological basis. This sheds light on prior empirical observations that, even when the bias can relate to the node degree, there must be further topological factors involved (Hill *et al.*, 2019).

Because *µ* _𝒴_ and 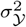 are multiplicative constants and inherent to the labels, the topology-related mean value and variance references of the *i*-th node are defined as follows. We assume *n*_*l*_ ≥ 2 because if *n*_*l*_ ∈ {0, 1} there is nothing to permute.

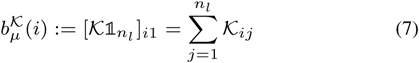

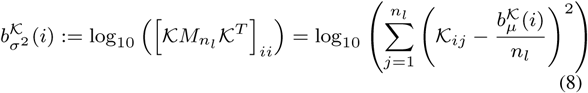

Eq. 5 implies that there are two scenarios free of the expected value bias: *µ*_𝒴_ = 0 (centered input), or *n*_*l*_ = *n* and a kernel *K* based on the unnormalised Laplacian, rendering 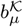 constant (see Supplement 1). The *i*-th null variance (eq. 6) can be exactly zero, either because 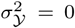 (constant input), or because the topology forces 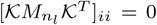. In practice, the latter is expected to happen in small connected components without any labelled nodes. Both cases render the *i*-th score constant, therefore lacking interest, and leave *f*_*z*_ undefined.

In the retrospective dataset, the reference of a given pathway *P*, conceived to summarise its properties into a single number, was defined by subtracting the median reference of its new genes, new(*P*) to that of the genes that never belonged to it, others(*P*):

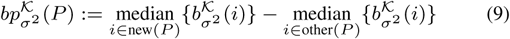

The mathematical proofs of the properties and illustrative examples can be found in Supplement 1.

### Synthetic signals in yeast

#### Bias in diffusion scores

Supported by eq. 5, the presence of unlabelled nodes originated different expected values among the nodes. We hypothesised that *f*_*raw*_ would be biased to favour nodes with high 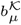, whereas *f*_*mc*_ and *f*_*z*_ would prioritise in a more unbiased manner. Figure 1A confirms both trends. The data imbalance (negatives outnumbered positives in the input) had the opposite biasing effect on *f*_*ml*_, favouring nodes with low 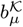.

**Fig. 1.**
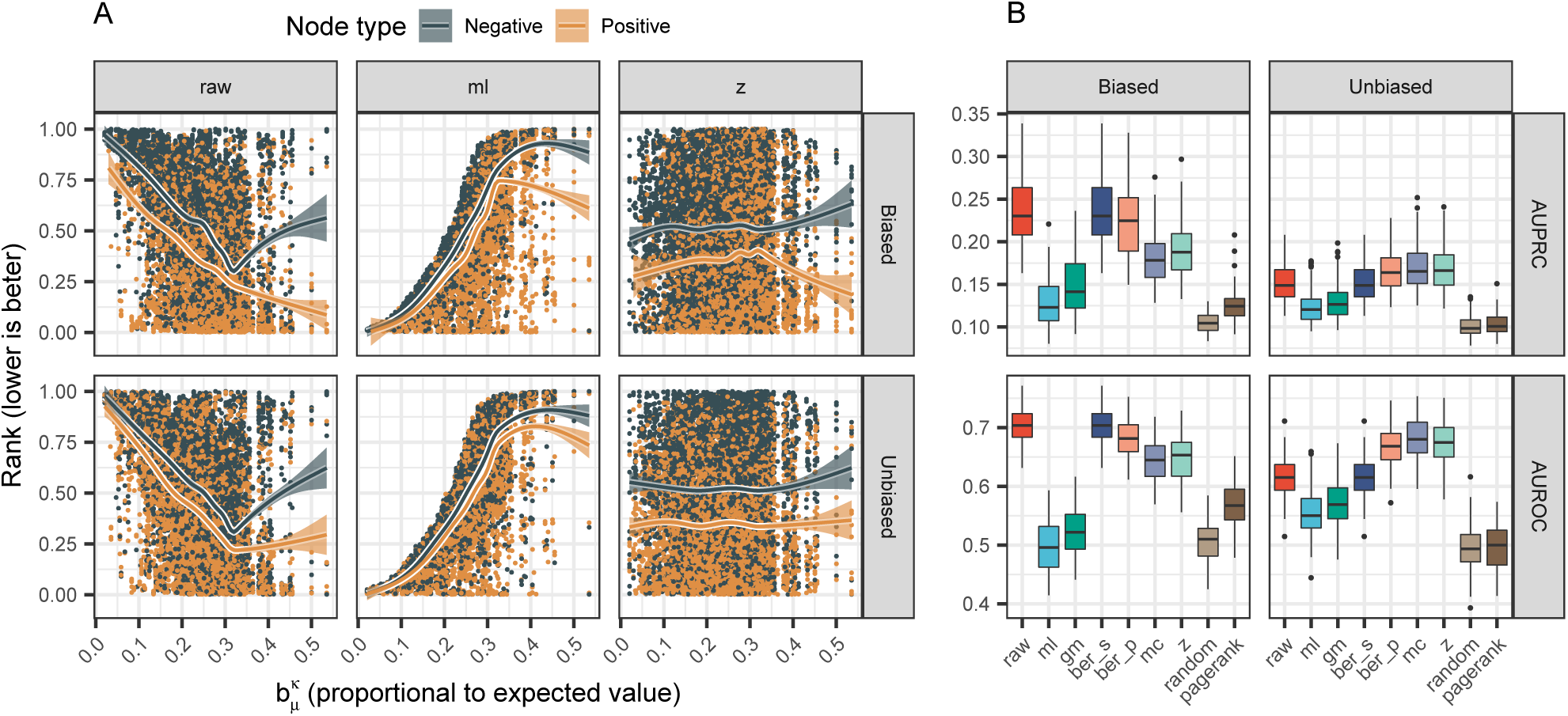
Analysis of biased and unbiased synthetic signals on the yeast network. Nodes showed a mean value-related bias, see Supplement 2. (A) Effects of the mean value bias in on the average node ranking, under biased and unbiased signals. Lines correspond to Generalised Additive Models with *y* ∼ *s*(*x, bs* = “*cs*”) and 0.95 confidence intervals. `raw` and `ml` tended to find positives with high and low 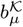, respectively. `z` found positives in a more uniform manner. (B) Performance in terms of AUROC and AUPRC. `raw` was better suited for biased signals, for which the `pagerank` baseline also outperformed a random predictor. Conversely, `z` worked best on unbiased signals.

#### Performance

In biased signals, target nodes with higher 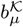 were sampled as positives more often (see Supplement 2), which (i) benefited the unnormalised scores `raw` over `z`, and (ii) endowed the `pagerank` baseline with predictive power. Unbiased signals led to a uniform density of positives across 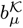, which (iii) was better handled by `z` than by `raw` (figure 1B). Claims (i), (ii) and (iii) were statistically significant for AUROC and AUPRC (Tukey’s method, FDR < 10^−10^ in all cases, see Supplement 2). Also, 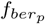 was a good compromise between `raw` and `z`.

Based on these results, we suggest a systematic criterion to choose whether to normalise in the general case, by assessing (1) the presence of the expected value-related bias by checking if 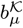 is constant among the nodes to be prioritised, and (2) the expected or hypothetical dependence between 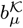 and the labels to be predicted. In this proof of concept, differences in 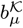 bias were present and normalisation was discouraged when 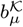 was aligned with the positives. If 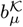 is constant, 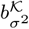 should be examined instead, see the retrospective pathway dataset.

### Simulated differential expression

#### Bias in diffusion scores

Analogously to the yeast dataset, the presence of unlabelled nodes led to differences in 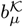 among nodes, see figure 2A. We hypothesised that the main source of bias would arise from such heterogeneity, i.e. that unnormalised scores would be prone to find positives among highest expected values. In both arrays, the nodes belonging to one or more pathways had, compared with nodes outside, (i) larger 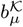 within the unlabelled genes, but (ii) lower 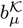 within the labelled nodes. Overall, (iii) labelled genes showed larger 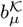 than unlabelled genes. Figure 2A portrays the claims (i), (ii) and (iii) in both arrays – the six statements were significant with *p* < 10^−16^, two-sided Wilcoxon test (see Supplement 3).

**Fig. 2.**
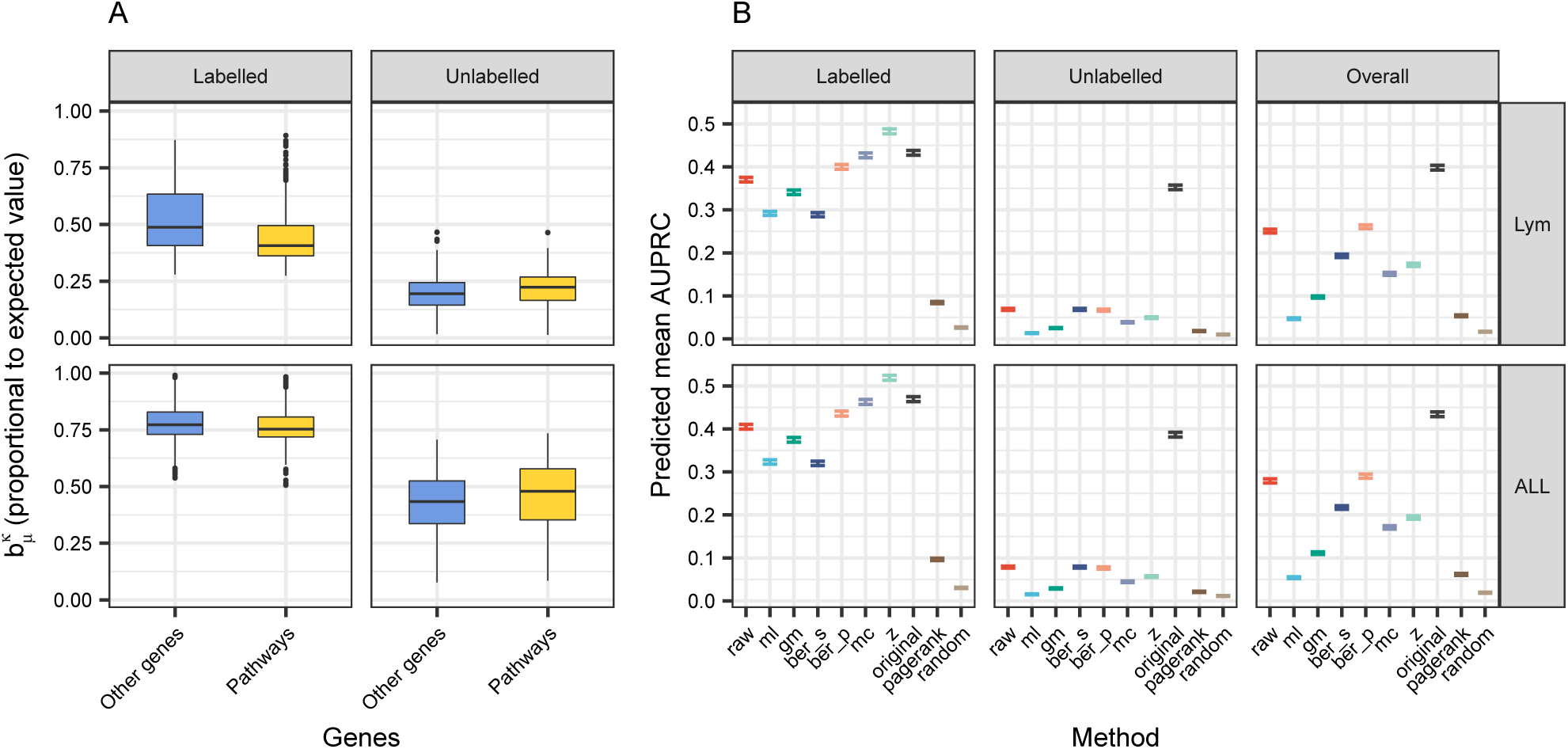
Performance in the DLBCL dataset. (A) Expected value-related bias. Within the labelled genes of both arrays, those in pathways had lower 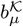 that those outside. Within the unlabelled genes, this tendency was inverted. Overall, labelled genes had higher 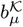 than unlabelled genes. (B) Predicted AUPRC (0.95 confidence interval) using the explanatory model in Table 1 and Supplement 3. Besides diffusion scores, three baselines were included: `original` (ranking by the p-values), `pagerank` and `random`. In both arrays (ALL and Lym), `raw` outperformed `z` in unlabelled nodes and overall, while `z` was preferable in the labelled genes.

#### Performance

The performance, as predicted by the explanatory models, was influenced by the background used to compute the metrics, especially for AUPRC. Taking as reference *f*_*raw*_ and *f*_*z*_, `raw` performed best in the unlabelled background and overall whereas `z` was preferable in the labelled background (figure 2B). The three claims were significant in both arrays (Tukey’s method, *p* < 10^−10^, see Supplement 3).

Differences in performance were consistent with the expected value-related bias: potential positives suffered from lower 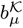 in the labelled genes and benefited from greater 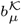 in the unlabelled part. In views of this, the natural choices were `z` and `raw`, respectively.

To understand why `raw` outperformed `z` in overall performance, note how by hypothesis the top candidates from `raw` should come from the labelled genes due to their high 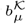 against the unlabelled genes, whilst `z` should equalise predictions from both backgrounds. Predictions from the labelled part were more reliable owing to the presence of prior data on the genes (figure 2B). `z` equalised both backgrounds, shuffling reliable and unreliable predictions, and undermined overall performance.

Finally, an indirect assessment of the bias (PageRank centrality) fell short to explain performance differences in (i) and suggested that biased scores were preferrable in the three cases, see Supplement 3. This highlights the importance of using a precise quantification of the bias.

### Prospective pathway prediction

#### Bias in diffusion scores

Here, 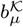 was constant among all the nodes, as a consequence of using the unnormalised Laplacian without unlabelled nodes (see Supplement 1). Differences still existed in terms of 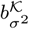 (figure 3A), implying that the normalisation would make a difference.

**Fig. 3.**
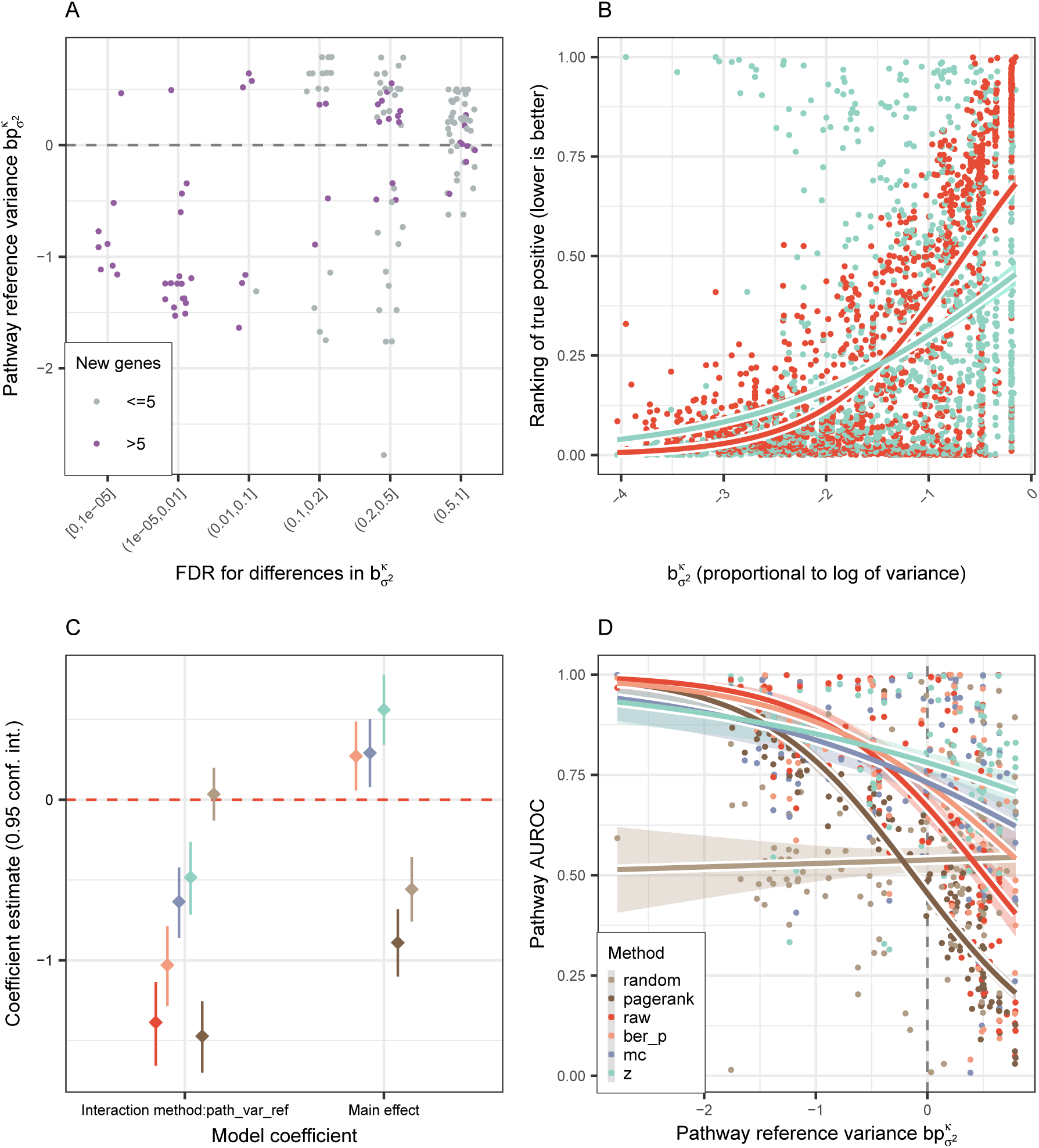
Analysis of the prospective dataset. (A) Pathway-wise comparison of new genes against the remaining genes outside the pathway, in terms of 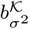. Several pathways showed significant differences in both directions (two-sided Wilcoxon test). The *x* axis was jittered for clarity. (B) Ranking of the positives using `raw` and `z`. Each data point is the relative ranking of a positive gene in one of the pathways, i.e. before computing pathway-level metrics. Lines correspond to a quasi-logistic fit with a 0.95 confidence interval. `raw` scores were more sensitive at low standard deviations, whereas `z` stood more uniform. (C) Coefficients of the model AUROC ∼ method + method : path_var_ref with a 0.95 confidence interval, where the interaction term involved the variance bias. The main effect of `raw` was not depicted because it was the reference level of `method`. (D) Predicted AUROC across all the pathways, as a function of the bias. `z` was less sensitive to the bias, due to its interaction term in (C) being closer to 0. Lines correspond to a quasi-logistic fit with a 0.95 confidence interval.

However, the interpretation of the normalisation impact was not as straightforward as for the expected value bias. With the paradigm of the z-scores `z`, deviations from the expected value exacerbate under small variances and shrink under large variances. Notice how this does not imply the natural hypothesis that nodes with larger variances (resp. smaller) must drop (resp. rise) in the ranking, because ranking modifications take place around the mean. Figure 3B reflects how `z` actually recovered more high-variance positive nodes than `raw`.

Similarly to prior observations from figure 1A, the normalised scores tended to find the positives in a less biased manner. Positive nodes with a high variance were rarely found by `raw`, whereas `z` distributed them more evenly along the ranking (figure 3B). This improvement came at the cost of missing positives with lower variances.

### Performance

The properties of the diffusion scores helped simplify this case study, as *f*_*ml*_, *f*_*gm*_ and *f*_*ber*_*s*_ were left out for being redundant with *f*_*raw*_. *f*_*ml*_ and *f*_*gm*_ for using the unnormalised Laplacian without unlabelled nodes, and 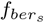 because the genes to be prioritised were always labelled as negative in the input (see corollary 1 and proposition 3 in Supplement 1).

The prospective prediction of pathway genes was a challenging task, given the low predicted AUPRCs for all the methods (see Supplement 4). On the other hand, AUROC conveyed a richer view of the differences between methods. The explanatory model (figures 3C, 3D) showed that unnormalised scores were more affected by the presence of bias, reflected in the larger magnitude of their interaction terms (−1.387 for `raw` against −0.484 for `z`, *p* < 10^−4^, Tukey’s method). Overall, the casuistic among the bias of new pathway genes favoured `z` over `raw` (FDR = 5.39 · 10^−9^, two-sided paired Wilcoxon test). This conclusion did not apply to early retrieval, as it could not be proven for AUPRC (FDR = 0.701).

The negative sign of the interaction terms was also insightful: all the proper methods encountered more difficulties in finding loosely connected genes. This was expected, since there is less network data involving such genes, translating into unreliable predictions.

## Conclusion

In this study, we ratified that diffusion scores are biased due to the graph topology. We introduced two direct quantifications of the bias, in terms of the expected value and variance of the null distribution of the diffusion scores under input permutation. We analysed the benefits and pitfalls of using unbiased, statistically normalised scores and discussed several choices of the label weights when defining the diffusion process.

We proved equivalences between scores under certain conditions, helping simplify the setup of the diffusion, and discovered that normalised alternatives are invariant under label weights changes. We found an explicit link between principal directions of the null covariance and the spectral features of the network.

We applied the diffusion-based prioritisation on three scenarios: two with a mean value-related bias and one with a variance-related bias. Class imbalance and node topology had an impact in unnormalised scores, whereas normalised scores were more robust to both phenomena given their weight-independent definition. The parametric normalisation requires no permutations compared to Monte Carlo trials and performed equally or better, providing a convenient way to normalise. While mean value bias was straightforward to characterise, variance bias was less intuitive albeit of noticeable impact. In general terms, the statistical normalisation is advised if the positives are not aligned with the bias, and discouraged otherwise. The statistical background, i.e. which nodes are permuted, is a key piece that should be clearly stated in every application. Bias assessment should be carried through its direct quantification instead of indirect indicators, which can be misleading.

We conclude that the statistical normalisation can be beneficial or detrimental, and the decision should follow from the dependence between the node bias and the hypothetical or desired properties of the new positives. Topology-related bias can manifest in different ways (mean value- or variance-related bias) and each instance should be properly characterised.

## Acknowledgements

SP thanks Guillem Belda-Ferrín for reviewing the mathematical proofs. SP thanks Imanol Morata Martínez and Camellia Sarkar for fruitful discussions and useful suggestions.

