## Supplement 1 for "The effect of statistical normalisation on network propagation scores"

November 19, 2018

### Introduction

This document outlines and proves several properties of the diffusion scores discussed in the main body. We mainly derive equivalences between scores and properties of the statistical normalisations and their null distributions.

#### Notation

Tables 1 and 2 contain an overview of the notation used to formulate and prove the properties.

Table 1: Notation of matrices, vectors and scalars.

| Notation | Data type | Description |
|---|---|---|
| $n$ | Scalar | Number of nodes in the graph |
| $n_+$ | Scalar | Number of positive nodes |
| $n_-$ | Scalar | Number of negative nodes |
| $n_l$ | Scalar | Number of labelled nodes, $n_l = n_+ + n_-$ |
| $n_u$ | Scalar | Number of unlabelled nodes, $n_u = n - n_l$ |
| $L$ | Matrix | Unnormalised graph Laplacian $n \times n$ matrix |
| $v_i$ | Column matrix | $i$-th eigenvector of $L$ |
| $v_i^T$ | Row matrix | $i$-th eigenvector of $L$ |
| $\lambda_i$ | Scalar | $i$-th eigenvalue of $L$ |
| $K$ | Matrix | Graph kernel $n \times n$ matrix from [2] |
| $\mathcal{K}$ | Matrix | $n \times n_l$ sub-matrix from K (columns indexed by labelled nodes) |
| $K_{ij}$ | Scalar | Entry $(i, j)$ from $K$ |
| $K_{i*}$ | Row matrix | $i$-th row of kernel matrix K |
| $K_{*j}$ | Column matrix | $j$-th column of kernel matrix K |
| $y_{raw}$ | Column matrix | Vector in $\mathbb{R}^n$ with the input scores to `raw` diffusion scores |
| $y_{raw}(i)$ | Scalar | Input score of $i$-th node in the `raw` method |
| $\mathcal{Y}_{raw}$ | Column matrix | Vector in $\mathbb{R}^{n_l}$, $y_{raw}$ restricted to labelled nodes only |
| $y^+_{raw}$ | Scalar | Weight of the positive class for the `raw` scores |
| $y^-_{raw}$ | Scalar | Weight of the negative class for the `raw` scores |
| $y^u_{raw}$ | Scalar | Weight of the unlabelled class for the `raw` scores |
| $f_{raw}$ | Column matrix | Vector with the `raw` scores |
| $f_{raw}(i)$ | Scalar | `raw` score of $i$-th node |
| $\mathbb{1}_k$ | Column matrix | Vector whose $k$ entries are 1 |
| $I_k$ | Matrix | $k \times k$ identity matrix |

The graph Laplacian $L$ is a real $n \times n$ matrix defined as $L := D - W$, where $D = D_{ii}$ is the (diagonal) degree matrix and $W = W_{ij}$ the adjacency matrix. This definition assumes an undirected graph, either unweighted or with weights $W_{ij} \in [0, \infty)$ [2].

Table 2: Notation of functions and operators.

| Notation | Function/operator | Notes |
|---|---|---|
| $r(\lambda)$ | Regularisation function [2] | $\lambda \geq 0$, $r(\lambda) \geq 0$ monotonically increasing |
| $\frac{1}{r(\lambda)}$ | Inverse of $r(\lambda)$ | By convention, $0^{-1} \equiv 0$, see [2] |
| $\mathbb{E}(X)$ | Expected value | $X$ random vector in $\mathbb{R}^k$, $\mathbb{E}(X) \in \mathbb{R}^k$, both column matrices |
| $\Sigma(X)$ | Covariance | $X$ as above, $\Sigma(X) \in \mathbb{R}^{k \times k}$ symmetric square matrix |

Importantly, some of the present proofs use a sub-matrix $\mathcal{K}$ of the whole graph kernel $K$ [2] (we focus on the finite dimension case). $\mathcal{K}$ contains the **rows corresponding to all the graph nodes and the columns corresponding to labelled nodes**. In the absence of unlabelled nodes, $\mathcal{K} = K$ will be a square matrix, i.e. the whole kernel matrix – see for instance proposition 1. Otherwise $\mathcal{K}$ will be a rectangular sub-matrix of it, containing all the original rows but only some of the columns; the latter is not a kernel matrix properly speaking. This simplifies the notation because (i) only one score in our study, `gm`, actually places non-null weights on the unlabelled class, and (ii) the normalised scores permute only the labelled nodes.

Likewise, the input vector $\mathcal{Y}$ represents $y$ indexed by the labelled entries. For instance, the vector of input labels for the `raw` scores $\mathcal{Y}_{raw}$ contains only the nodes under the positive ($y_{raw}(i) = y_{raw}^{+} = 1$) and negative classes ($y_{raw}(i) = y_{raw}^{-} = 0$), but not the unlabelled nodes.

Therefore, the matrix-vector product $f_{raw} = \mathcal{K}\mathcal{Y}_{raw}$ is properly defined in terms of dimensionality. If $n$ is the number of nodes in the graph and $n_u$ the number of unlabelled nodes, then $f_{raw} \in \mathbb{R}^n$, $\mathcal{K} \in \mathbb{R}^{n \times (n-n_u)}$ and $\mathcal{Y}_{raw} \in \mathbb{R}^{n-n_u}$. This is equivalent to including the unlabelled with a weight of $y_{raw}^{u} = 0$; this is, $f_{raw} = K y_{raw}$, with $f_{raw} \in \mathbb{R}^n$, $K \in \mathbb{R}^{n \times n}$ and $y_{raw} \in \mathbb{R}^n$.

#### Definition of the scores

While the input can be quantitative, we focus on the case where the entities can only be positive, negative or unlabelled. The `raw` diffusion scores are defined as $f_{raw} = K y_{raw} = \mathcal{K}\mathcal{Y}_{raw}$, according to [1]. $y_{raw}$ takes by default these weights: $y^{+} = 1$ for the positives, $y^{-} = y^{u} = 0$ for the negatives and unlabelled nodes. In general, a diffusion score $f$ can be computed with other weights as $f = Ky$, where $K$ is the graph kernel and $y$ the input coded by another choice of $y^{+}$, $y^{-}$ and $y^{u}$.

We use the diffuStats package [1], equipped with the possibilities that are studied in the main body. One of the aims of the following properties is to prove equivalences between label choices in certain conditions are met.

### Equivalences between scores

**Proposition 1.** *Consider* $f_{raw} = Ky_{raw}$ *of finite dimension in a scenario with no unlabelled nodes (*$K$ *square matrix), i.e. the label of the* $i$*-th node* $y_i = y^+_{raw} = 1$ *for the positives and* $y_i = y^-_{raw} = 0$ *for the negatives, and* $n_u = 0$*. Let* $f = Ky$ *be another score using* $y^+ > y^-$ *as new real numbered weights for positives and negatives. Then, if the kernel* $K$ *is a spectral transformation of the unnormalised graph Laplacian* $L$*, the result of ranking (prioritising) the nodes using* $f_{raw}$ *and using* $f$ *is identical.*

*Proof.* Let $f_{raw}(i_1) \geq f_{raw}(i_2) \geq \ldots \geq f_{raw}(i_n)$ be the ranking of the nodes using the $f_{raw}$ scores, i.e. their prioritisation through decreasing $f_{raw}$, being the top suggestions the highest scores. If we prove that $f_{raw}(i) \geq f_{raw}(j) \Leftrightarrow f(i) \geq f(j)$, then the ranking using the scores $f$ must be identical. Note that ties in $f_{raw}$ must happen in $f$ and vice versa, because $f_{raw}(i) = f_{raw}(j) \Leftrightarrow f_{raw}(i) \geq f_{raw}(j) \wedge f_{raw}(i) \leq f_{raw}(j) \Leftrightarrow f(i) \geq f(j) \wedge f(i) \leq f(j) \Leftrightarrow f(i) = f(j)$.

As $K$ is a spectral transformation of the unnormalised Laplacian $L$, it can be written in the following form, see [2]:

$$K = \sum_{j=1}^{n} \frac{1}{r(\lambda_j)} v_j v_j^T$$

Being $v_j$ the eigenvectors as column vectors and $\lambda_j$ the eigenvalues of $L$. The constant vector $v_1 = \frac{1}{\sqrt{n}} (1, \ldots, 1)^T$ is an eigenvector of $L$ of eigenvalue 0. Therefore, it is also an eigenvector of $K$ of eigenvalue $\frac{1}{r(\lambda_1)} = \frac{1}{r(0)} \in \mathbb{R}$ (by convention, $\frac{1}{0} \equiv 0$). Therefore, the $i$-th row of $K$, denoted $K_{i*}$, $1 \leq i \leq n$, has a constant sum, as $Kv_1 = \frac{1}{r(0)} v_1$, and because $v_1 = \frac{1}{\sqrt{n}} (1, \ldots, 1)^T$:

$$\sum_{j=1}^{n} K_{ij} = \frac{1}{r(0)}, 1 \leq i \leq n$$

On the other hand,

$$\begin{aligned} f_{raw}(i) \geq f_{raw}(j) &\Leftrightarrow K_{i*} y_{raw} \geq K_{j*} y_{raw} \\ &\Leftrightarrow (K_{i*} - K_{j*}) y_{raw} \geq 0 \end{aligned}$$

Note that if $\mathbb{1}_n$ is the column vector full of ones, then

$$y_{raw} = \frac{1}{y^+ - y^-} (y - y^- \mathbb{1}_n)$$

where $y^+ - y^- > 0$. Therefore,

$$
\begin{aligned}
(K_{i*} - K_{j*})y_{raw} \geq 0 &\Leftrightarrow (K_{i*} - K_{j*})\frac{1}{y^+ - y^-}(y - y^-\mathbb{1}_n) \geq 0 \\
&\Leftrightarrow (K_{i*} - K_{j*})(y - y^-\mathbb{1}_n) \geq 0 \\
&\Leftrightarrow (K_{i*} - K_{j*})y - y^-(K_{i*} - K_{j*})\mathbb{1}_n \geq 0 \\
&\Leftrightarrow (K_{i*} - K_{j*})y - y^-\left(\frac{1}{r(0)} - \frac{1}{r(0)}\right) \geq 0 \\
&\Leftrightarrow (K_{i*} - K_{j*})y \geq 0 \\
&\Leftrightarrow f(i) \geq f(j)
\end{aligned}
$$

□

Note how a more general version of this property can be proved in a very similar way for quantitative $y_{raw}$, so that transformations of the kind $y = \alpha y_{raw} + \beta\mathbb{1}_n$, with $\alpha, \beta \in \mathbb{R}, \alpha > 0$, lead to the same node ranking as $f_{raw}$.

**Corollary 1.** *The scores $f_{raw}$, $f_{ml}$ and $f_{gm}$ lead to the same node ranking (prioritisation) if there are no unlabelled nodes and the kernel $K$ is a spectral transformation of the unnormalised graph Laplacian $L$ of finite dimension.*

Note that this result does not hold in general if the kernel stems from the normalised Laplacian, or with the presence of unlabelled nodes besides positives and negatives (see counterexamples on figure 1). In both cases, the row sums are no longer constant and their respective sums do not cancel out.

The same property holds if instead of having only positives and negatives there are only positive and unlabelled nodes:

**Proposition 2.** *Consider $f_{raw} = Ky_{raw}$ of finite dimension in a scenario with no negative nodes, i.e. the label of the i-th node $y_i = y^+_{raw} = 1$ for the positives and $y_i = y^u_{raw} = 0$ for the unlabelled nodes, and $n_- = 0$. Let $f$ be another score using $y^+ > y^u$ as new real numbered weights for positives and unlabelled nodes. Then, if the kernel $K$ is a spectral transformation of the unnormalised graph Laplacian $L$, the result of ranking (prioritising) the nodes using $f_{raw}$ and using $f$ is identical.*

*Proof.* The proof is identical to proposition 1, but switching the roles of unlabelled and negative nodes. □

**Corollary 2.** *The scores $f_{raw}$, $f_{ml}$ and $f_{gm}$ lead to the same node ranking if there are no negative nodes and the kernel $K$ is a spectral transformation of the unnormalised graph Laplacian $L$ of finite dimension.*

**Proposition 3.** *The ranking using $f_{raw}$ and $f_{ber_s}$ is identical within the positive, the negative and the unlabelled nodes, provided that $\epsilon > 0$*

*Proof.* If the $i$-th node is a positive, then:

$$f_{ber_s}(i) = \frac{f_{raw}(i)}{y_{raw}(i) + \epsilon} = \frac{f_{raw}(i)}{1 + \epsilon}$$

Therefore, there is only a positive, multiplicative constant between $f_{raw}$ and $f_{ber_s}$.

If the $i$-th node is a negative or unlabelled, then:

$$f_{ber_s}(i) = \frac{f_{raw}(i)}{y_{raw}(i) + \epsilon} = \frac{f_{raw}(i)}{\epsilon}$$

Where $f_{raw}$ and $f_{ber_s}$ clearly lead to the same ranking (prioritisation) again. □

### Normalisations are invariant under label codification

**Proposition 4.** *In $f_z$, the choice of $y^+$ and $y^-$ is irrelevant. More generally, computing $\tilde{f}_z$ using $\tilde{\mathcal{Y}} = \alpha\mathcal{Y} + \beta\mathbb{1}_{n_l}$ instead of $\mathcal{Y}$, with $\alpha, \beta \in \mathbb{R}, \alpha > 0$, is equivalent to computing $f_z$.*

*Proof.* Using the same notation as in the first property, we start from the definition of the z-score from the main text, which normalises the $f_{raw}$ score by the mean and standard deviation of its distribution when the labelled nodes are permuted:

$$f_z(i) = \frac{f_{raw}(i) - \mathbb{E}(X_f(i))}{\sqrt{\mathrm{Var}(X_f(i))}} = \frac{\mathcal{K}_{i*}\mathcal{Y}_{raw} - \mathcal{K}_{i*}\mathbb{E}(X_\mathcal{Y})}{\sqrt{\mathcal{K}_{i*}\Sigma(X_\mathcal{Y})\mathcal{K}_{i*}^T}}$$

According to the main text, $X_\mathcal{Y}$ is a random permutation of the labelled nodes in the input $\mathcal{Y}_{raw}$, and $X_f = \mathcal{K}X_\mathcal{Y}$ is the random vector of null diffusion scores.

We prove that computing $f_z$ with an input vector $\mathcal{Y}$ is identical to doing the same with $\tilde{\mathcal{Y}} = \alpha\mathcal{Y} + \beta\mathbb{1}_{n_l}$. Let $X_\mathcal{Y}$ be a random permutation of the input $\mathcal{Y}$, treated as a random vector, and let $X_{\tilde{\mathcal{Y}}}$ be the same permutation applied to $\tilde{\mathcal{Y}}$. The raw diffusion score of the $i$-th node is $\mathcal{K}_{i*}\mathcal{Y}$, whereas its null distribution is the random variable $\mathcal{K}_{i*}X_\mathcal{Y}$. Analogously, using $\tilde{\mathcal{Y}}$, the score is $\mathcal{K}_{i*}\tilde{\mathcal{Y}}$ and the null distribution is $\mathcal{K}_{i*}X_{\tilde{\mathcal{Y}}}$. The idea is that subtracting the expected value cancels out the constant term $\beta$, whereas dividing by the standard deviation cancels out the multiplicative constant $\alpha$.

For any node $i$:

$$\begin{aligned}
\tilde{f}_z(i) &= \frac{\mathcal{K}_{i*}\tilde{\mathcal{Y}} - \mathcal{K}_{i*}\mathbb{E}(X_{\tilde{\mathcal{Y}}})}{\sqrt{\mathcal{K}_{i*}\Sigma(X_{\tilde{\mathcal{Y}}})\mathcal{K}_{i*}^T}} \\
&= \frac{\mathcal{K}_{i*}(\alpha\mathcal{Y} + \beta\mathbb{1}_{n_l}) - \mathcal{K}_{i*}\mathbb{E}(\alpha X_{\mathcal{Y}} + \beta\mathbb{1}_{n_l})}{\sqrt{\mathcal{K}_{i*}\Sigma(\alpha X_{\mathcal{Y}} + \beta\mathbb{1}_{n_l})\mathcal{K}_{i*}^T}} \\
&= \frac{\alpha\mathcal{K}_{i*}\mathcal{Y} + \beta\mathcal{K}_{i*}\mathbb{1}_{n_l} - \mathcal{K}_{i*}(\alpha\mathbb{E}(X_{\mathcal{Y}}) + \beta\mathbb{1}_{n_l})}{\sqrt{\alpha^2\mathcal{K}_{i*}\Sigma(X_{\mathcal{Y}})\mathcal{K}_{i*}^T}} \\
&= \frac{\alpha\mathcal{K}_{i*}\mathcal{Y} - \alpha\mathcal{K}_{i*}\mathbb{E}(X_{\mathcal{Y}})}{|\alpha|\sqrt{\mathcal{K}_{i*}\Sigma(X_{\mathcal{Y}})\mathcal{K}_{i*}^T}} \\
&= \frac{\mathcal{K}_{i*}\mathcal{Y} - \mathcal{K}_{i*}\mathbb{E}(X_{\mathcal{Y}})}{\sqrt{\mathcal{K}_{i*}\Sigma(X_{\mathcal{Y}})\mathcal{K}_{i*}^T}} \\
&= f_z(i)
\end{aligned}$$

□

As a consequence, the score $f_z$ is independent from the choice of the label weights: not just the final ranking is the same, but the values of the scores are identical.

**Proposition 5.** *In $f_{mc}$, the choice of $y^+$ and $y^-$ is irrelevant. More generally, computing $\tilde{f}_{mc}$ using $\tilde{\mathcal{Y}} = \alpha\mathcal{Y} + \beta\mathbb{1}_{n_l}$ instead of $\mathcal{Y}$, with $\alpha, \beta \in \mathbb{R}, \alpha > 0$, is equivalent to computing $f_{mc}$.*

*Proof.* Remember that $f_{mc}$ is defined, for the $i$-th node, as:

$$f_{mc}(i) = 1 - \frac{r_i + 1}{N + 1}$$

This measures the amount of permutations (null trials) $r_i$, out of a total of $N$, in which $f_{raw}^{null}(i) \geq f_{raw}(i)$, where $f_{raw}^{null}(i)$ is the $f_{raw}$ score of the $i$-th node using the permuted input $\mathcal{Y}_{raw}^{null}$ instead of $\mathcal{Y}_{raw}$.

We will focus on the outcome of a single random trial among the $N$ null trials, denoted by the superindex *null*. In other words, $f_{raw}^{null}(i)$ is the null score of the $i$-th node on this random trial. Let $\tilde{f}_{mc}(i)$ be the scores computed from $\tilde{\mathcal{Y}}$ instead of $\mathcal{Y}$. It suffices to prove that $f_{raw}^{null}(i) \geq f_{raw}(i) \Leftrightarrow \tilde{f}_{raw}^{null}(i) \geq \tilde{f}_{raw}(i)$. If the former is true for any permutation, then $\tilde{r}_i = r_i$ (the same $N$ random trials would lead to the same estimate), thus $\tilde{f}_{mc}(i) = f_{mc}(i)$.

$$\begin{aligned}
\tilde{f}^{null}_{raw}(i) \geq \tilde{f}_{raw}(i) &\Leftrightarrow \mathcal{K}_{i*}\tilde{\mathcal{Y}}^{null}_{raw} \geq \mathcal{K}_{i*}\tilde{\mathcal{Y}}_{raw} \\
&\Leftrightarrow \mathcal{K}_{i*}(\tilde{\mathcal{Y}}^{null}_{raw} - \tilde{\mathcal{Y}}_{raw}) \geq 0 \\
&\Leftrightarrow \mathcal{K}_{i*}(\alpha\mathcal{Y}^{null}_{raw} + \beta\mathbb{1}_{n_l} - \alpha\mathcal{Y}_{raw} - \beta\mathbb{1}_{n_l}) \geq 0 \\
&\Leftrightarrow \mathcal{K}_{i*}\alpha(\mathcal{Y}^{null}_{raw} - \mathcal{Y}_{raw}) \geq 0 \\
&\Leftrightarrow \mathcal{K}_{i*}(\mathcal{Y}^{null}_{raw} - \mathcal{Y}_{raw}) \geq 0 \\
&\Leftrightarrow f^{null}_{raw}(i) \geq f_{raw}(i)
\end{aligned}$$

□

As with $f_z$, the definition of $f_{mc}$ conveniently avoids the choice of weights for positives and negatives. Also note that propositions 4 and 5 hold for kernels based on the normalised and the unnormalised graph Laplacian. See figure 1 for examples on both properties.

### Expected values and covariance matrix of null scores

The following property provides the closed expressions of the null expected vector and covariance matrix. For instance, these are useful for computing $f_z$ without the need of permutations to estimate the expected values and variances.

**Proposition 6.** *Let $f_{raw}$ be the raw diffusion scores computed from a kernel $\mathcal{K}$ and an input $\mathcal{Y}_{raw}$. Let $X_f$ be the random vector of diffusion scores computed from a permuted input, $X_f = \mathcal{K}X_{\mathcal{Y}}$, where $X_{\mathcal{Y}}$ is the random vector that results from permuting (shuffling) $\mathcal{Y}_{raw}$, i.e. $X_{\mathcal{Y}} = \pi(\mathcal{Y}_{raw})$ for a random permutation $\pi$. Then, if $M_k = I_k - \frac{1}{k}\mathbb{1}_k\mathbb{1}_k^T$ and $n_l \geq 2$:*

*(i)* $\mathbb{E}(X_{\mathcal{Y}}) = \mu_{\mathcal{Y}}\mathbb{1}_{n_l}$

*(ii)* $\mathbb{E}(X_f) = \mu_{\mathcal{Y}}\mathcal{K}\mathbb{1}_{n_l}$

*(iii)* $\Sigma(X_{\mathcal{Y}}) = \sigma^2_{\mathcal{Y}}M_{n_l}$

*(iv)* $\Sigma(X_f) = \sigma^2_{\mathcal{Y}}\mathcal{K}M_{n_l}\mathcal{K}^T$

*being $\mu_{\mathcal{Y}} = \frac{1}{n_l}\sum_{i=1}^{n_l}\mathcal{Y}_i$ the mean of the labels and $\sigma^2_{\mathcal{Y}} = \frac{1}{n_l-1}\sum_{i=1}^{n_l}(\mathcal{Y}_i - \mu_{\mathcal{Y}})^2$ their variance.*

*Proof.* (i) $\mathbb{E}(X_{\mathcal{Y}})$ is, by symmetry, a constant $n_l$-th dimensional vector, so we can write $\mathbb{E}(X_{\mathcal{Y}}) = \mu\mathbb{1}_{n_l}$ for some $\mu \in \mathbb{R}$. Under the permutations, all the elements of the original vector have a uniform probability of ending in a given position, therefore $\mu = \frac{1}{n_l}\sum_{i=1}^{n_l}\mathcal{Y}_i = \mu_{\mathcal{Y}}$

(ii) Using property (i), $\mathbb{E}(X_f) = \mathbb{E}(\mathcal{K}X_{\mathcal{Y}}) = \mathcal{K}\mathbb{E}(X_{\mathcal{Y}}) = \mu_{\mathcal{Y}}\mathcal{K}\mathbb{1}_{n_l}$

(iii) By the symmetry of the permutations, the covariance matrix $\Sigma(X_\mathcal{Y})$ can only have two different elements: (1) the variances $\sigma^2$ on the diagonal, and (2) the covariances $\rho$ on the off-diagonal. This can be written as:

$$\Sigma(X_\mathcal{Y}) = (\sigma^2 - \rho)I_{n_l} + \rho \mathbb{1}_{n_l}\mathbb{1}_{n_l}^T$$

To find $\sigma^2$, it suffices to see that $\sigma^2$ is actually the (exact) variance of each position in the permuted vector, i.e. $\sigma^2 = \frac{1}{n_l}\sum_{i=1}^{n_l}(\mathcal{Y}_i - \mu_\mathcal{Y})^2 = \sigma_\mathcal{Y}^2 \frac{n_l - 1}{n_l}$.

To find the covariance between two positions $\rho$, it is useful to notice that $\mathbb{1}_{n_l}^T\mathcal{Y} = n_l\mu_\mathcal{Y} = \mathbb{1}_{n_l}^T X_\mathcal{Y}$ is constant, because the elements of $X_\mathcal{Y}$ must have a constant sum regardless of the permutation. Therefore, its covariance is 0:

$$\Sigma(\mathbb{1}_{n_l}^T X_\mathcal{Y}) = 0 = \mathbb{1}_{n_l}^T\Sigma(X_\mathcal{Y})\mathbb{1}_{n_l}$$

Using this, and knowing that $\mathbb{1}_{n_l}^T\mathbb{1}_{n_l} = n_l$,

$$\mathbb{1}_{n_l}^T\left[(\sigma^2 - \rho)I_{n_l} + \rho\mathbb{1}_{n_l}\mathbb{1}_{n_l}^T\right]\mathbb{1}_{n_l} = 0$$

$$(\sigma^2 - \rho)n_l + \rho n_l^2 = 0$$

$$\rho = \sigma^2\frac{-1}{n_l - 1}$$

Therefore, the desired covariance matrix is

$$\Sigma(X_\mathcal{Y}) = (\sigma^2 - \rho)I_{n_l} + \rho\mathbb{1}_{n_l}\mathbb{1}_{n_l}^T = \sigma^2\left[\frac{n_l}{n_l - 1}I_{n_l} - \frac{1}{n_l - 1}\mathbb{1}_{n_l}\mathbb{1}_{n_l}^T\right] = \sigma^2\frac{n_l}{n_l - 1}M_{n_l} = \sigma_\mathcal{Y}^2 M_{n_l}$$

(iv) Using property (iii), $\Sigma(X_f) = \Sigma(\mathcal{K}X_\mathcal{Y}) = \mathcal{K}\Sigma(X_\mathcal{Y})\mathcal{K}^T = \sigma_\mathcal{Y}^2\mathcal{K}M_{n_l}\mathcal{K}^T$

□

Note that these statistical moments are valid in the general case where $y_{raw}$ is quantitative. Another remark is that the proofs for propositions 4, 5 and the statistical moments in proposition 6 do not actually use that $\mathcal{K}$ comes from a kernel. Therefore, they stand **valid in random-walk and other approaches that can be defined as a matrix-vector product but fall outside the kernel formalism**.

**Corollary 3.** *In the absence of unlabelled nodes and using a kernel from a spectral transformation of the unnormalised Laplacian, the expected values of all the nodes is coincidental.*

*Proof.* As shown in proposition 1, if the unnormalised Laplacian is used, the kernel rows have the same sum because $\mathbb{1}_n$ is an eigenvector of $\mathcal{K} = K$ with eigenvalue $\frac{1}{r(0)}$. Using property (ii) from proposition 6 and the fact that there are no unlabelled nodes, i.e. $n_l = n$, the vector with expected values becomes

$$\mathbb{E}(X_f) = \mu_y K \mathbb{1}_n = \mu_y \frac{1}{r(0)} \mathbb{1}_n$$

Therefore, all the expected value are coincidental and equal to $\mu_y \frac{1}{r(0)}$.

□

This implies that $f_z$, in fact, modifies the ranking of $f_{raw}$ only because of different standard deviations under these conditions. This does not hold in the general case with a nonempty set of unlabelled nodes.

On the other hand, property (iv) in proposition 6 has an implication in understanding the covariance of the null distribution:

**Proposition 7.** *The covariance of the null distribution of diffusion scores is directly related to the covariance between the kernel vectors. Specifically, the covariance between two nodes is, up to a multiplicative constant, their sample covariance using as samples the labelled nodes and as features their kernel values to all the nodes.*

*Proof.* Starting from point (iv) in proposition 6, $\Sigma(X_f) = \sigma_{\mathcal{Y}}^2 \mathcal{K} M_{n_l} \mathcal{K}^T$, note that $M_k M_k = I_k - 2\frac{1}{k}\mathbb{1}_k\mathbb{1}_k^T + \frac{1}{k^2}\mathbb{1}_k k \mathbb{1}_k^T = I_k - \frac{1}{k}\mathbb{1}_k\mathbb{1}_k^T = M_k$. Also, by its own definition, $M_k$ is the matrix such that it centers the rows (resp. columns) of a matrix $A \in \mathbb{R}^{k\times k}$ when it is multiplied by the right (resp. left) of $A$.

Back to the expression of $\Sigma(X_f)$, we can write:

$$\Sigma(X_f) = \sigma_{\mathcal{Y}}^2 \mathcal{K} M_{n_l} \mathcal{K}^T = \sigma_{\mathcal{Y}}^2 \mathcal{K} M_{n_l} M_{n_l} \mathcal{K}^T = \sigma_{\mathcal{Y}}^2 \hat{\mathcal{K}} \hat{\mathcal{K}}^T$$

Being $\hat{\mathcal{K}} = \mathcal{K} M_{n_l}$ the row-centered matrix $\mathcal{K}$. This implies that the product $\hat{\mathcal{K}}\hat{\mathcal{K}}^T$ is just the sample covariance of $\mathcal{K}^T$, up to a multiplicative constant, and therefore the whole covariance $\Sigma(X_f)$ is proportional to the sample covariance of $\mathcal{K}^T$.

□

To illustrate this, a feature matrix can be built from the kernel matrix, using as *samples* the labelled nodes in the network and as *features* their similarity, using the kernel, to all the network nodes.

On one hand, the sample covariance of such a dataset (matrix whose $(i, j)$-th entry is the covariance between features $i$ and $j$) is proportional to the covariance of the null distribution in the diffusion process.

On the other hand, the leading eigenvectors of the null covariance are equivalent to the principal components (loadings) of such a dataset.

In turn, these eigenvectors are related to the spectral properties of the graph, such as the Fiedler-vector in graph partitioning [2]. The Fiedler-vector is defined as the eigenvector $v_2$ of the graph Laplacian with the second smallest eigenvalue $\lambda_2$. $v_2$ is also an eigenvector of any graph kernel as defined in [2] with eigenvalue $\frac{1}{r(\lambda_2)}$.

To wrap up these properties, we prove a particular case in which the leading eigenvectors of the null covariance convey the same data as the Fiedler-vector and its successive components.

**Proposition 8.** *Let $K$ be a kernel from a spectral transformation $r(\lambda) > 0$ on the unnormalised graph Laplacian $L$, of finite dimension. In the absence of unlabelled nodes, i.e. $n_u = 0$ and $n_l = n$, then the eigenvector $v_{i+1}$ with the $i+1$-th smallest eigenvalue of $L$, $1 \leq i \leq n-1$, is equal to the eigenvector $u_i$ with $i$-th largest eigenvalue from the null covariance $\Sigma(X_f)$.*

*Proof.* We use the definition of the kernel $K = \sum_{j=1}^{n} \frac{1}{r(\lambda_j)} v_j v_j^T$ [2], the definition of $\hat{K} = KM_n$ from property 8 and that of $\Sigma(X_f) = \sigma_y^2 \hat{K}\hat{K}^T$ from property 6. Note that the matrices $L$, $K$, $M_n$, $\hat{K}$, $\Sigma(X_f)$ belong to $\mathbb{R}^{n \times n}$ and, in particular, are square. To build the proof, we will write the eigenspaces of such matrices in ascending eigenvalues.

We denote the eigenvectors of $L$ as $v_1, v_2, \dots, v_n$, with the following eigenvalues: $0 = \lambda_1 \leq \lambda_2 \leq \dots \leq \lambda_n$ Without loss of generality, we can take $v_1 = \frac{1}{\sqrt{n}} \mathbb{1}_n = \frac{1}{\sqrt{n}}(1, \dots, 1)^T$ – that does not apply to the normalised Laplacian in general.

Because $r(\lambda) > 0$ and $r(\lambda)$ is increasing, the eigenspace of $K$ consists of the eigenvectors in reversed order $u_1 = v_n, \dots, u_n = v_1$, with eigenvalues $\frac{1}{r(\lambda_n)} \leq \dots \leq \frac{1}{r(\lambda_2)} \leq \frac{1}{r(0)}$

Regarding $M_n$, we show that its eigenspace is $w_1 = v_1, \dots, w_n = v_n$, with eigenvalues $0, 1, \dots, 1$. On one hand, $M_n v_1 = v_1 - \frac{1}{n}\mathbb{1}_n \mathbb{1}_n^T v_1 = v_1 - \frac{1}{n}\mathbb{1}_n \mathbb{1}_n^T \frac{1}{\sqrt{n}}\mathbb{1}_n = v_1 - \frac{1}{n}\mathbb{1}_n \sqrt{n} = v_1 - v_1 = 0$. On the other hand, if $1 < i \leq n$, $M_n v_i = v_i - \frac{1}{n}\mathbb{1}_n \mathbb{1}_n^T v_i = v_i - 0 = v_i$, because $v_i$ is orthogonal to $v_1$, a multiple of $\mathbb{1}_n$. We conclude that $v_i$ has eigenvalue 1.

The eigensystem of $\hat{K} = KM_n$ can be characterised using that $K$ and $M_n$ share all the eigenvectors. The eigenvalues of $\hat{K}$ are the product of the respective eigenvalues of $K$ and $M_n$. The eigenvectors $t_1 = v_1, t_2 = v_n, t_3 = v_{n-1}, \dots, t_n = v_2$ have as eigenvalues $0 \leq \frac{1}{r(\lambda_n)} \leq \frac{1}{r(\lambda_{n-1})} \leq \dots \leq \frac{1}{r(\lambda_2)}$, i.e. the leading eigenvalue of $K$, $\frac{1}{r(0)}$, has now collapsed to 0, whereas the rest are unchanged.

The proof is completed by pointing out that the eigenvectors of $\Sigma(X_f) = \sigma_y^2 \hat{K}\hat{K}^T$ are those of $\hat{K}$, and that their eigenvalues are $0 \leq \sigma_y^2 \frac{1}{r(\lambda_n)^2} \leq \sigma_y^2 \frac{1}{r(\lambda_{n-1})^2} \leq \dots \leq \sigma_y^2 \frac{1}{r(\lambda_2)^2}$. The order of the eigenvectors and the eigenvalues is preserved because $\sigma_y^2 \geq 0$ and $r(\lambda) > 0$. From here, the vector with the largest eigenvalue from $\Sigma(X_f)$, $t_n$, is equal to $v_2$, the eigenvector with second smallest eigenvalue from $L$; the second largest is $t_{n-1} = v_3$, the eigenvector with the third smallest eigenvalue from $L$, and so on.

□

**Corollary 4.** *On a graph with the same premises as in proposition 8, provided that $\lambda_2 < \lambda_3$, the leading eigenvector of the null covariance $\Sigma(X_f)$ is the Fiedler-vector, up to a change of sign.*

*Proof.* The only observation is that, given that $\lambda_2 < \lambda_3$, the Fiedler-vector is unique and, by property 8, is the leading vector of the null covariance, up to a sign change. □

Proposition 8 is illustrated in toy graphs with the lattice (figure 2) and the Barabási-Albert (figure 3) architectures. The presence of unlabelled nodes leads to a leading covariance eigenvector analogous to the Fiedler-vector, but taking into account the unobservable nature of part of the network.

In views of these results, the null covariance of a given instance can be of interest per se, because it reflects the effect of measuring (and normalising) a specific set of nodes in terms of spectral network properties.

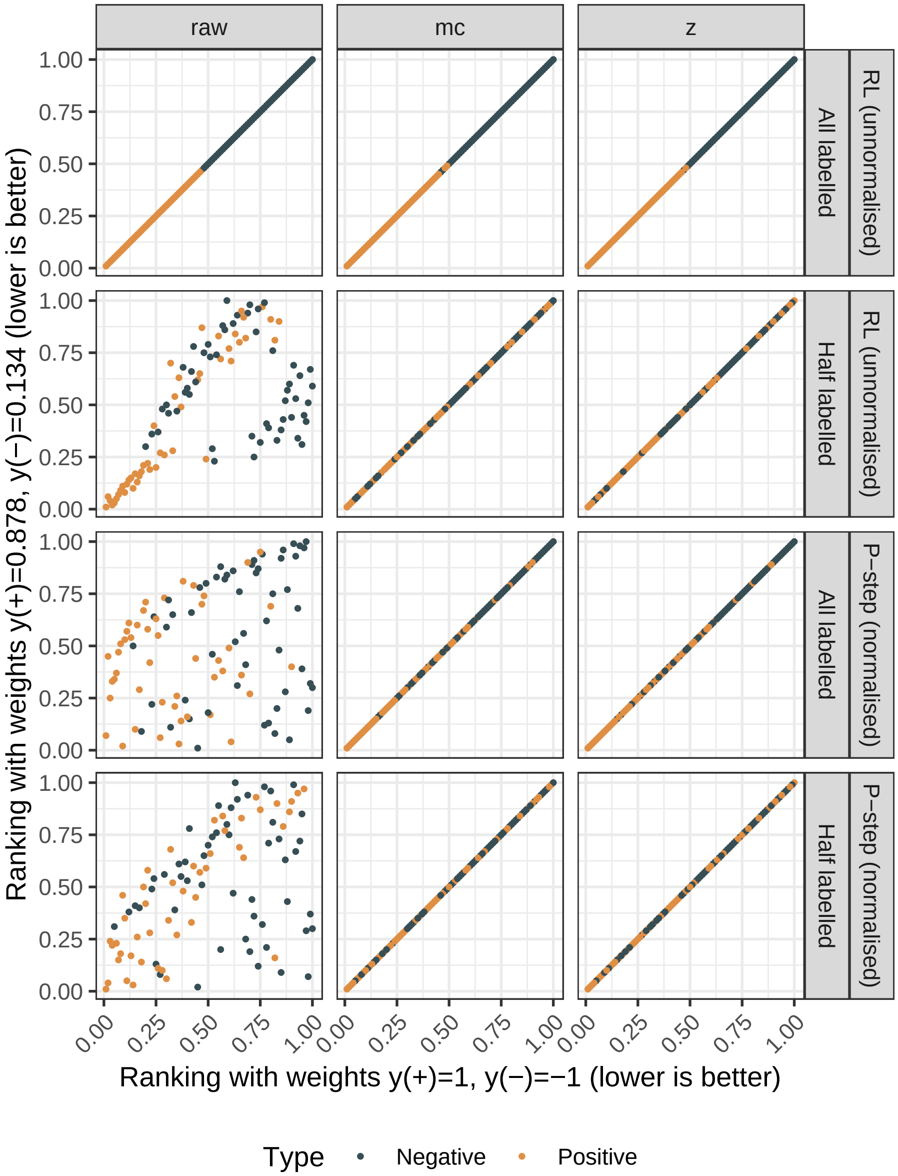


Figure 1: Effect of label encoding on $f_{raw}$, $f_{mc}$ and $f_z$, depending on (i) the kernel, and (ii) presence of unlabelled nodes. A small graph of order 100 was generated with `igraph::barabasi.game(n = 100, m = 3, directed = F)`. The 100 nodes were assigned to the positive and negative classes once, with $\frac{1}{2}$ probability each. Those labels were either all available or half available (only the first 50), considering the other half as unlabelled. Two encodings were used to rank the nodes: $y^+ = 1$, $y^- = -1$ (like $f_{ml}$) and two random weights $y^+ > y^-$. Two kernels were compared, one from the normalised Laplacian (p-step kernel, $a = 2$, $p = 5$) and one from the unnormalised (RL: regularised Laplacian, $\sigma^2 = 1$). $f_{raw}$: both encodings are equivalent only for the RL kernel without unlabelled nodes, consistent with proposition 1. $f_z$ and $f_{mc}$: as stated in propositions 4 and 5, both scores are weight-independent, regardless of the kernel and the presence of unlabelled nodes.

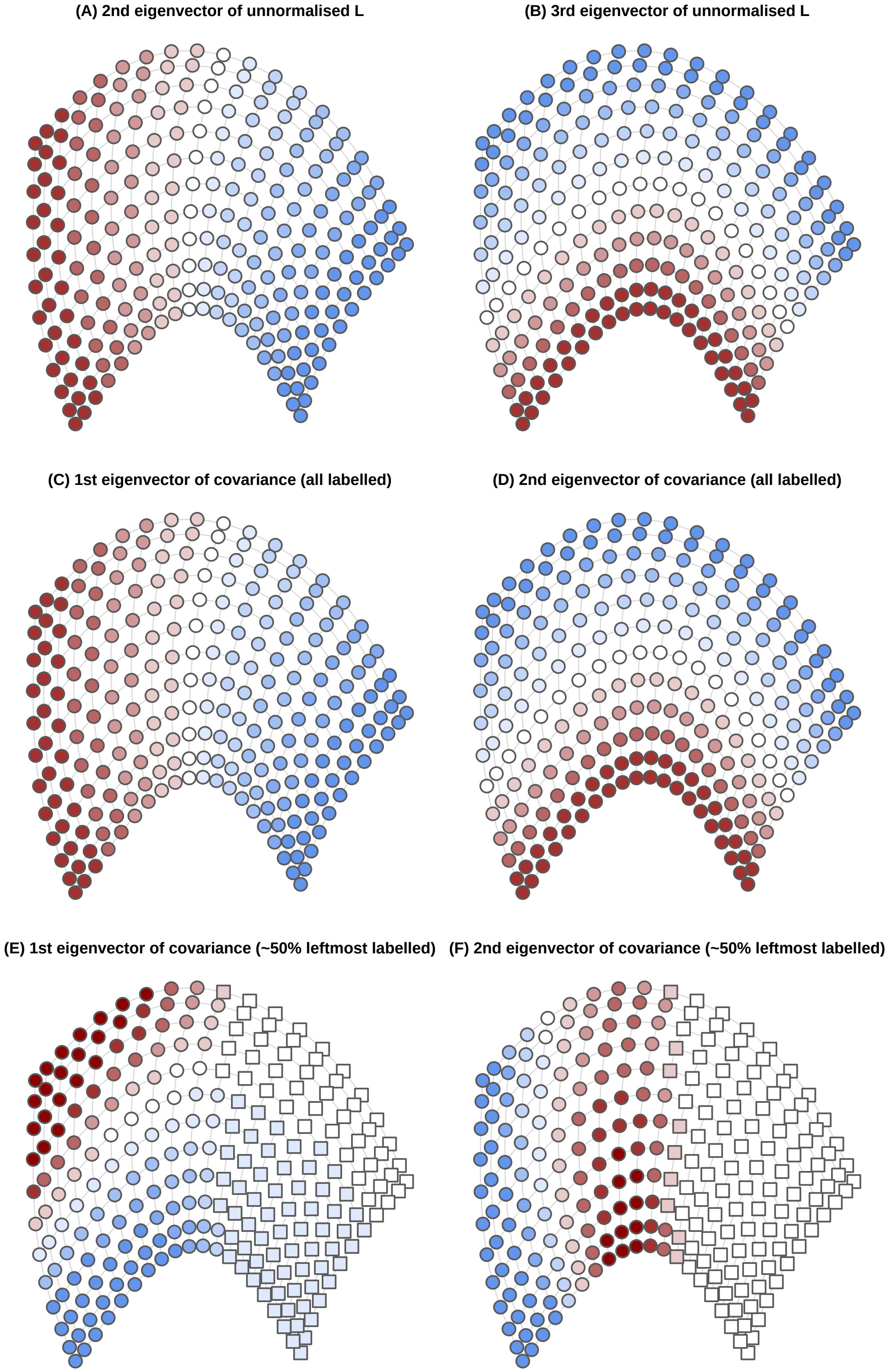


Figure 2: Comparison of the eigenvectors of the unnormalised Laplacian (panels **A-B**) and the null covariance using the regularised Laplacian kernel (panels **C-F**), on a toy lattice graph of $20 \times 12$ nodes, `igraph::graph.lattice(dimvector = c(20, 12))` in R. The null covariance is computed in two scenarios: all the nodes are labelled, i.e. $\mathcal{K} = K$ (panels **C-D**), or only half of them are, so $\mathcal{K}$ contains half of the columns in $K$ (panels **E-F**).

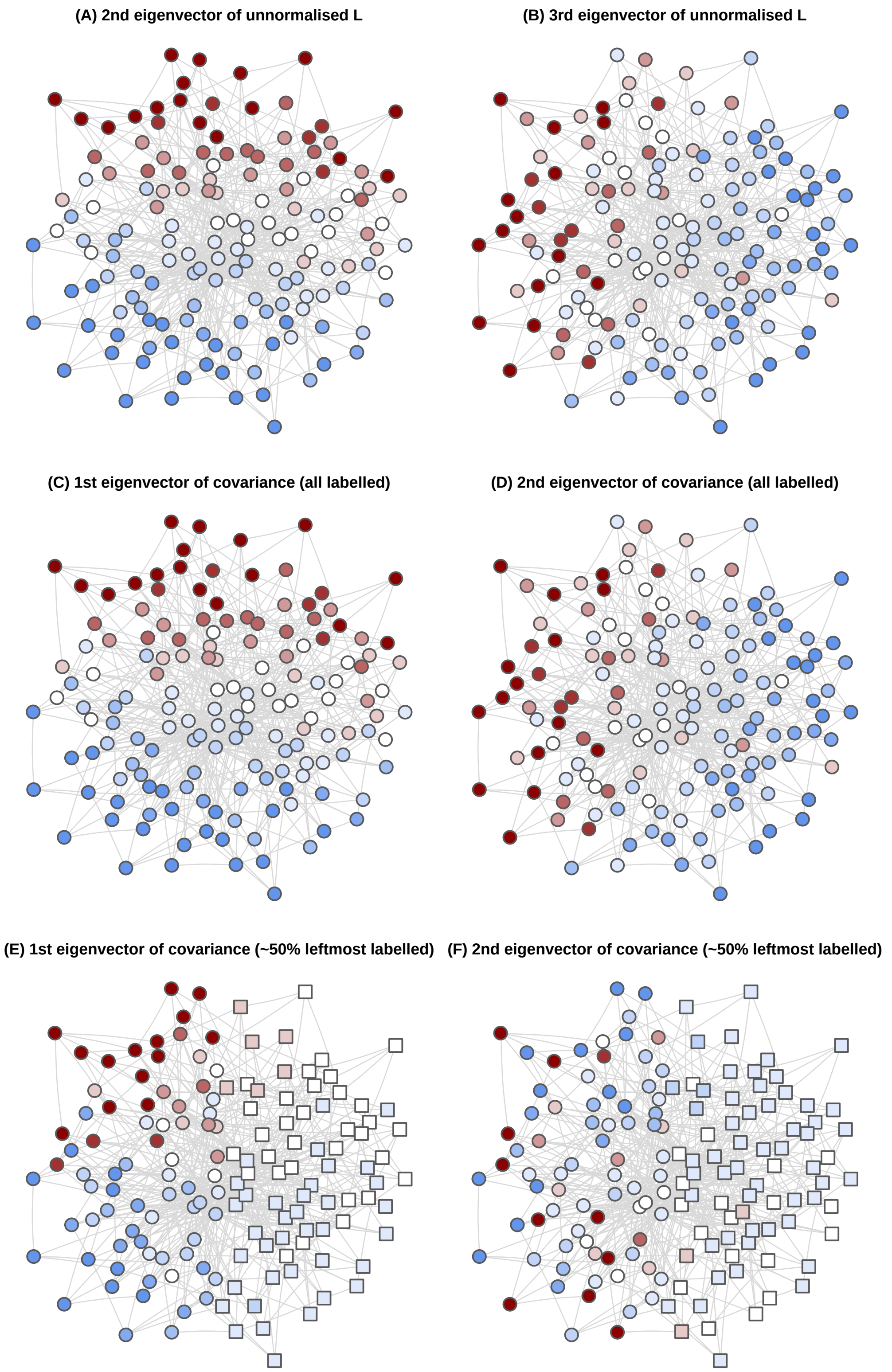


Figure 3: Comparison of the eigenvectors of the unnormalised Laplacian (panels **A-B**) and the null covariance using the regularised Laplacian kernel (panels **C-F**), on a toy synthetic Barabási graph, `igraph::barabasi.game(n = 150, m = 4, directed = F)` in R. The null covariance is computed in two scenarios: all the nodes are labelled, i.e. $\mathcal{K} = K$ (panels **C-D**), or only half of them are, so $\mathcal{K}$ contains half of the columns in $K$ (panels **E-F**).
