## Supplement 2 for "The effect of statistical normalisation on network propagation scores"

*October 30, 2018*

**4 Conclusions** **13**

**5 Metadata** **14**

**References** **15**

### 1 Introduction

This additional file contains details on the synthetic signals generated on a yeast interactome, both in biased and unbiased ways. Using a controlled environment, we characterised the behaviour of the diffusion scores to derive guidelines in terms of two key factors: the presence of bias in the positives and the class imbalance. This document can be re-built anytime by knitting its corresponding `.Rmd` file.

#### 1.1 The network

The yeast interactome was originally published in (Von Mering et al. 2002) and downloaded using the `igraphdata` R package (Csardi 2015). Only the largest connected component was used, which consisted of 2375 nodes and $1.1693 \times 10^4$ edges. A summary of the network is provided below:

```
## IGRAPH 410ccdc UN-- 2375 11693 -- Yeast protein interactions, von Mering e
## + attr: name (g/c), Citation (g/c), Author (g/c), URL (g/c),
## | Classes (g/x), name (v/c), Class (v/c), Description (v/c),
## | Confidence (e/c)
```

#### 1.2 Synthetic signal generation

Biased and unbiased signals were generated in order to compare normalised and unnormalised diffusion scores. As shown in the diffusion scores properties in Supplement 1, if all the nodes are labelled and the regularised unnormalised Laplacian kernel is used, then the expected values of the null distribution are constant for all the nodes in the network. In order to have differences in expected values (and therefore noticeable biases), nodes were randomly divided in three classes:

- Labelled nodes: the labelled nodes in the input
- Target nodes: the unlabelled nodes that had to be prioritised
- Filler nodes: the rest of unlabelled nodes

The presence of filler and target nodes, considered as unlabelled in the diffusion inputs, promoted differences in the expected values of all nodes. Each class contained around one third of the nodes:

```
##
##   Filler Labelled   Target
##      793      791      791
```

The purpose was to sample $n_{labelled}$ nodes from the labelled nodes and $n_{target}$ nodes from the target nodes in each instance. The sampled nodes were deemed positives, whereas the rest were negatives. Diffusion scores were fed with the labelled nodes in order to prioritise the target nodes, on which the performance metrics were computed.

##### 1.2.1 Biased sampling

First, the $n_{labelled}$ nodes were uniformly sampled from the labelled nodes, giving a binary input vector $y$. Then, the raw scores were computed: $f_{raw} = Ky$. Exactly $n_{target}$ nodes were sampled, where the probability of the $i$-th node was proportional to $f_{raw}(i)$. This sampling scheme was biased because, by hypothesis, nodes with higher expected value would become positives more frequently.

##### 1.2.2 Unbiased sampling

Like in the biased sampling, the $n_{labelled}$ nodes were uniformly sampled to obtain the binary input vector $y$. The $n_{target}$ nodes were sampled with a probability proportional to $f_{mc}(i) + \frac{1}{N+1}$, where $N = 10^4$ is the

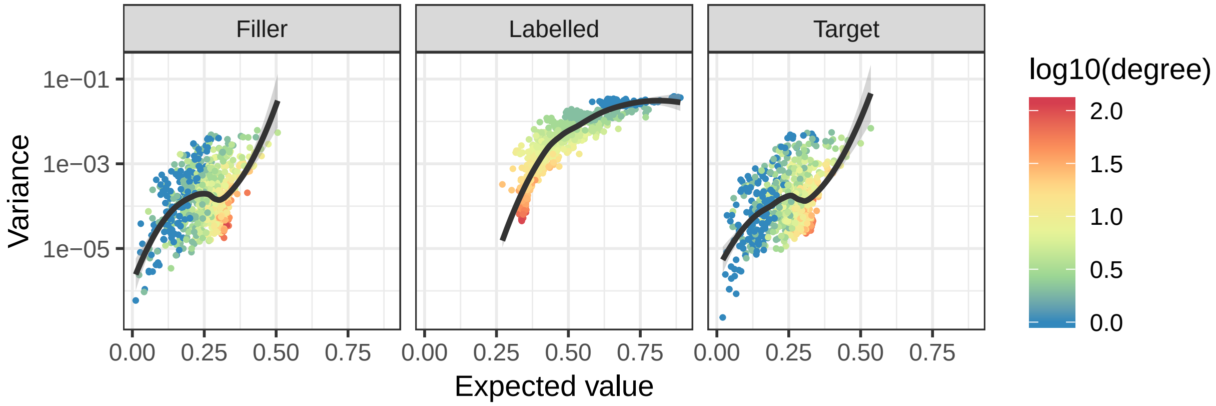

Figure 1: Expected value, variance and node degree in every node category. Loess fit in black, shaded with 0.95 confidence interval. Note how the effect of node degree on its expected value had opposed directions in labelled and unlabelled nodes. Differences were also present in the magnitude of expected values, variances, and their trend.

number of simulations. $f_{mc}(i)$ was (roughly) their empirical cumulative distribution function applied to the scores $f_{raw} = Ky$, which removed the bias by its own definition.

### 2 Descriptive statistics

#### 2.1 Expected value and covariance matrices

After defining the node classes and the basic input parameters, we computed the theoretical mean vector and covariance matrix. The fact that the number of positives in the input was constant in these simulations led to fixed $\mu_y$ and $\sigma_y^2$ values, allowing a single representation of the expected values and variances of the null distributions in figure 1. The figure confirms that *labelled* nodes exhibited properties different that those of *filler* and *target* nodes: *labelled* nodes had higher expected values, variances, and different trends between expected value, variance, and degree. Likewise, *filler* and *target* nodes were undistinguishable, expected by their definition.

Figure 2 offers a closer look at differences in reference mean values the *target* nodes, which is a property of the network. The *target* nodes were of special interest because predictions and performance metrics were computed on them.

#### 2.2 Input lists

A total of 100 biased and 100 unbiased instances were generated, each with a proportion of 0.1 labelled nodes and a proportion of 0.1 target nodes with positive labels. To generate the unbiased inputs, `mc` scores were computed by permuting $10^4$ times. The regularised (unnormalised) Laplacian kernel was used.

The frequency of target nodes and the reference expected value were expected to be uncorrelated in the unbiased signals, whilst positively correlated in the biased case. By definition, the input nodes should be independent from the reference expected value as well. Figure 3 supports all the claims above.

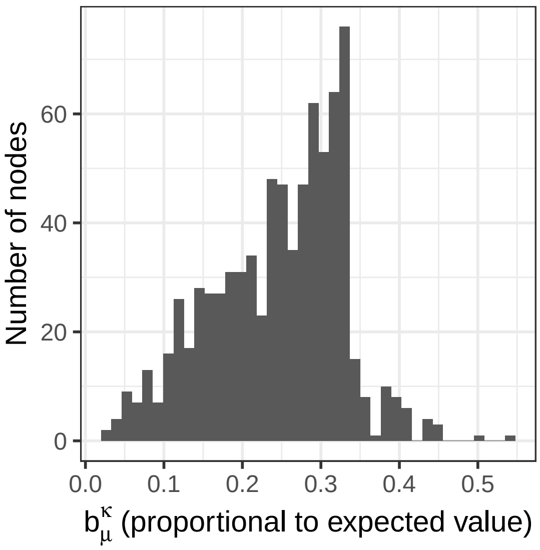

Figure 2: Histogram of the reference expected value of the target nodes.

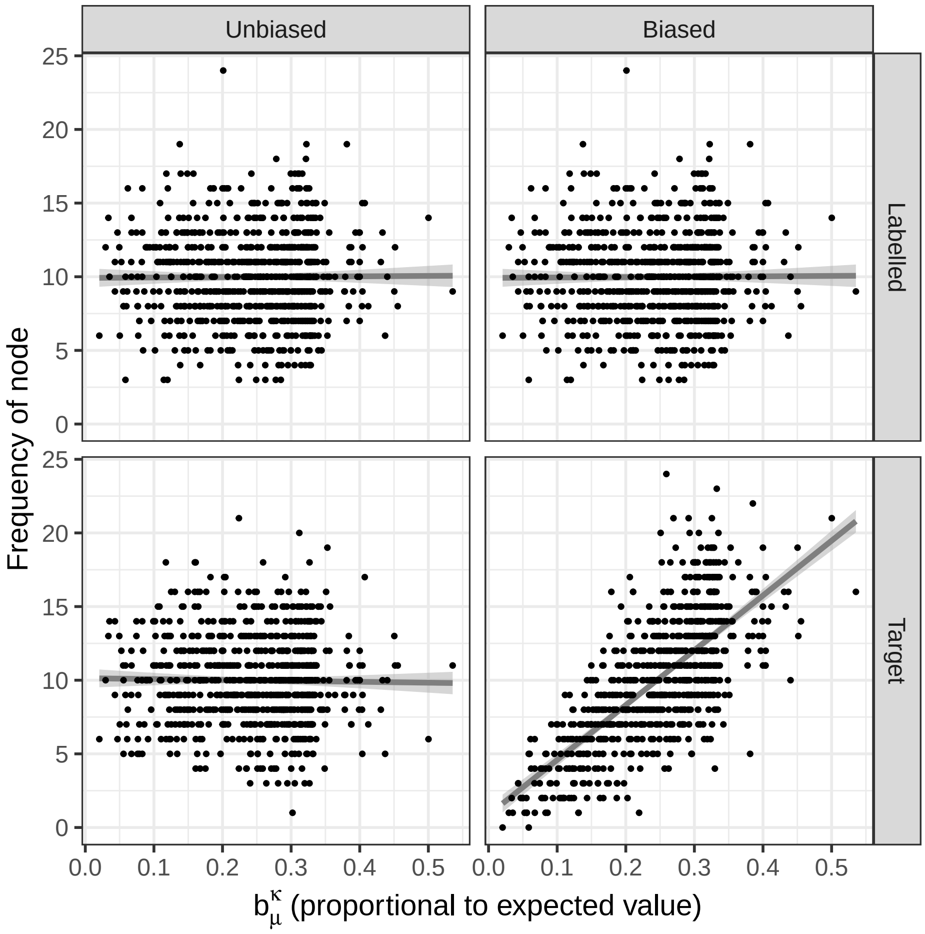

Figure 3: Frequency of positive nodes among the targeted nodes, as a function of the node reference expected value. Gray lines correspond to linear models with a 0.95 confidence interval.

##### 2.3 Diffusion scores

$10^4$ permutations were used to compute `mc` and `ber_p` scores. Figure 4 compares the rankings from each method, stratified by positives and negatives, and shows their correlation. This suggests groups of methods with similar behaviours: (i) `ml` and `gm`, or (ii) `ber_p`, `mc` and `z`. Also, top ranked `raw` nodes were usually top ranked in `z`, but the converse was not true.

Figure 5 depicts the correlations between methods. First, this shows how `ber_s` is equivalent to `raw` in terms of ranking, as proven in the properties in Supplement 1. `raw` correlates with normalised scores `mc` and `z`, the hybrid `ber_p` and `pagerank`. On the other hand, `pagerank` strongly anticorrelates with `ml` and `gm` (`raw` does as well, but only slightly). The scores `ml` and `gm` diffuse $-1$ on the negatives, which outnumber the positives 9 to 1 and dominate them. The nodes are expected to be ranked roughly by the (negative) reference expected value, also correlated with `pagerank`. This is supported by the strong anticorrelation between the node ranking (`ml`, `gm`) and `pagerank`.

Figure 6 depicts the concordance between the top 10 ranked nodes under each diffusion score. This scenario is slightly different from that in figure 5 and suggests that methods with highest similarity are (i) `raw` and `ber_p`, (ii) `ml` and `gm`, (iii) `mc` and `z`.

##### 2.4 Bias within prioritisations

Our main hypothesis on the fundamental impact of normalising the scores (`mc`, `z` versus `raw`) was that normalisation attained a more uniform power across the nodes of the network. In other words, unnormalised scores kept a higher power on a certain kind of nodes, driven by the reference expected value $b_\mu^\mathcal{K}$ in the null distribution. Figure 7 illustrates this behaviour, present in biased and unbiased signals: positives with high $b_\mu^\mathcal{K}$ were top ranked by `raw`, at the expenses of missing positives with low $b_\mu^\mathcal{K}$. However, the overall impact on performance was not obviously derived from figure 7 alone, because we needed to account for the density of true positives across the reference expected value (i.e. "are the positive nodes biased?"), as shown in figure 3.

Other remarks from figure 7: `ber_p` behaves halfway between `raw` and `mc`; and `ml` is biased in the other direction, that is, favouring nodes with a low reference expected value. The latter relates with the prior observations on how `ml` anticorrelates with `pagerank` because it diffuses $-1$ on the negatives, which outnumber the positives. Figure 7 casts doubt on the peformance of `ml` because the mean ranking of positives and negatives is qualitatively indistinguishable at the low reference expected value region, where `ml` should excel.

#### 3 Performance

##### 3.1 Aditive model

The AUROC and AUPRC were computed for each diffusion score and input, with its corresponding simulated ground truth (figure 8). Each box contained 100 data points. Differences were described in terms of the following additive quasibinomial (logit link) model, summarised in table 1:

$$\text{performance} \sim \text{method} + \text{method:biased} + \text{metric}$$

Provided that AUROC and AUPRC showed similar trends (figure 8) and that both range between 0 and 1, they were combined and modelled with the `metric` categorical covariate. `biased` referred to the nature of the signal, biased or unbiased.

Figure 8 suggests that the unnormalised scores `raw` were preferrable if the signal was biased, whereas `mc` and `z` were best suited for unbiased signals. Likewise, the hybrid scores `ber_p` standed out as a good compromise between both.

Table 2 contains confidence intervals on the predictions of the model for each combination of factors.

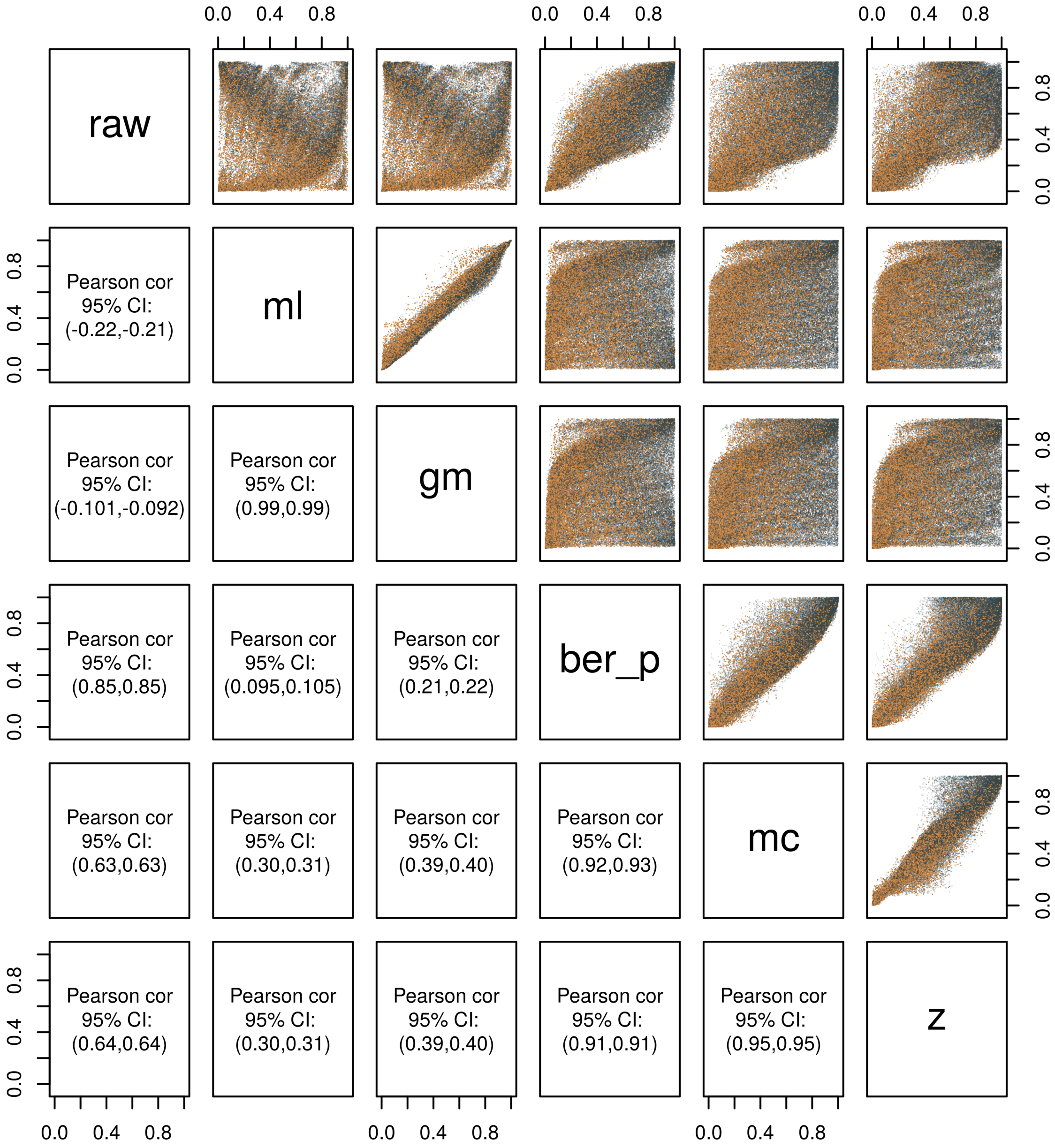

Figure 4: Pairs plot between the rankings by each diffusion score (and baseline). Top-ranked nodes are closer to 0. Positives and negatives are represented in orange and gray. The color legend has an adjusted transparency that corrects the fact that negatives greatly outnumbered positives.

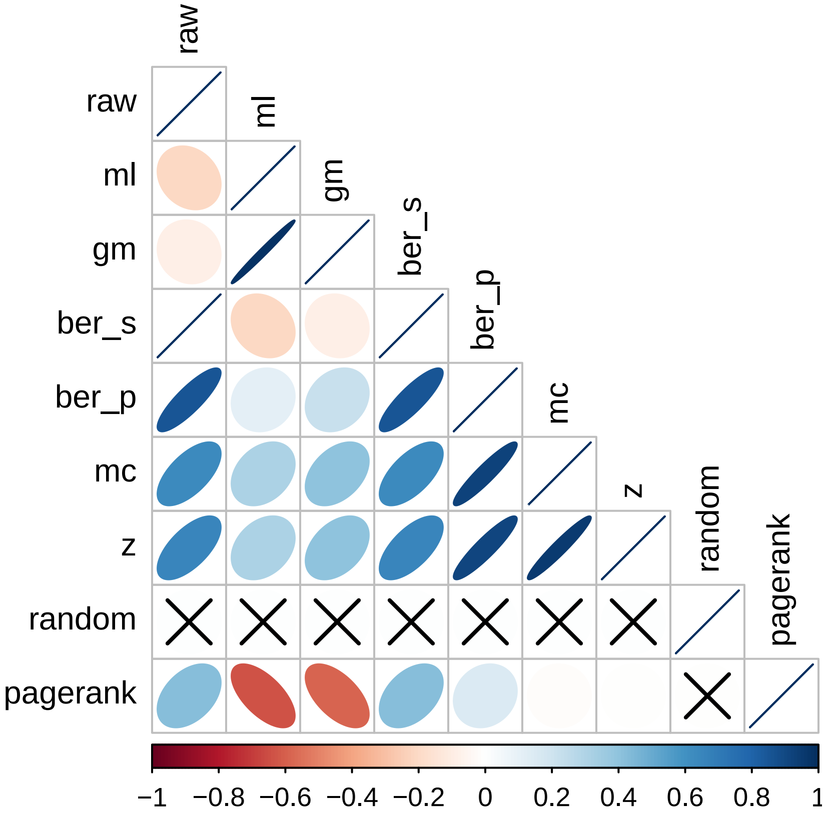

Figure 5: Spearman correlation between node rankings for all the diffusion scores and baselines. Crossed correlations had false discovery rates larger than 0.05.

Table 1: Quasibinomial model for AUROC and AUPRC. Estimates with 0.95 confidence intervals.

| | |
|---|---|
| methodml | $-0.848^{***}$ $(-0.884, -0.811)$ |
| methodgm | $-0.719^{***}$ $(-0.755, -0.683)$ |
| methodber_p | $-0.086^{***}$ $(-0.122, -0.051)$ |
| methodmc | $-0.304^{***}$ $(-0.339, -0.268)$ |
| methodz | $-0.266^{***}$ $(-0.302, -0.230)$ |
| methodrandom | $-0.895^{***}$ $(-0.932, -0.859)$ |
| methodpagerank | $-0.653^{***}$ $(-0.690, -0.617)$ |
| metricAUROC | $2.131^{***}$ $(2.118, 2.145)$ |
| methodraw:biasedUnbiased | $-0.455^{***}$ $(-0.491, -0.419)$ |
| methodml:biasedUnbiased | $0.155^{***}$ $(0.118, 0.192)$ |
| methodgm:biasedUnbiased | $0.092^{***}$ $(0.056, 0.129)$ |
| methodber_p:biasedUnbiased | $-0.186^{***}$ $(-0.221, -0.150)$ |
| methodmc:biasedUnbiased | $0.075^{***}$ $(0.039, 0.111)$ |
| methodz:biasedUnbiased | $0.011$ $(-0.025, 0.047)$ |
| methodrandom:biasedUnbiased | $-0.031$ $(-0.069, 0.006)$ |
| methodpagerank:biasedUnbiased | $-0.271^{***}$ $(-0.308, -0.234)$ |
| Constant | $-1.229^{***}$ $(-1.255, -1.202)$ |
| Observations | 3,200 |
| *Note:* | $^{*}$p<0.05; $^{**}$p<0.01; $^{***}$p<0.001 |

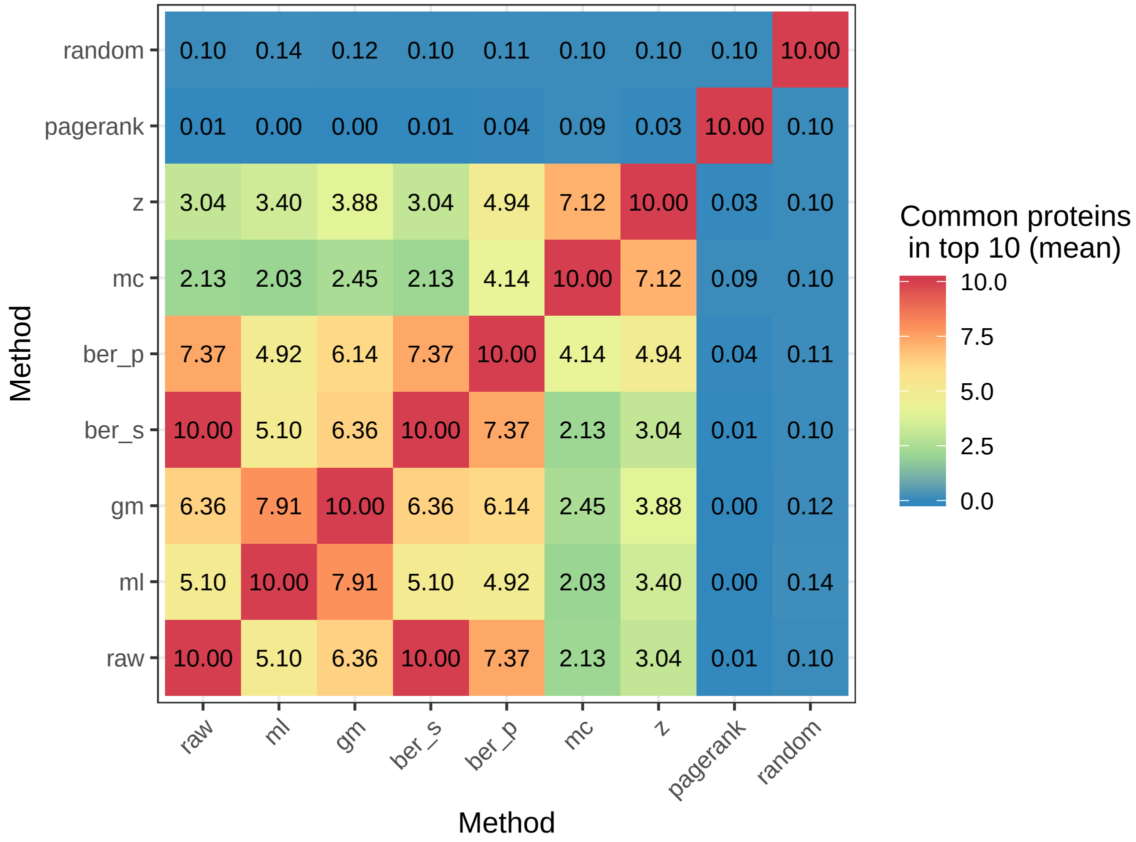

Figure 6: Common hits within the top 10 suggestions of all methods.

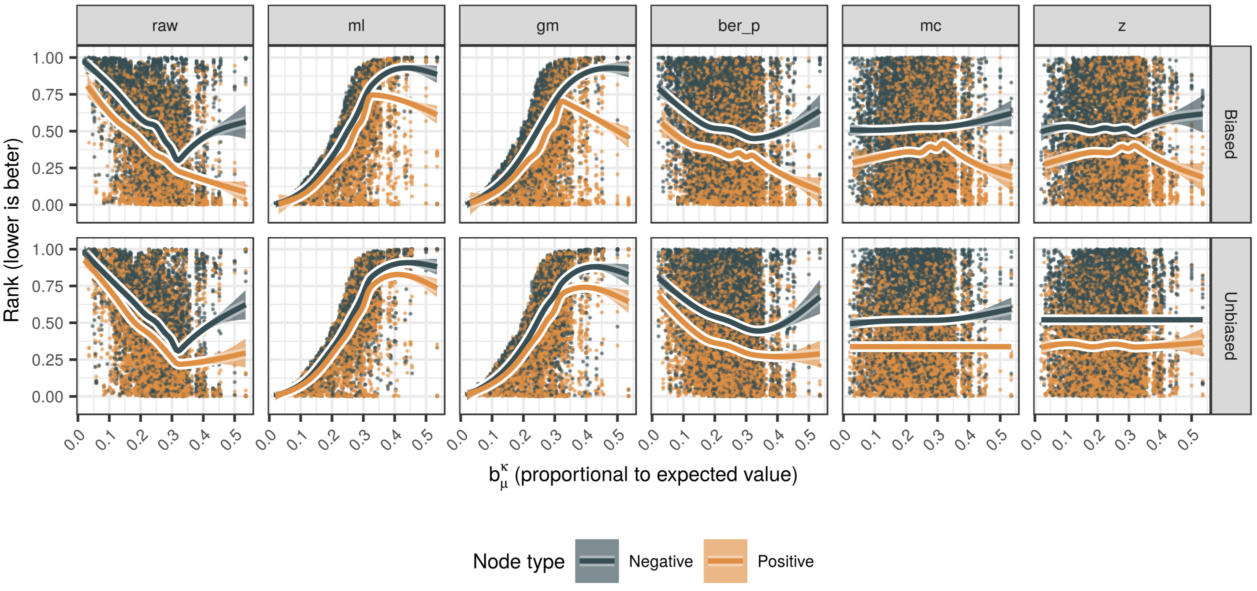

Figure 7: Ranking of the positives and negatives within the target nodes as a function of their reference expected value, divided by method and signal bias. Best rankings are those close to 0. The smoothing was fitted using the default gam method in ggplot2. For visual and computational purposes, only a fraction of 0.1 of the negatives were represented.

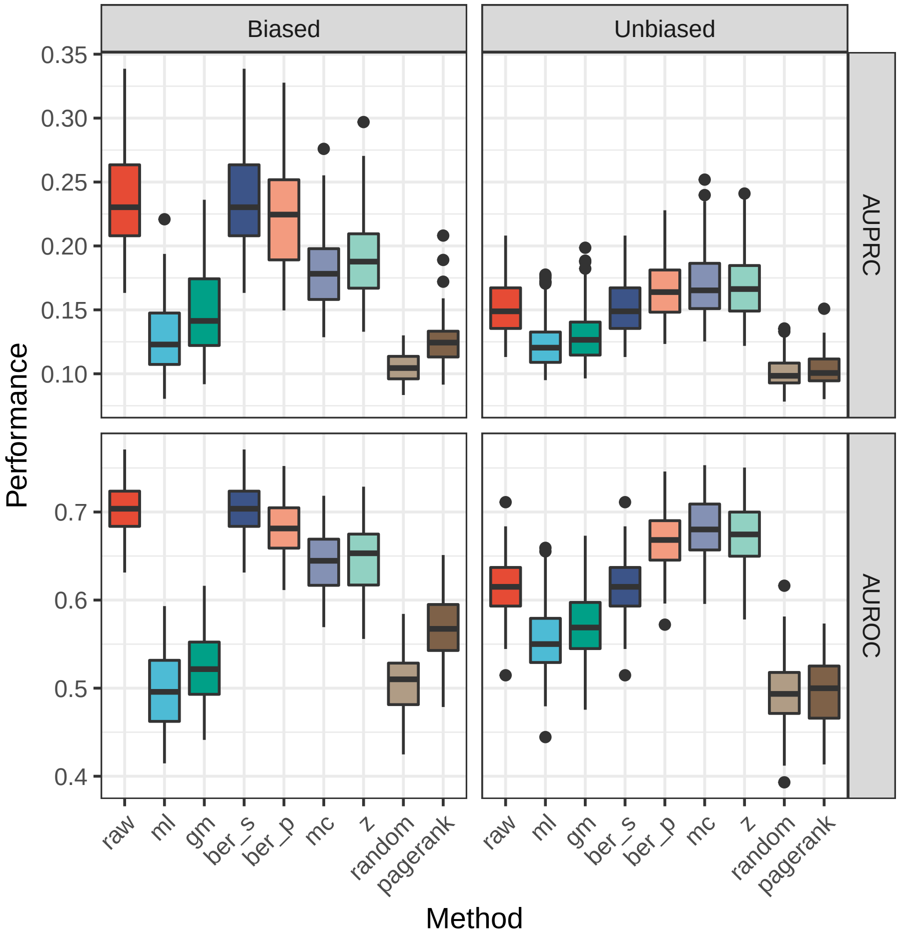

Figure 8: AUROC and AUPRC of diffusion scores for biased and unbiased signals.

| Bias_signal | Method | AUPRC | AUROC |
|---|---|---|---|
| Biased | raw | (0.222, 0.231) | (0.706, 0.717) |
| Biased | ml | (0.109, 0.114) | (0.507, 0.520) |
| Biased | gm | (0.122, 0.128) | (0.539, 0.552) |
| Biased | ber_p | (0.207, 0.216) | (0.688, 0.699) |
| Biased | mc | (0.174, 0.182) | (0.639, 0.651) |
| Biased | z | (0.179, 0.187) | (0.648, 0.660) |
| Biased | random | (0.104, 0.110) | (0.495, 0.508) |
| Biased | pagerank | (0.129, 0.135) | (0.555, 0.568) |
| Unbiased | raw | (0.153, 0.160) | (0.604, 0.616) |
| Unbiased | ml | (0.125, 0.131) | (0.546, 0.559) |
| Unbiased | gm | (0.132, 0.138) | (0.562, 0.575) |
| Unbiased | ber_p | (0.178, 0.186) | (0.647, 0.658) |
| Unbiased | mc | (0.185, 0.193) | (0.657, 0.668) |
| Unbiased | z | (0.181, 0.189) | (0.651, 0.662) |
| Unbiased | random | (0.101, 0.107) | (0.487, 0.501) |
| Unbiased | pagerank | (0.101, 0.107) | (0.488, 0.501) |

Table 2: Confidence intervals (0.95) on predicted AUROC and AUPRC.

We tested for differences between the predictions of `raw` and `z` in the four cases using Tukey's method, confirming that the differences discussed above were statistically significant:

```
##   metric                  contrast   estimate         SE  df   z.ratio
## 1  AUPRC     Biased,raw - Biased,z  0.2657691 0.01824644 Inf  14.56553
## 2  AUPRC Unbiased,raw - Unbiased,z -0.2006233 0.01836834 Inf -10.92223
## 3  AUROC     Biased,raw - Biased,z  0.2657691 0.01824644 Inf  14.56553
## 4  AUROC Unbiased,raw - Unbiased,z -0.2006233 0.01836834 Inf -10.92223
##        p.value
## 1 0.000000e+00
## 2 2.137179e-13
## 3 0.000000e+00
## 4 2.137179e-13
```

Another interesting remark from figure 8: `pagerank` had predictive power only in the biased setup. The predictive power of an input-naive centrality measure like `pagerank` can be a reason to suspect that the signal is biased towards high-degree nodes.

```
##   metric                            contrast     estimate         SE  df
## 1  AUPRC     Biased,random - Biased,pagerank -0.241916366 0.01888432 Inf
## 2  AUPRC Unbiased,random - Unbiased,pagerank -0.002012079 0.01915065 Inf
## 3  AUROC     Biased,random - Biased,pagerank -0.241916366 0.01888432 Inf
## 4  AUROC Unbiased,random - Unbiased,pagerank -0.002012079 0.01915065 Inf
##       z.ratio p.value
## 1 -12.8104369       0
## 2  -0.1050659       1
## 3 -12.8104369       0
## 4  -0.1050659       1
```

#### 3.2 Correlation between method performances

Finally, we examined the similarities between diffusion scores at the performance level. Figure 9 shows the Spearman correlation between the performance metrics of the diffusion scores. Small differences were observed between AUROC and AUPRC: `ml` and `gm` correlated with `raw` with AUPRC but not with AUROC.

In general, all the proper diffusion scores tended to correlate, even more than we observed in figure 5.

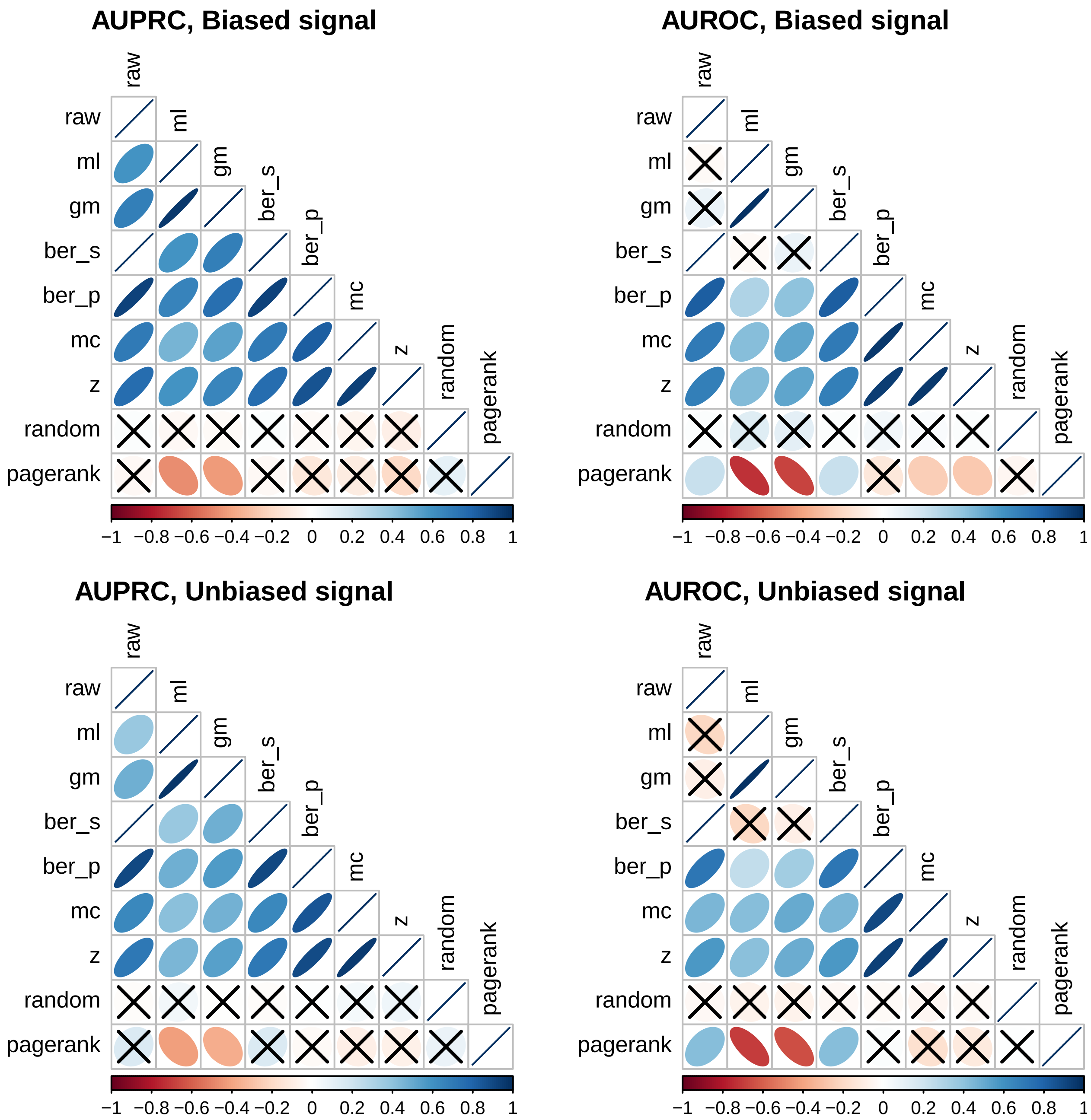

Figure 9: Spearman correlation between performance metrics of diffusion scores and baselines. Correlations not significant at $p < 0.05$ after multiple testing were crossed out.

### 4 Conclusions

The main findings from this proof of concept:

- Our definitions of biased and unbiased signals seemed consistent: unbiased signals were uncorrelated with the reference expected value, whereas biaseds ones showed a positive correlation.
- Changing the labels for diffusion (e.g. `ml` versus `raw`) and normalising the scores had a noticeable impact on the prioritisations and their performance.
- `mc` and `z` had a similar behaviour. This was expected, as they are the parametric and non-parametric alternatives for normalising.
- The adequateness of normalising lied on the distribution of positives across the reference expected value $b_{\mu}^{\mathcal{K}}$. Biased signals favoured `raw` by definition, whereas `mc` and `z` were preferrable on unbiased signals. Even within a hypothetical case study without overall performance differences, `raw` and `z`/`mc` would be expected to behave differently.
- Class imbalance backfired in `ml` and `gm`, as the properties of the negatives overshadowed those of the positives. A hypothetical case where positives outnumbered negatives might cause a similar effect `raw` as well.
- The complementarity of `raw` and `mc` leaves `ber_p` as a good compromise between both.

### 5 Metadata

```
## [1] "Sun Jan 19 17:57:29 2020"
```

```
## R version 3.5.3 (2019-03-11)
## Platform: x86_64-pc-linux-gnu (64-bit)
## Running under: Ubuntu 16.04.6 LTS
##
## Matrix products: default
## BLAS: /usr/lib/atlas-base/atlas/libblas.so.3.0
## LAPACK: /usr/lib/atlas-base/atlas/liblapack.so.3.0
##
## locale:
##  [1] LC_CTYPE=en_US.UTF-8       LC_NUMERIC=C
##  [3] LC_TIME=en_US.UTF-8        LC_COLLATE=en_US.UTF-8
##  [5] LC_MONETARY=en_US.UTF-8    LC_MESSAGES=en_US.UTF-8
##  [7] LC_PAPER=en_US.UTF-8       LC_NAME=C
##  [9] LC_ADDRESS=C               LC_TELEPHONE=C
## [11] LC_MEASUREMENT=en_US.UTF-8 LC_IDENTIFICATION=C
##
## attached base packages:
## [1] grid      stats     graphics  grDevices utils     datasets  methods
## [8] base
##
## other attached packages:
##  [1] bindrcpp_0.2.2    xtable_1.8-3      data.table_1.11.8
##  [4] extrafont_0.17    gtable_0.2.0      GGally_1.4.0
##  [7] ggsci_2.9         ggplot2_3.1.0     tidyr_0.8.2
## [10] dplyr_0.7.8       plyr_1.8.4        reshape2_1.4.3
## [13] magrittr_1.5      diffuStats_1.2.0  igraphdata_1.0.1
## [16] igraph_1.2.2
##
## loaded via a namespace (and not attached):
##  [1] Rcpp_1.0.0               mvtnorm_1.0-8
##  [3] lattice_0.20-38          zoo_1.8-4
##  [5] assertthat_0.2.0         rprojroot_1.3-2
##  [7] digest_0.6.18            R6_2.3.0
##  [9] backports_1.1.2          evaluate_0.12
## [11] pillar_1.3.0             rlang_0.3.0.1
## [13] lazyeval_0.2.1           multcomp_1.4-8
## [15] extrafontdb_1.0          Matrix_1.2-15
## [17] rmarkdown_1.10           labeling_0.3
## [19] splines_3.5.3            stringr_1.3.1
## [21] munsell_0.5.0            compiler_3.5.3
## [23] xfun_0.4                 pkgconfig_2.0.2
## [25] mgcv_1.8-27              htmltools_0.3.6
## [27] tidyselect_0.2.5         tibble_1.4.2
## [29] expm_0.999-3             bookdown_0.7
## [31] codetools_0.2-16         reshape_0.8.8
## [33] crayon_1.3.4             withr_2.1.2
## [35] MASS_7.3-51.1            nlme_3.1-137
## [37] Rttf2pt1_1.3.7           scales_1.0.0
## [39] RcppParallel_4.4.1       estimability_1.3
## [41] stringi_1.2.4            RcppArmadillo_0.9.200.4.0
```

```
## [43] sandwich_2.5-0          TH.data_1.0-9
## [45] stargazer_5.2.2         RColorBrewer_1.1-2
## [47] tools_3.5.3             glue_1.3.0
## [49] purrr_0.2.5             emmeans_1.3.0
## [51] survival_2.43-3         yaml_2.2.0
## [53] colorspace_1.3-2        corrplot_0.84
## [55] knitr_1.20              bindr_0.1.1
## [57] precrec_0.9.1
```
