## Supplement 3 for "The effect of statistical normalisation on network propagation scores"

*October 15, 2018*

**4 Reproducibility** **15**

**References** **17**

### 1 Introduction

This additional file contains details on the DLBCL dataset, the human proteome and the synthetic signals generated on it. This document can be re-built anytime by knitting its corresponding `.Rmd` file.

#### 1.1 The network

We used the HPRD network (Mishra et al. 2006) as used in the DLBCL package (M. Dittrich and Beisser 2010), which provides a case study for the BioNet R package (M. T. Dittrich et al. 2008). Below is a summary of the network, obtained by taking the largest connected component from the original network `interactome`:

```
## IGRAPH e764cea UNW- 8989 34325 --
## + attr: kegg_mapped (g/x), info (g/c), name (v/c), geneID (v/c),
## | geneSymbol (v/c), obs_lym (v/l), obs_all (v/l), weight (e/n)
```

The network contained 8989 nodes and 34325 edges and was connected by construction. The edges were unweighted, as they had a constant, unitary weight:

```
##    Min. 1st Qu.  Median    Mean 3rd Qu.    Max.
##       1       1       1       1       1       1
```

### 2 Descriptive statistics

#### 2.1 Simulated signals

Signals to benchmark the diffusion scores were obtained by sub-sampling the Kyoto Encyclopedia of Genes and Genomes (KEGG) pathways (Kanehisa et al. 2017), using the release:

```
## [1] "T01001           Homo sapiens (human) KEGG Genes Database"
## [2] "hsa              Release 83.0+/09-09, Sep 17"
## [3] "                 Kanehisa Laboratories"
## [4] "                 39,524 entries"
```

Pathways were used like gene sets, without taking further network data from the KEGG database. After mapping the pathways to the network, their size followed the distribution in figure 1. Only pathways with a minimum of $N_{min} = 30$ genes were considered.

Likewise, figure 2 depicts the amount of pathways in which each gene participates. Although some genes are ubiquitous, most of them belong to less than 10 pathways.

#### 2.2 Sub-sampling

The sub-sampling was governed by three key parameters: the number of affected pathways $k \in \{1, 3, 5, 10\}$, the proportion of differentially expressed genes $r \in \{0.3, 0.5, 0.7\}$ and the maximum p-value for differential expression, $p_{max} \in \{0.01, 0.001, 10^{-4}, 10^{-5}\}$. The extreme values $k = 10$ and $p_{max} = 10^{-5}$ led to redundant results and were left out of the main analyses.

As described in the main body, in each run $k$ pathways were uniformly sampled and their genes were tagged as positives. A proportion of $r$ positives was uniformly sampled to show differential expression, with their p-values uniformly sampled in $[0, p_{max}]$. The remaining proportion of $1 - r$ genes were not differentially expressed, imposed by sampling their p-values uniformly in $[0, 1]$. For each combination of parameters, a total of $N = 50$ repetitions were generated. Regardless of which nodes were considered as *unlabelled* or *labelled*, the p-values were generated for all the nodes in the network.

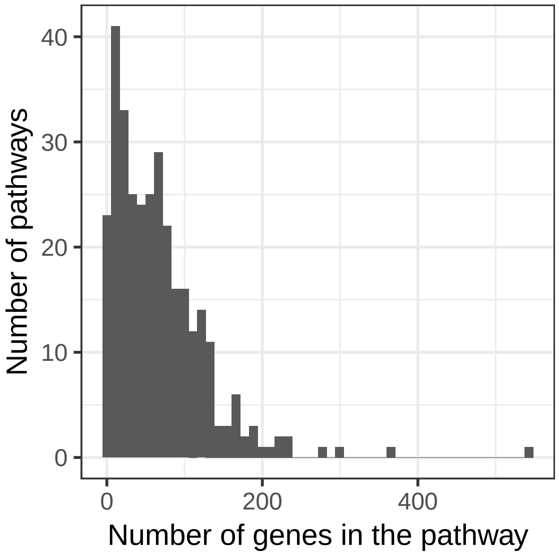

Figure 1: Histogram with number of pathways involving each gene

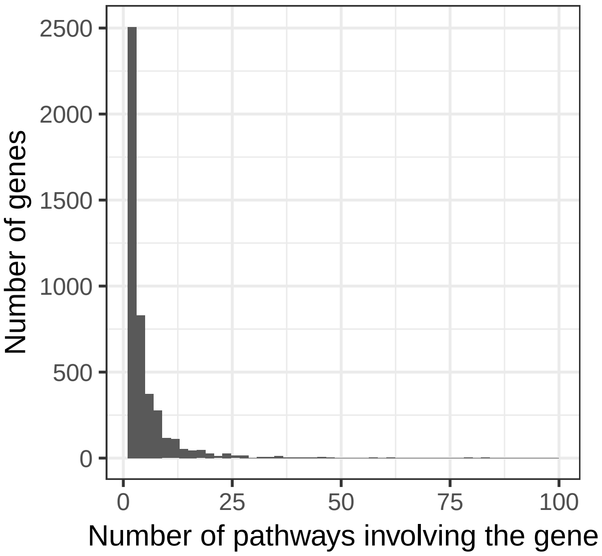

Figure 2: Histogram with number of genes in each pathway

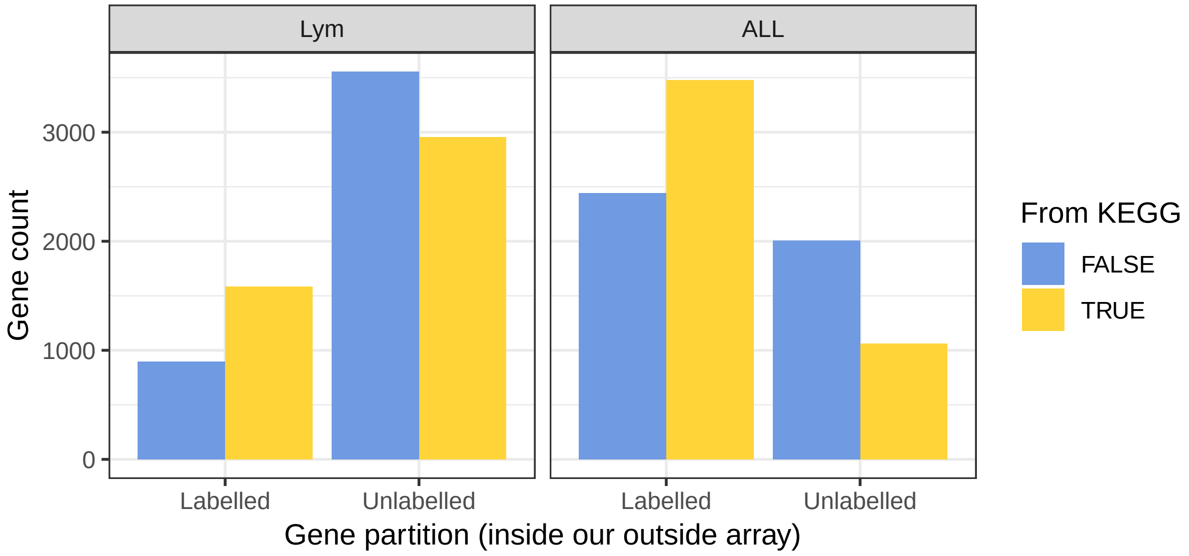

Figure 3: Number of genes inside and outside pathways, stratified by observability and array

#### 2.3 Array-based backgrounds

In order to evaluate the effect of the statistical background, genes from the network were partitioned by **observability** into *labelled* and *unlabelled*. *Labelled* nodes were defined as those belonging to an array, whereas *unlabelled* nodes were those outside it. Two arrays were used: the *ALL* array (Chiaretti et al. 2004), obtained from the ALL R package (Li 2009), and the *Lym* array (Rosenwald et al. 2002) from the DLBCL package (M. Dittrich and Beisser 2010). Gene identifiers in *ALL* were mapped to the network through `BioNet::mapByVar()` from the BioNet package (M. T. Dittrich et al. 2008), whereas those of *Lym* were already mapped in the data package. Each array had its own *labelled* and *unlabelled* genes: figure 3 represents the amount of genes within each background and their belonging to the KEGG pathways.

The exact numbers are found in the following snippet, which also includes the proportion of KEGG pathways that could be observed in both arrays. The size of *ALL* exceeded that of *Lym* by more than two-fold and was therefore expected to outperform it.

```
##   array n_labelled n_unlabelled n_labelled_kegg n_unlabelled_kegg
## 1   Lym       2482         6507            1586              2953
## 2   ALL       5921         3068            3479              1060
##   prop_labelled_kegg
## 1          0.3494162
## 2          0.7664684
```

Finally, we show the overlap between the KEGG pathways and the *obs* and *Lym* arrays. The table below counts the number of genes lying in the intersections. Most of the genes of the smaller array *Lym* are part of *ALL* as well.

```
##      kegg  ALL  Lym
## kegg 4539 3479 1586
## ALL  3479 5921 2006
## Lym  1586 2006 2482
```

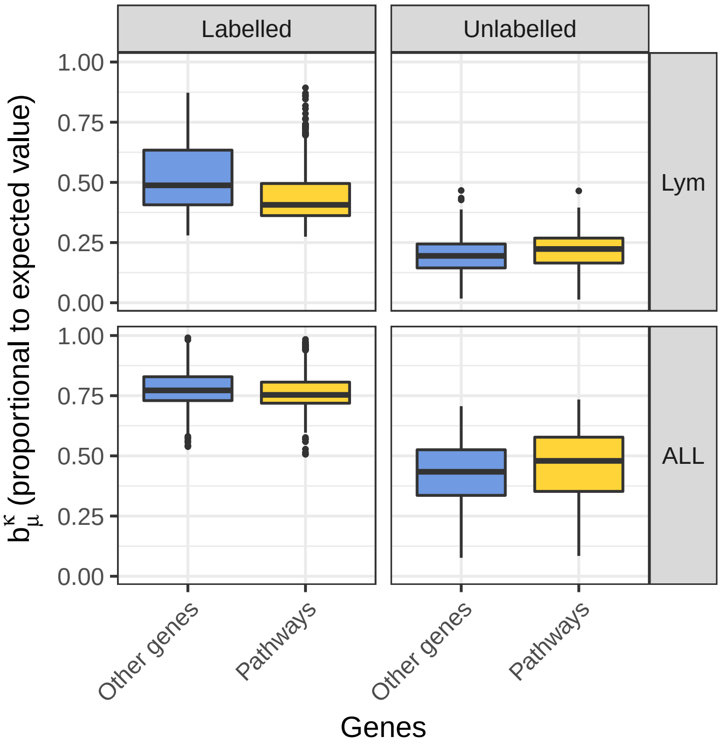

Figure 4: Expected values inside and outside pathways, stratified by observability and array

#### 2.4 Theoretical bias in diffusion scores

As exposed in the main body, the diffusion scores are expected to be biased in terms of their expected value for each node under input permutations. According to the definitions therein, the expected value of a node $i$ is proportional to its reference expected value $b_\mu^\mathcal{K}(i)$. Figure 4 depicts this magnitude, stratified by pathway membership, observability and array.

The following claims were statistically significant in both arrays (Wilcoxon rank-sum test):

1. In the *labelled* genes, pathway genes had a **lower** reference expected value than non-pathway genes.
2. In the *unlabelled* genes, pathway genes had a **higher** reference expected value than non-pathway genes.
3. *Labelled* genes had a **higher** reference expected value than *unlabelled* genes.

Claims 1 and 2:

```
##    obs_label array difference_medians pvalue_wilcox          fdr
## 1   Labelled   Lym        -0.08107461  2.355835e-44 4.711670e-44
## 2   Labelled   ALL        -0.01861420  6.771854e-17 6.771854e-17
## 3 Unlabelled   Lym         0.02852555  4.061263e-39 8.122525e-39
## 4 Unlabelled   ALL         0.04525165  2.472910e-18 2.472910e-18
```

Claim 3:

```
##   array difference_median_bias pvalue_wilcox fdr
## 1   Lym              0.2262074             0   0
## 2   ALL              0.3119931             0   0
```

As every pathway gene was a potential positive, in general terms `raw` should benefit from the bias in (2) and `z` from that in (1). As for *overall* performance (3), `z` equalises *labelled* and *unlabelled* nodes, mixing high and low-confidence predictions. Reliable predictions from the *labelled* part should be masked by those in the *unlabelled* part and the *overall* performance is expected to decrease.

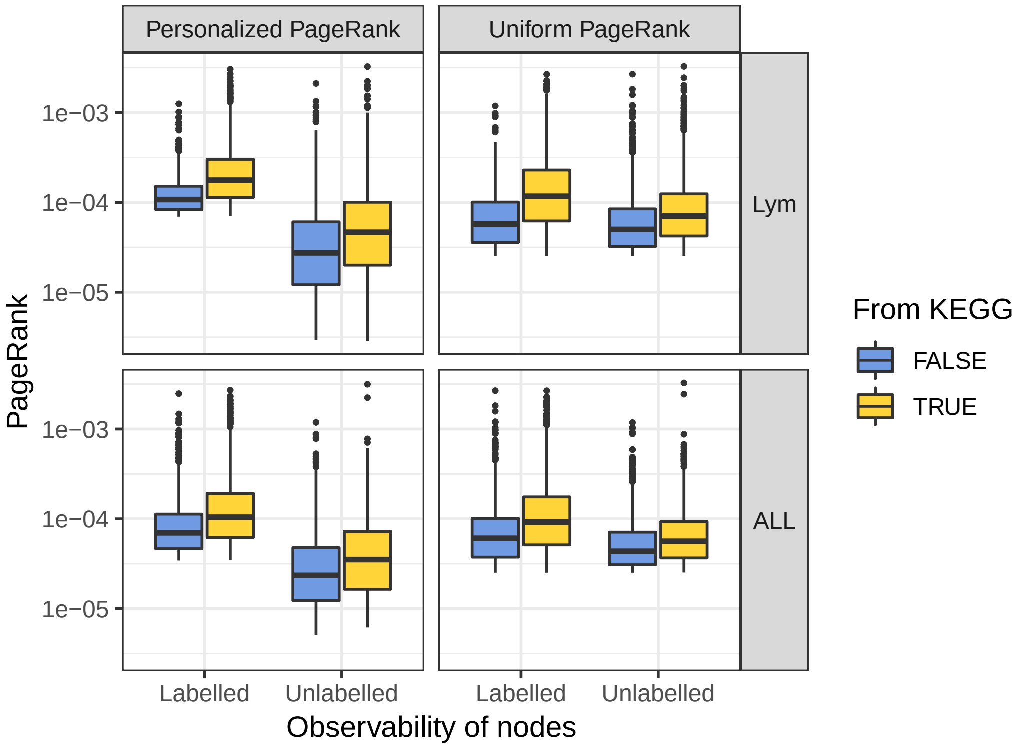

Figure 5: PageRank centralities inside and outside pathways, stratified by observability and array

An important difference exists between indirect bias measurements and the direct quantification of $b_{\mu}^{\mathcal{K}}$. The claims above would be different if PageRank was used as a measure of centrality, under the hypothesis that the bias favours central genes. Figure 5 depicts the PageRank scores (`damping = 0.85`) of all the genes, organised into: both arrays, inside and outside KEGG pathways, labelled and unlabelled nodes. Two PageRank flavours are included: (i) uniform prior and (ii) personalised prior, starting at the labelled genes of each array. In both alternatives, this point of view suggests that `raw` should outperform `z` in the three scenarios, implying that claim (1) would reverse and (2) and (3) would hold.

Genes inside pathways have significantly higher PageRank scores than those outside, in each one of the eight combinations in figure 5:

```
##    obs_label array                 Prior difference_medians pvalue_wilcox
## 1   Labelled   Lym Personalized PageRank       6.906910e-05  2.969662e-76
## 2   Labelled   Lym      Uniform PageRank       5.949779e-05  3.422633e-70
## 3   Labelled   ALL Personalized PageRank       3.437695e-05  3.309309e-90
## 4   Labelled   ALL      Uniform PageRank       3.122078e-05  4.584032e-82
## 5 Unlabelled   Lym Personalized PageRank       1.903473e-05  1.399483e-56
## 6 Unlabelled   Lym      Uniform PageRank       2.034443e-05  8.219762e-58
## 7 Unlabelled   ALL Personalized PageRank       1.174467e-05  8.556411e-19
## 8 Unlabelled   ALL      Uniform PageRank       1.271804e-05  2.362005e-18
##            fdr
## 1 5.939324e-76
## 2 3.422633e-70
## 3 6.618618e-90
## 4 4.584032e-82
## 5 1.399483e-56
## 6 1.643952e-57
## 7 1.711282e-18
```

```
## 8 2.362005e-18
```

#### 2.5 Diffusion inputs

In order to binarise the labels for the diffusion, the false discovery rate, or FDR (Benjamini and Hochberg 1995), of the *labelled* nodes was computed. *Labelled* nodes were defined as positive if their FDR was below 0.1 and negative otherwise. Nodes from the *unlabelled* pool were naturally deemed unlabelled for the diffusion process.

Note that the input could contain false positives due to false positives in hypothesis testing. Likewise, false negatives were expected by the definition of the signal, because only a portion of the genes of the affected pathways will show changes. Occasionally, especially in weak signals (low $r$, $k$ and high $p_{max}$), none of the *labelled* genes would be significant at the specified FDR. Along with other degenerate cases (i.e. no positives in the *unlabelled* nodes), these instances were discarded.

A summary of the metrics table illustrates how the number of instances increased with increasing $r$, $k$ and decreasing $p_{max}$:

```
##  array              strat           method          auroc
##  Lym:61440   Labelled  :40080   raw     :12024   Min.   :0.01272
##  ALL:58800   Unlabelled:40080   ml      :12024   1st Qu.:0.60475
##              Overall   :40080   gm      :12024   Median :0.75456
##                                 ber_s   :12024   Mean   :0.72590
##                                 ber_p   :12024   3rd Qu.:0.86399
##                                 mc      :12024   Max.   :1.00000
##                                 (Other):48096
##      auprc              Column            k           r
##  Min.   :0.0001651   Length:120240      1 :25920   0.3:38160
##  1st Qu.:0.0676635   Class :character   3 :29610   0.5:40650
##  Median :0.1925403   Mode  :character   5 :30900   0.7:41430
##  Mean   :0.2813699                      10:33810
##  3rd Qu.:0.4753768
##  Max.   :1.0000000
##
##    pmax
##  1e-02:15180
##  1e-03:33120
##  1e-04:35940
##  1e-05:36000
##
##
##
```

For methods requiring permutations, the number of permutations was set to 1000 for computational reasons. In all cases, the regularised (unnormalised) Laplacian kernel was used.

### 3 Models

#### 3.1 Model definition

The performance of the diffusion scores in the two arrays under the three signal parameters was best described through explanatory models. Positives in validation were defined as the union of the $k$ pathways that generated each signal. AUROC and AUPRC were computed in three ways: in all the nodes (*overall*), only in the *labelled* part and only in the *unlabelled* part.

Three reference methods were kept. First, `original` ranked the nodes according to their p-value before computing the FDR. In the *labelled* genes, this quantifies the added value of the diffusion process beyond the original signal, i.e. does the diffusion improve the findings obtained by prioritising the genes by their p-value? Regarding the *unlabelled* genes, `original` serves as a reference, as diffusion ignored such p-values by design was not expected to outperform them, especially if $r$ was high or $p_{max}$ was small. Diffusion performance was compared to a hypothetical case in which we knew the original signal – although in general an imperfect one, with false positives and false negatives.

The remaining baselines were `pagerank`, a centrality measure that ignored every input and suggested central genes as top candidates, and `random`, a uniformly random re-ordering of the genes.

The metrics AUROC and AUPRC were modelled through dispersion-adjusted quasibinomial logit models, see `?stats::quasibinomial` in an R console:

$$\text{metric} \sim \text{method} + \text{method:strat} + \text{array} + k + r + p_{max}$$

All the variables were treated as categorical. The interaction term `method:strat` ensured that methods were allowed to have differential performance in the *labelled*, *unlabelled* and *overall* node stratifications. The values $p_{max} = 10^{-5}$ and $k = 10$ were left out due to their respective similarity to $p_{max} = 10^{-4}$ and $k = 5$. Each model is described in its own section.

#### 3.2 AUROC

In this instance, AUROC did not stand out as the ideal metric – details on its model can be found in table 1.

One reason is that, although significant differences existed between methods among *labelled*, *unlabelled* and all nodes, such differences always happened in a narrow range.

More importantly, the performances of diffusion scores (except the ones diffusing $-1$ on the negatives, `ml` and `gm`) were comparable to the `original` p-values in the *unlabelled* genes. The fact that diffusion-based method had no prior data on the *unlabelled* genes should hinder their performance within them, compared to (i) the *labelled* fold, and especially (ii) to the original, unobserved p-values, more notably if $r$ was large. This was not the case, as depicted in figure 6, with predictions by array and partition.

#### 3.3 AUPRC

Contrary to AUROC, AUPRC (see table 2) was more informative for this task.

The quasibinomial model confirmed expected phenomena regarding performance, such as the positive influence of increasing $k$ and $r$ and decreasing $p_{max}$ and the superiority of the *ALL* array. Contrary to AUROC, performance of diffusion scores suffered a pronounced drop in the *unlabelled* genes. Therefore, there was a notable gap in terms of early retrieval between both, something not that apparent from AUROC alone.

Figure 7 shows the expected behaviour of the diffusion scores (actual values in Table 3), in terms of the aforementioned reference expected value $b_{\mu}^{\mathcal{K}}$. As anticipated, `raw` outperformed `z` in the *unlabelled* nodes and *overall*, whereas `z` outperformed `raw` in the *labelled* nodes.

Table 1: Quasilogistic model for AUROC

| | |
|---|---|
| methodml | $-0.393^{***}$ ($-0.432$, $-0.355$) |
| methodgm | $-0.274^{***}$ ($-0.313$, $-0.235$) |
| methodber_s | $-0.135^{***}$ ($-0.175$, $-0.095$) |
| methodber_p | $-0.044^{*}$ ($-0.085$, $-0.004$) |
| methodmc | $-0.199^{***}$ ($-0.238$, $-0.159$) |
| methodz | $-0.162^{***}$ ($-0.201$, $-0.122$) |
| methodoriginal | $-0.750^{***}$ ($-0.787$, $-0.714$) |
| methodpagerank | $-1.059^{***}$ ($-1.095$, $-1.023$) |
| methodrandom | $-1.953^{***}$ ($-1.988$, $-1.918$) |
| k3 | $0.044^{***}$ (0.034, 0.055) |
| k5 | $0.019^{***}$ (0.009, 0.030) |
| r0.5 | $0.237^{***}$ (0.227, 0.247) |
| r0.7 | $0.426^{***}$ (0.415, 0.436) |
| pmax1e-03 | $0.717^{***}$ (0.705, 0.729) |
| pmax1e-04 | $0.797^{***}$ (0.785, 0.809) |
| arrayALL | $0.145^{***}$ (0.137, 0.153) |
| methodraw:stratUnlabelled | $-0.786^{***}$ ($-0.822$, $-0.749$) |
| methodml:stratUnlabelled | $-1.887^{***}$ ($-1.919$, $-1.855$) |
| methodgm:stratUnlabelled | $-1.818^{***}$ ($-1.851$, $-1.785$) |
| methodber_s:stratUnlabelled | $-0.650^{***}$ ($-0.686$, $-0.615$) |
| methodber_p:stratUnlabelled | $-0.700^{***}$ ($-0.736$, $-0.663$) |
| methodmc:stratUnlabelled | $-0.657^{***}$ ($-0.692$, $-0.622$) |
| methodz:stratUnlabelled | $-0.706^{***}$ ($-0.741$, $-0.671$) |
| methodoriginal:stratUnlabelled | $-0.051^{**}$ ($-0.083$, $-0.019$) |
| methodpagerank:stratUnlabelled | $-0.365^{***}$ ($-0.395$, $-0.336$) |
| methodrandom:stratUnlabelled | 0.004 ($-0.024$, 0.031) |
| methodraw:stratOverall | $-0.309^{***}$ ($-0.348$, $-0.270$) |
| methodml:stratOverall | $-1.352^{***}$ ($-1.384$, $-1.320$) |
| methodgm:stratOverall | $-1.270^{***}$ ($-1.303$, $-1.237$) |
| methodber_s:stratOverall | $-0.234^{***}$ ($-0.271$, $-0.196$) |
| methodber_p:stratOverall | $-0.274^{***}$ ($-0.313$, $-0.236$) |
| methodmc:stratOverall | $-0.289^{***}$ ($-0.325$, $-0.252$) |
| methodz:stratOverall | $-0.349^{***}$ ($-0.386$, $-0.313$) |
| methodoriginal:stratOverall | $-0.017$ ($-0.050$, 0.015) |
| methodpagerank:stratOverall | $-0.041^{**}$ ($-0.071$, $-0.011$) |
| methodrandom:stratOverall | 0.001 ($-0.027$, 0.028) |
| Constant | $0.974^{***}$ (0.942, 1.005) |
| Observations | 59,430 |
| *Note:* | $^{*}p<0.05$; $^{**}p<0.01$; $^{***}p<0.001$ |

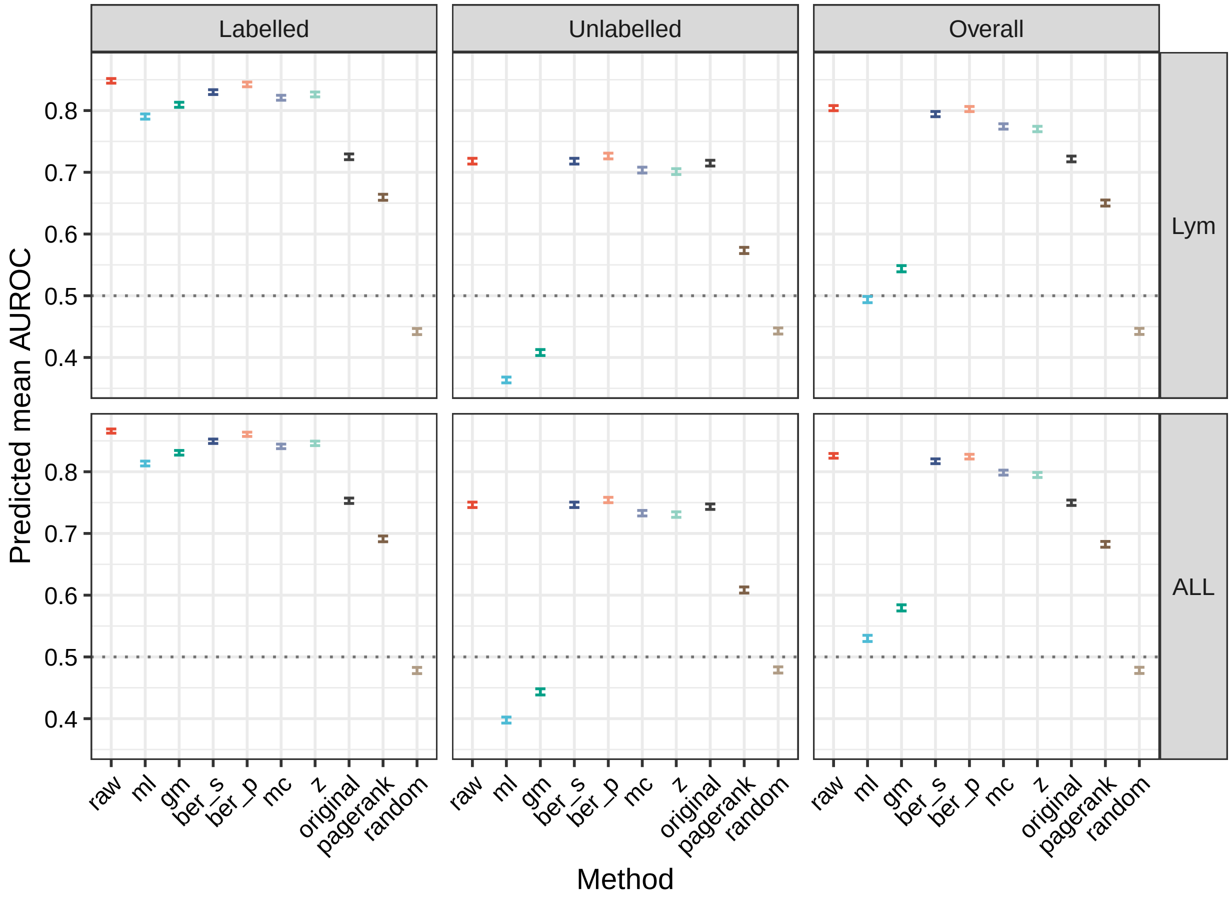

Figure 6: Predictions using the AUROC model (0.95 confidence intervals). Predictions were averaged over the other categorical covariates.

Table 2: Quasilogistic model for AUPRC

| | |
|---|---|
| methodml | $-0.355^{***}$ $(-0.386, -0.324)$ |
| methodgm | $-0.128^{***}$ $(-0.159, -0.097)$ |
| methodber_s | $-0.370^{***}$ $(-0.401, -0.339)$ |
| methodber_p | $0.125^{***}$ $(0.094, 0.156)$ |
| methodmc | $0.236^{***}$ $(0.205, 0.267)$ |
| methodz | $0.461^{***}$ $(0.429, 0.492)$ |
| methodoriginal | $0.261^{***}$ $(0.230, 0.292)$ |
| methodpagerank | $-1.853^{***}$ $(-1.890, -1.815)$ |
| methodrandom | $-3.082^{***}$ $(-3.136, -3.028)$ |
| k3 | $0.447^{***}$ $(0.434, 0.460)$ |
| k5 | $0.618^{***}$ $(0.605, 0.631)$ |
| r0.5 | $0.512^{***}$ $(0.499, 0.525)$ |
| r0.7 | $0.912^{***}$ $(0.899, 0.924)$ |
| pmax1e-03 | $1.548^{***}$ $(1.528, 1.568)$ |
| pmax1e-04 | $1.672^{***}$ $(1.652, 1.692)$ |
| arrayALL | $0.147^{***}$ $(0.136, 0.157)$ |
| methodraw:stratUnlabelled | $-2.075^{***}$ $(-2.114, -2.035)$ |
| methodml:stratUnlabelled | $-3.424^{***}$ $(-3.496, -3.353)$ |
| methodgm:stratUnlabelled | $-3.001^{***}$ $(-3.056, -2.946)$ |
| methodber_s:stratUnlabelled | $-1.705^{***}$ $(-1.745, -1.665)$ |
| methodber_p:stratUnlabelled | $-2.230^{***}$ $(-2.270, -2.190)$ |
| methodmc:stratUnlabelled | $-2.918^{***}$ $(-2.965, -2.871)$ |
| methodz:stratUnlabelled | $-2.889^{***}$ $(-2.933, -2.845)$ |
| methodoriginal:stratUnlabelled | $-0.338^{***}$ $(-0.369, -0.307)$ |
| methodpagerank:stratUnlabelled | $-1.594^{***}$ $(-1.660, -1.529)$ |
| methodrandom:stratUnlabelled | $-0.969^{***}$ $(-1.061, -0.877)$ |
| methodraw:stratOverall | $-0.564^{***}$ $(-0.595, -0.532)$ |
| methodml:stratOverall | $-2.121^{***}$ $(-2.165, -2.077)$ |
| methodgm:stratOverall | $-1.568^{***}$ $(-1.604, -1.531)$ |
| methodber_s:stratOverall | $-0.528^{***}$ $(-0.561, -0.496)$ |
| methodber_p:stratOverall | $-0.636^{***}$ $(-0.667, -0.604)$ |
| methodmc:stratOverall | $-1.430^{***}$ $(-1.464, -1.397)$ |
| methodz:stratOverall | $-1.500^{***}$ $(-1.533, -1.467)$ |
| methodoriginal:stratOverall | $-0.142^{***}$ $(-0.173, -0.111)$ |
| methodpagerank:stratOverall | $-0.486^{***}$ $(-0.533, -0.438)$ |
| methodrandom:stratOverall | $-0.462^{***}$ $(-0.541, -0.384)$ |
| Constant | $-2.434^{***}$ $(-2.465, -2.403)$ |
| Observations | 59,430 |
| *Note:* | $^{*}$p<0.05; $^{**}$p<0.01; $^{***}$p<0.001 |

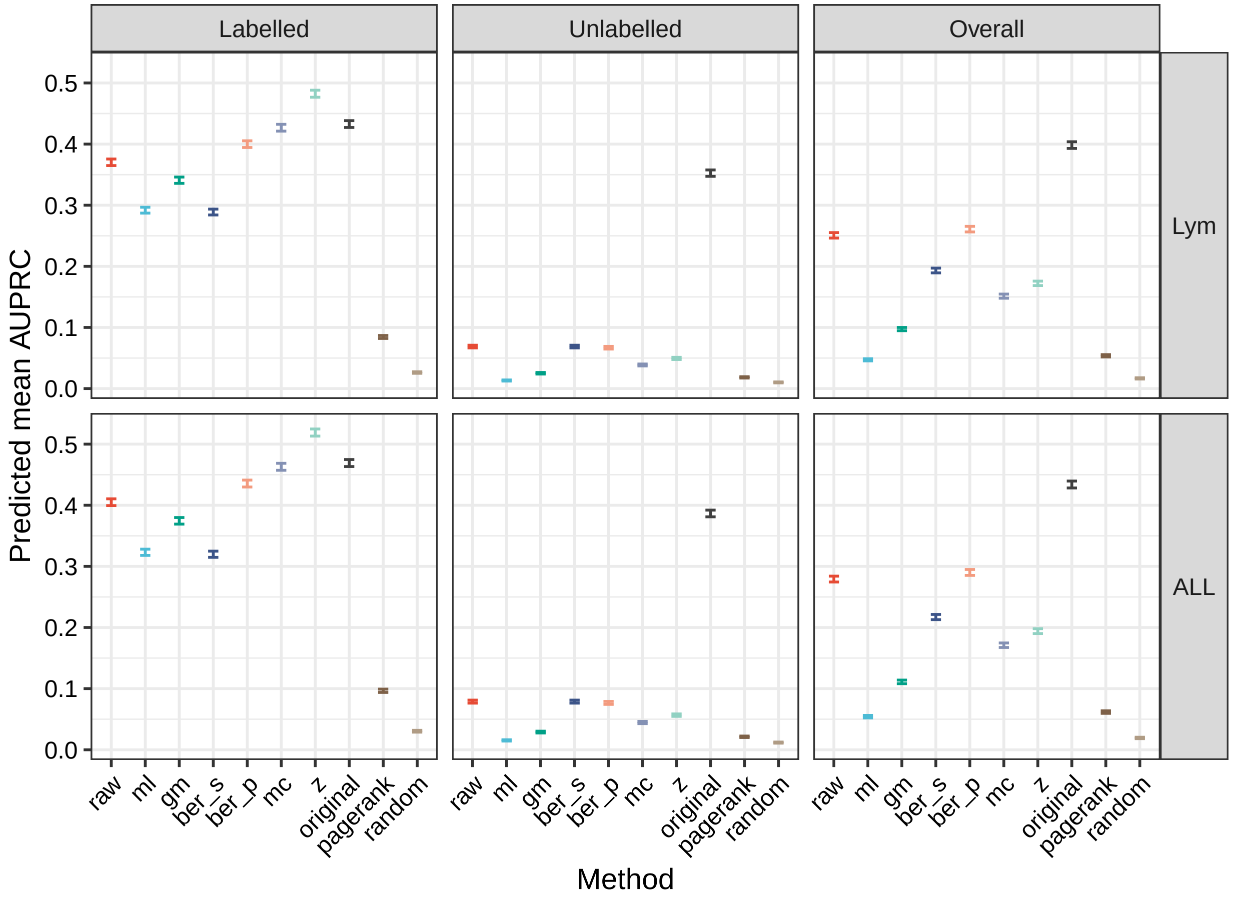

Figure 7: Predictions using the AUPRC model (0.95 confidence intervals). Predictions were averaged over the other categorical covariates.

| array | method | Labelled | Unlabelled | Overall |
|---|---|---|---|---|
| Lym | raw | (0.365, 0.376) | (0.067, 0.071) | (0.246, 0.255) |
| Lym | ml | (0.287, 0.297) | (0.012, 0.014) | (0.045, 0.049) |
| Lym | gm | (0.336, 0.346) | (0.024, 0.026) | (0.095, 0.100) |
| Lym | ber_s | (0.284, 0.294) | (0.067, 0.071) | (0.189, 0.197) |
| Lym | ber_p | (0.394, 0.405) | (0.065, 0.069) | (0.256, 0.265) |
| Lym | mc | (0.421, 0.432) | (0.037, 0.040) | (0.148, 0.155) |
| Lym | z | (0.477, 0.488) | (0.048, 0.051) | (0.169, 0.176) |
| Lym | original | (0.427, 0.438) | (0.347, 0.358) | (0.393, 0.404) |
| Lym | pagerank | (0.082, 0.087) | (0.017, 0.019) | (0.052, 0.056) |
| Lym | random | (0.025, 0.028) | (0.009, 0.011) | (0.016, 0.018) |
| ALL | raw | (0.399, 0.411) | (0.076, 0.081) | (0.274, 0.284) |
| ALL | ml | (0.318, 0.328) | (0.014, 0.016) | (0.052, 0.056) |
| ALL | gm | (0.369, 0.380) | (0.028, 0.030) | (0.108, 0.114) |
| ALL | ber_s | (0.315, 0.325) | (0.076, 0.081) | (0.213, 0.221) |
| ALL | ber_p | (0.430, 0.441) | (0.074, 0.079) | (0.285, 0.295) |
| ALL | mc | (0.457, 0.469) | (0.043, 0.046) | (0.167, 0.175) |
| ALL | z | (0.513, 0.525) | (0.055, 0.059) | (0.190, 0.198) |
| ALL | original | (0.463, 0.475) | (0.381, 0.392) | (0.428, 0.440) |
| ALL | pagerank | (0.094, 0.099) | (0.020, 0.022) | (0.060, 0.064) |
| ALL | random | (0.029, 0.032) | (0.011, 0.013) | (0.018, 0.020) |

Table 3: Confidence intervals (0.95) on predicted AUPRC, averaged over covariates.

Below are the results of the statistical test between `raw` and `z` that back up the claims in this section.

```
##                               contrast odds.ratio p.value
## 1     raw,Labelled,Lym / z,Labelled,Lym  0.6308394       0
## 2 raw,Unlabelled,Lym / z,Unlabelled,Lym  1.4240117       0
## 3       raw,Overall,Lym / z,Overall,Lym  1.6090716       0
## 4     raw,Labelled,ALL / z,Labelled,ALL  0.6308394       0
## 5 raw,Unlabelled,ALL / z,Unlabelled,ALL  1.4240117       0
## 6       raw,Overall,ALL / z,Overall,ALL  1.6090716       0
```

##### 3.4 Other remarks

- Using an indirect measure of bias might be misleading. Here, by using PageRank as a centrality measure and assuming that `raw` scores will favour highly connected nodes, we would expect that `raw` outperforms `z` in the *labelled* nodes. However, this is inded the opposite to what $b_\mu^\mathcal{K}$ (a direct quantification of the expected value-related bias) suggests and to what we observe in terms of performance.
- The `original` baseline was difficult to improve upon, even in the *labelled* genes, in terms of AUPRC. This was not the case for AUROC, implying that although diffusion had a positive and noticeable impact in the overall ranking, early retrieval was a challenging task.
- `ber_p` had the best *overall* performance, suggesting that a consensus between normalised and unnormalised scores can be beneficial.
- `ml` and `gm` suffered from this imbalanced datasets, where positives were vastly outnumbered by negatives.
- Within normalised scores, `z` outperformed `mc`, possibly due to the presence of ties and the stochastic nature of the latter.

### 4 Reproducibility

```
##  [1] "R version 3.5.3 (2019-03-11)"
##  [2] "Platform: x86_64-pc-linux-gnu (64-bit)"
##  [3] "Running under: Ubuntu 16.04.6 LTS"
##  [4] ""
##  [5] "Matrix products: default"
##  [6] "BLAS: /usr/lib/atlas-base/atlas/libblas.so.3.0"
##  [7] "LAPACK: /usr/lib/atlas-base/atlas/liblapack.so.3.0"
##  [8] ""
##  [9] "locale:"
## [10] " [1] LC_CTYPE=en_US.UTF-8       LC_NUMERIC=C              "
## [11] " [3] LC_TIME=en_US.UTF-8        LC_COLLATE=en_US.UTF-8    "
## [12] " [5] LC_MONETARY=en_US.UTF-8    LC_MESSAGES=en_US.UTF-8   "
## [13] " [7] LC_PAPER=en_US.UTF-8       LC_NAME=C                 "
## [14] " [9] LC_ADDRESS=C               LC_TELEPHONE=C            "
## [15] "[11] LC_MEASUREMENT=en_US.UTF-8 LC_IDENTIFICATION=C       "
## [16] ""
## [17] "attached base packages:"
## [18] "[1] grid      stats     graphics  grDevices utils     datasets  methods  "
## [19] "[8] base     "
## [20] ""
## [21] "other attached packages:"
## [22] " [1] bindrcpp_0.2.2    xtable_1.8-3      data.table_1.11.8"
## [23] " [4] extrafont_0.17    gtable_0.2.0      ggsci_2.9        "
## [24] " [7] ggplot2_3.1.0     stargazer_5.2.2   emmeans_1.3.0    "
## [25] "[10] magrittr_1.5      tidyr_0.8.2       dplyr_0.7.8      "
## [26] "[13] plyr_1.8.4        igraph_1.2.2     "
## [27] ""
## [28] "loaded via a namespace (and not attached):"
## [29] " [1] zoo_1.8-4                 tidyselect_0.2.5         "
## [30] " [3] xfun_0.4                  reshape2_1.4.3           "
## [31] " [5] purrr_0.2.5               splines_3.5.3            "
## [32] " [7] lattice_0.20-38           expm_0.999-3             "
## [33] " [9] colorspace_1.3-2          htmltools_0.3.6          "
## [34] "[11] yaml_2.2.0                survival_2.43-3          "
## [35] "[13] rlang_0.3.0.1             pillar_1.3.0             "
## [36] "[15] glue_1.3.0                withr_2.1.2              "
## [37] "[17] multcomp_1.4-8            bindr_0.1.1              "
## [38] "[19] stringr_1.3.1             munsell_0.5.0            "
## [39] "[21] mvtnorm_1.0-8             codetools_0.2-16         "
## [40] "[23] evaluate_0.12             labeling_0.3             "
## [41] "[25] RcppArmadillo_0.9.200.4.0 knitr_1.20               "
## [42] "[27] Rttf2pt1_1.3.7            TH.data_1.0-9            "
## [43] "[29] Rcpp_1.0.0                backports_1.1.2          "
## [44] "[31] scales_1.0.0              RcppParallel_4.4.1       "
## [45] "[33] precrec_0.9.1             digest_0.6.18            "
## [46] "[35] stringi_1.2.4             bookdown_0.7             "
## [47] "[37] rprojroot_1.3-2           tools_3.5.3              "
## [48] "[39] diffuStats_1.2.0          sandwich_2.5-0           "
## [49] "[41] lazyeval_0.2.1            tibble_1.4.2             "
## [50] "[43] crayon_1.3.4              extrafontdb_1.0          "
## [51] "[45] pkgconfig_2.0.2           MASS_7.3-51.1            "
## [52] "[47] Matrix_1.2-15             estimability_1.3         "
```

```
## [53] "[49] assertthat_0.2.0          rmarkdown_1.10            "
## [54] "[51] R6_2.3.0                  compiler_3.5.3            "
```

Kanehisa, Minoru, Miho Furumichi, Mao Tanabe, Yoko Sato, and Kanae Morishima. 2017. "KEGG: New Perspectives on Genomes, Pathways, Diseases and Drugs." *Nucleic Acids Research* 45 (D1). Oxford University Press: D353–D361.

Li, Xiaochun. 2009. *ALL: A Data Package*.

Mishra, Gopa R, M Suresh, K Kumaran, N Kannabiran, Shubha Suresh, P Bala, K Shivakumar, et al. 2006. "Human Protein Reference Database—2006 Update." *Nucleic Acids Research* 34 (suppl_1). Oxford University Press: D411–D414.

Rosenwald, Andreas, George Wright, Wing C Chan, Joseph M Connors, Elias Campo, Richard I Fisher, Randy D Gascoyne, et al. 2002. "The Use of Molecular Profiling to Predict Survival After Chemotherapy for Diffuse Large-B-Cell Lymphoma." *New England Journal of Medicine* 346 (25). Mass Medical Soc: 1937–47.
