## Supplement 4 for "The effect of statistical normalisation on network propagation scores"

*October 18, 2018*

**4 Reproducibility** **17**

**References** **19**

### 1 Introduction

This additional file contains details on the prospective pathway prediction case study. A protein-protein interaction network and biological pathways, both from year 2011, were used to predict new genes in the same pathways from 2018.
This document can be re-built anytime by knitting its corresponding `.Rmd` file.

#### 1.1 The network

We used the BioGRID network (Chatr-aryamontri et al. 2017), weighting its interactions according to (Cao et al. 2014). Weights depend on the amount of experiments reporting an interaction and their throughput, favouring low-throughput methodologies.
In addition, in order to avoid circularity between the new pathway genes and the network construction, the network was restricted to interactions from publications in 2010 or older. This posed a realistic prospective scenario, in which the network might not consistently reflect the novel biology behing the newly added genes.

Below is a summary of the network:

```
## IGRAPH 5fd82d3 UNW- 11394 67573 --
## + attr: name (v/c), weight (e/n)
```

The network contained 11394 nodes and 67573 edges and was connected by construction (only the largest connected component was kept). The edges weights are displayed in figure 1, revealing two broad categories: low-confidence ones, with a weight of 0.25, and high-confidence ones, with a weight of 0.8 or higher.

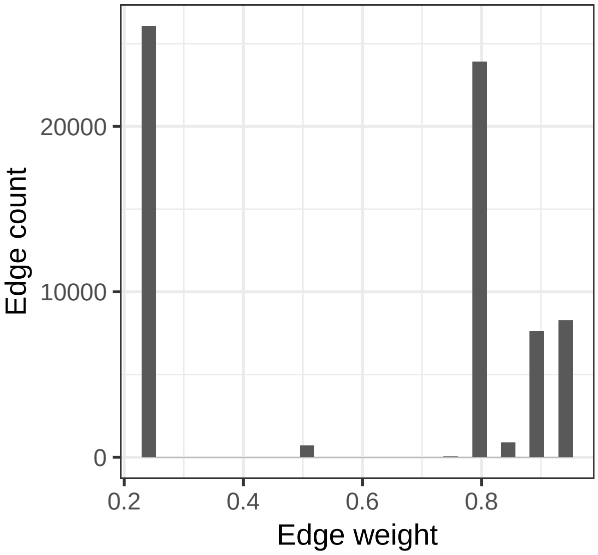

Figure 1: Distribution of the edge weights in the BioGRID network.

### 2 Descriptive statistics

#### 2.1 KEGG pathways

The Kyoto Encyclopedia of Genes and Genomes (KEGG) pathways (Kanehisa et al. 2017) was used as input and validation for the diffusion scores. Pathways were treated as gene sets, only relying on the network data from BioGRID.

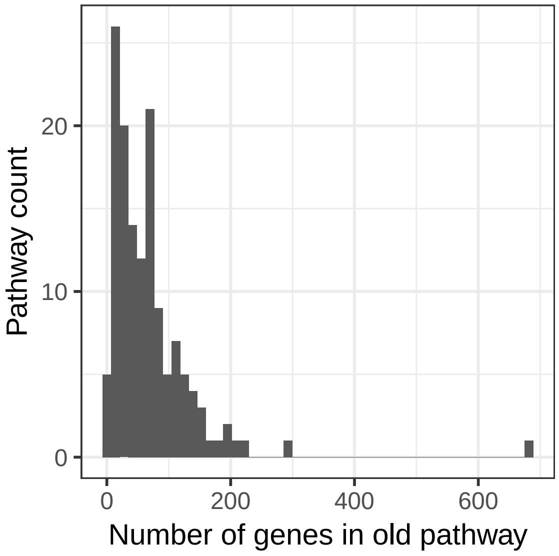

Figure 2: Number of genes per pathway in the older KEGG release

An older version of the pathways, dating from 2011, was used to predict new pathway genes in 2018. The last public version of KEGG, dated from March 14th 2011, was obtained from the `KEGG.db` package (Carlson 2016). Likewise, a more recent KEGG release was downloaded in August 18th, 2018 from `https://www.kegg.jp/kegg/rest/keggapi.html`.

A total of 139 KEGG pathways had at least one additional gene in the latest version, after mapping the genes to the BioGRID network. Figure 2 shows that most pathways contained up to 200 genes, while figure 3 depicts how they typically involved less than 20 new genes. Likewise, figure 4 describes how ubiquitous new genes were: most of the new genes belonged to a single pathway.

#### 2.2 Theoretical bias in diffusion scores

In this occasion, the inherent bias of the diffusion scores was not related to the expected value of each node under input permutations. Given the present setup, where all the nodes were considered as *labelled*, $b_{\mu}^{\mathcal{K}}$ is constant and thus the `raw` scores must have a constant expected value on all the nodes (see proofs on properties of diffusion scores from Supplement 1). However, differences existed in terms of **variance**. We hypothesised that this led to a variance-related bias, where some nodes would exhibit more stable diffusion scores whereas others could greatly vary under input permutations. Specifically, we hypothesised that `z` would improve the power on low-variance nodes. Variance-related bias was quantified through ther reference variance $b_{\sigma^2}^{\mathcal{K}}$, defined in the main text as proportional to the logarithm of the node variance.

Before framing the genes into pathways, figure 5 suggests that the variance was mainly driven by the node degree. The diffusion scores of highly connected nodes were therefore expected to be less sensitive to perturbations in the input.

Figure 6 depicts the reference variance $b_{\sigma^2}^{\mathcal{K}}$, dividing genes into four categories: *old* for the genes in the old and new pathway, *new* for the genes only in the new pathway, *old_fp* for the genes only in the old pathway and *other* for the rest of genes. Note that a gene can belong to several categories, i.e. *new* for one pathway and *other* for another. Figure 6 suggests that the properties of *old*, *new* and *other* genes are essentially different and linked to their topological properties.

The same magnitude was depicted in terms of pathways, representing the median value of $b_{\sigma^2}^{\mathcal{K}}(i)$ for its *new* genes, see figure 7. The plot suggest that the *new* genes can have two sorts of biases, specifically a standard deviation either (i) lower or (ii) higher than that of the *other* network genes in general.

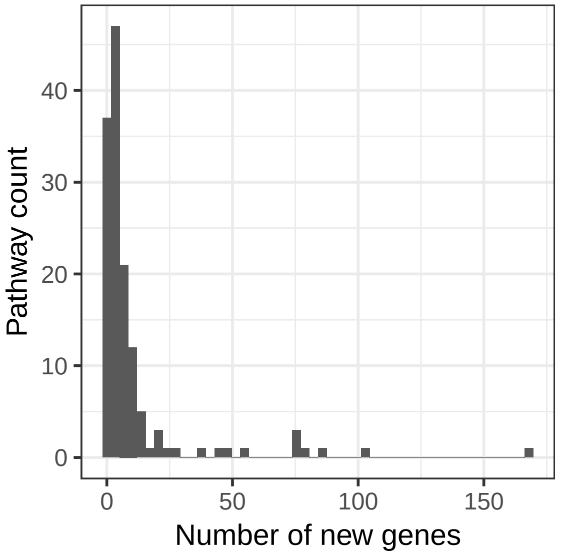

Figure 3: Number of new genes per pathway in the latest KEGG release

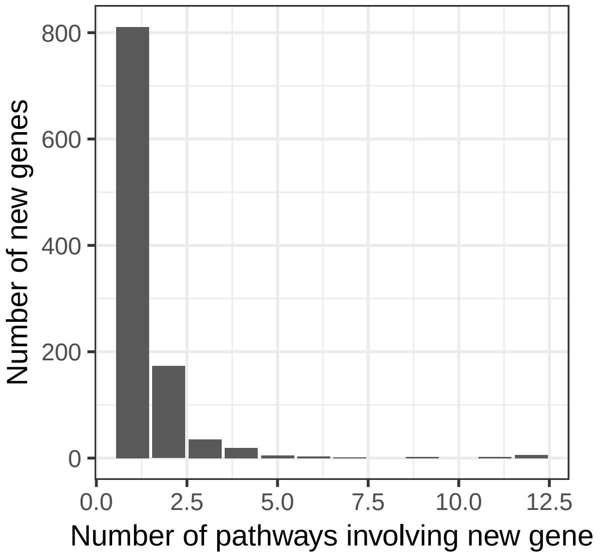

Figure 4: Number of pathways involving each new gene

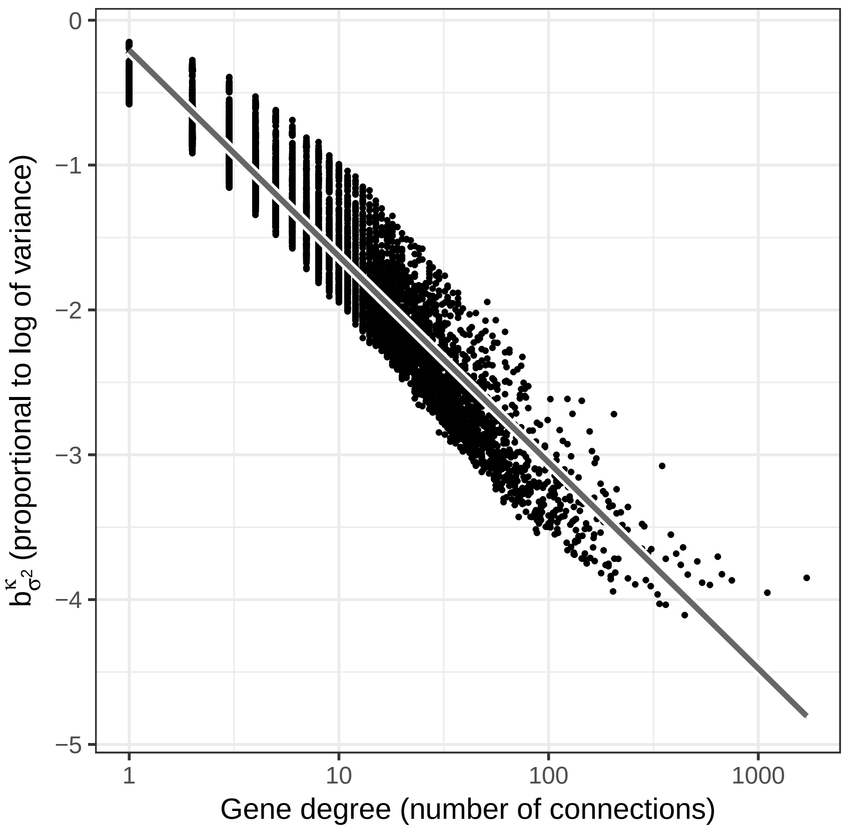

Figure 5: Variance-related bias across all the genes in terms of degree. The gray line shows the best linear fit.

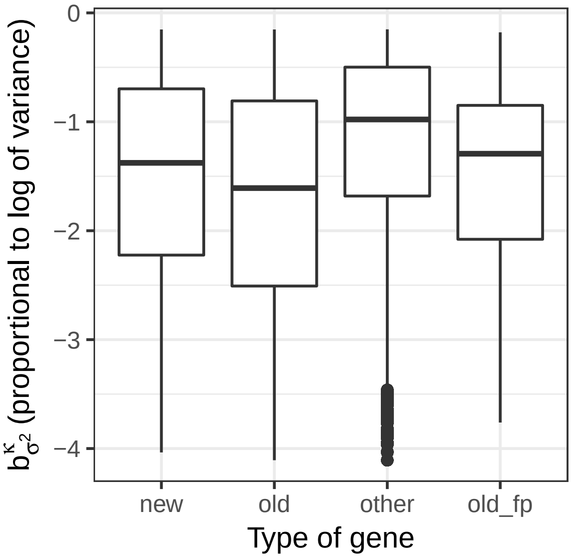

Figure 6: Variance-related bias across all the genes. Each unique gene appears exactly once for every pathway.

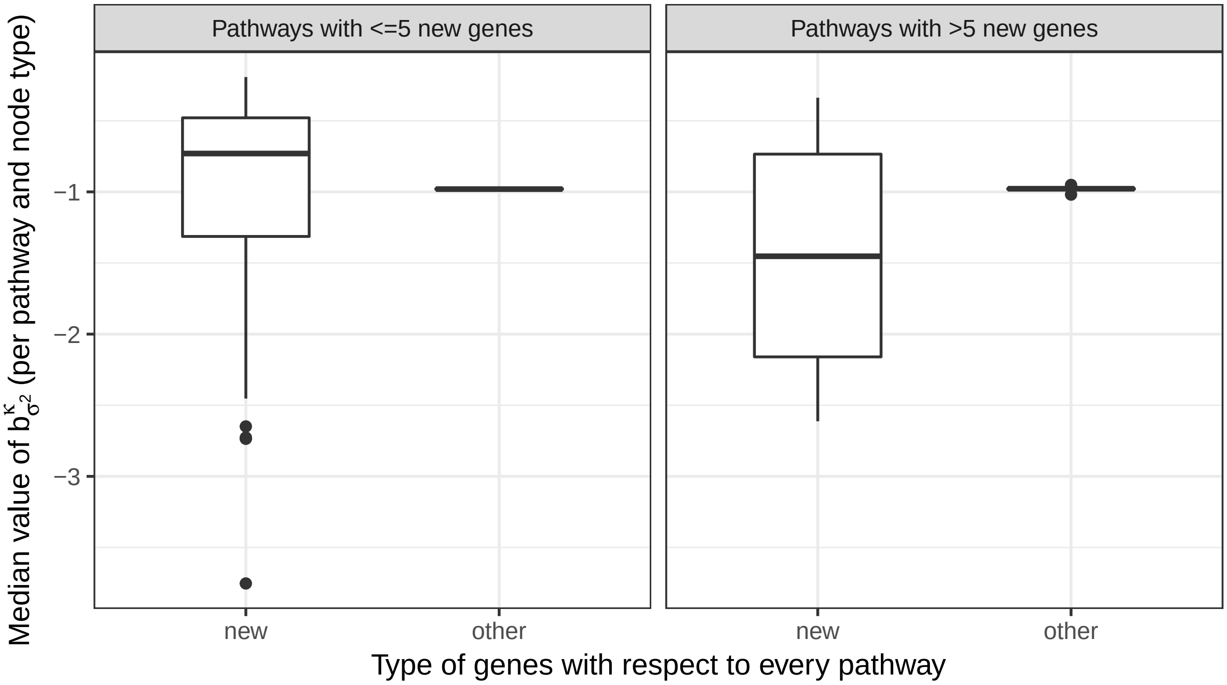

Figure 7: Variance-related bias across all the pathways. The median reference variance of the new and the other genes for each pathway is represented, leading to two data points per pathway. Pathways were divided in two groups according to their number of new genes.

Differences in $b^{\mathcal{K}}_{\sigma^2}(i)$ between *new* and *other* genes were tested using `wilcox.test` and correcting for False Discovery Rate (FDR) (Benjamini and Hochberg 1995), see figure 8. Differences at FDR $< 0.1$ could be proven for pathways some pathways, almost always with more than 5 *new* genes. Significant differences were usually negative (i.e. *other* genes having a greater median, in line with figure 7), but positive differences existed too.

#### 2.3 Diffusion inputs

As the pathways were treated as gene sets, inputs were naturally defined as binary labels without further modification. Note that pathways could contain genes that were present in the old release but dropped in the last one, acting as a *false positive*. In total, 139 instances (one per pathway) were defined and genes outside the original pathway were ranked. Afterwards, the AUROC and AUPRC metrics were computed and compared through explanatory models.

#### 2.4 Diffusion scores and bias

Before diving into pathway-wise performance metrics, diffusion scores `raw` and `z` were compared in views of the variance-related bias. Figure 9 sheds light on the expected behaviour of the statistical normalisation and supports the hypothesis that normalising the scores helps decorrelate power from the reference variance values. The actual impact on overall performance still depends on other factors, such as the density of positives throughout the reference variances.

As for method parameters, the regularised (unnormalised) Laplacian kernel was used and permutation-based scores used $10^4$ random trials. Methods `ml`, `gm` and `ber_s` were excluded from this comparison because their ranking is identical to that of `raw` in the current settings, see the diffusion scores equivalence properties 1

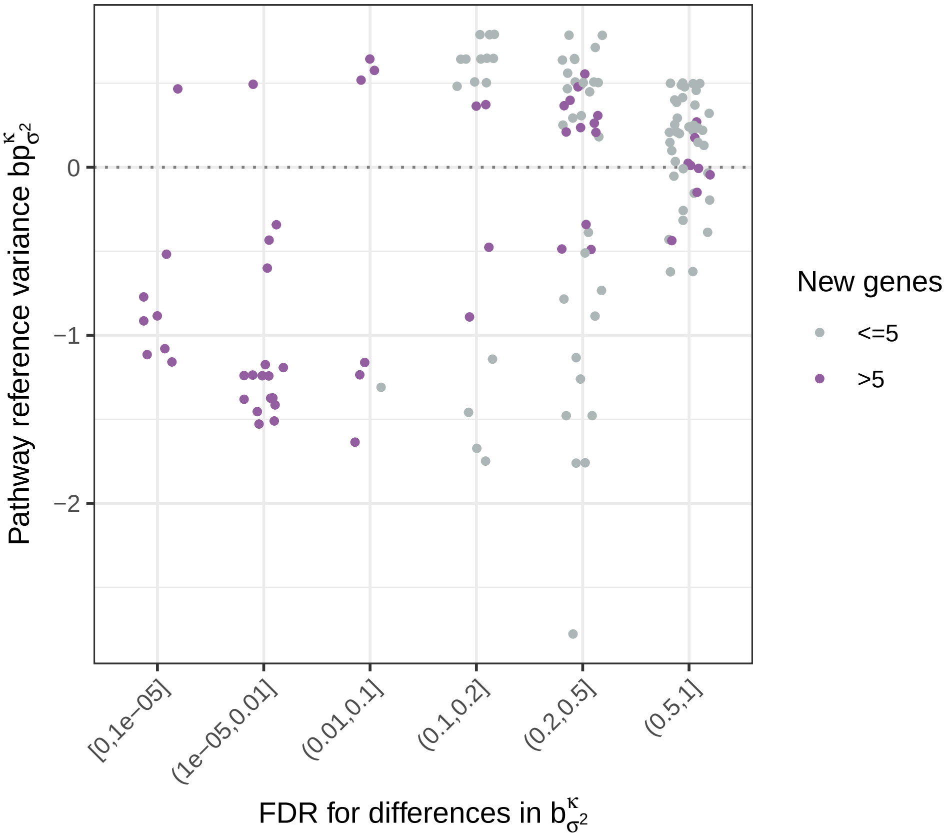

Figure 8: Statistical differences on the reference variances of new versus other genes in each pathway. Each data point represents a pathway. Differences were stratified by the amount of new genes, which affects the statistical power to spot differences.

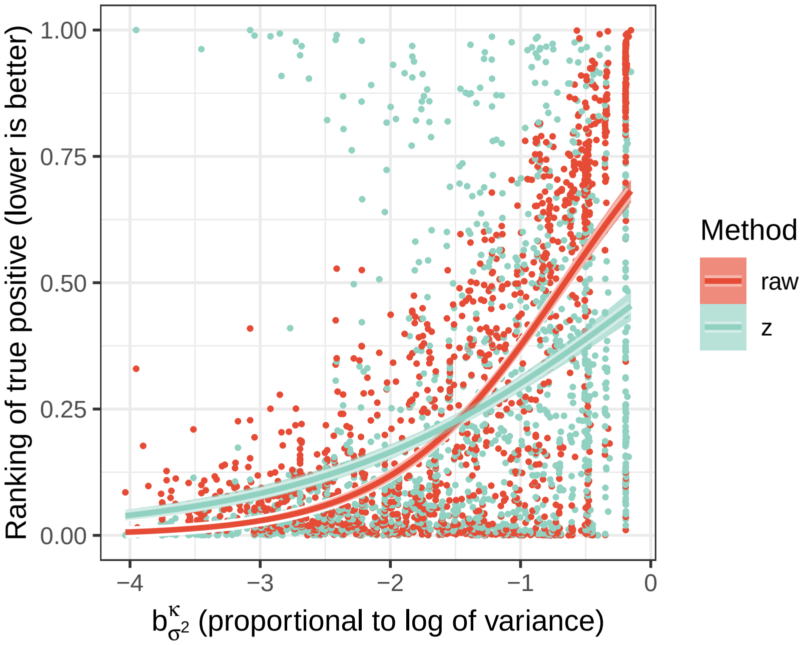

Figure 9: Ranking of true positives as a function of the variance-related bias; lines correspond to a logistic fit with 0.95 confidence intervals. This plot represents the union of the positives of each pathway and their relative ranking in their prioritisation. Nodes closer to 0 were top ranked for that specific pathway, and therefore well prioritised, whereas worst ranked nodes were close to 1. The unnormalised scores raw had more power on nodes with lower standard deviation, at the cost of being less sensitive among larger standard deviations. The normalised scores z showed a more bias-independent power, at the cost of missing positives with smaller standard deviations.

(`ml`, `gm`) and 3 (`ber_s`) in Supplement 1. Two baselines were considered: `pagerank` (with `damping = 0.85`), which tends to suggest central genes regardless of the input, and `random` (random prioritisation).

### 3 Models

#### 3.1 Model definition

The metrics AUROC and AUPRC were modelled through dispersion-adjusted quasibinomial logit models, see `?stats::quasibinomial` in an R console:

$$\text{metric} \sim \text{method} + \text{method:path\_var\_ref}$$

The categorical variable `method` could be `raw`, `ber_p`, `mc`, `z` or the baselines `pagerank` and `random`. The term `path_var_ref` was a pathway property, computed as the difference between the median of the reference variance $b^{\mathcal{K}}_{\sigma^2}(i)$ for the *new* genes in the pathway, and the median of $b^{\mathcal{K}}_{\sigma^2}(i)$ for the *other* genes, as depicted in figure 8. `path_var_ref` intended to summarise the bias of a whole pathway in a single number: positive (negative) values indicated that the *new* genes had more (less) variance than the average gene in the network. In order to test our hypothesis, the interaction term `method:path_var_ref` allowed methods to be affected in different ways by the pathway-wise bias.

#### 3.2 AUROC

Table 1 summarises the AUROC model. As this case study was not simulated, the number of data points was limited due to the prospective design, being notably lower than that of the other datasets.

Table 1: Quasibinomial model for AUROC

| | |
|---|---|
| methodber_p | 0.271** (0.057, 0.486) |
| methodmc | 0.291*** (0.079, 0.503) |
| methodz | 0.560*** (0.342, 0.779) |
| methodpagerank | −0.891*** (−1.100, −0.681) |
| methodrandom | −0.558*** (−0.758, −0.358) |
| methodraw:path_var_ref | −1.387*** (−1.648, −1.127) |
| methodber_p:path_var_ref | −1.030*** (−1.279, −0.782) |
| methodmc:path_var_ref | −0.635*** (−0.854, −0.417) |
| methodz:path_var_ref | −0.484*** (−0.710, −0.258) |
| methodpagerank:path_var_ref | −1.473*** (−1.695, −1.251) |
| methodrandom:path_var_ref | 0.035 (−0.129, 0.199) |
| Constant | 0.710*** (0.559, 0.861) |
| Observations | 834 |
| *Note:* | *p<0.1; **p<0.05; ***p<0.01 |

The model in table 1 supported the claim that `raw` was more affected than `z` by the reference variance. Figure 10 reflects this fact along the values of `path_var_ref`, whereas the contrast between the interaction terms (i.e. of the form `method:path_var_ref`) of `raw` and `z` was significant:

```
##  contrast             estimate        SE  df z.ratio p.value
##  raw - ber_p       -0.35722993 0.1837676 Inf  -1.944  0.3752
##  raw - mc          -0.75203314 0.1735438 Inf  -4.333  0.0002
##  raw - z           -0.90326861 0.1761406 Inf  -5.128  <.0001
##  raw - pagerank     0.08536864 0.1747622 Inf   0.488  0.9966
##  raw - random      -1.42200492 0.1570979 Inf  -9.052  <.0001
##  ber_p - mc        -0.39480321 0.1688678 Inf  -2.338  0.1789
##  ber_p - z         -0.54603868 0.1715353 Inf  -3.183  0.0182
##  ber_p - pagerank   0.44259857 0.1701197 Inf   2.602  0.0968
##  ber_p - random    -1.06477498 0.1519165 Inf  -7.009  <.0001
##  mc - z            -0.15123547 0.1605344 Inf  -0.942  0.9356
##  mc - pagerank      0.83740178 0.1590209 Inf   5.266  <.0001
##  mc - random       -0.66997178 0.1393756 Inf  -4.807  <.0001
##  z - pagerank       0.98863725 0.1618508 Inf   6.108  <.0001
##  z - random        -0.51873631 0.1425959 Inf  -3.638  0.0037
##  pagerank - random -1.50737356 0.1408898 Inf -10.699  <.0001
##
## P value adjustment: tukey method for comparing a family of 6 estimates
```

Predictions with confidence intervals in the mean value of `path_var_ref` are shown in figure 11, whereas their raw values can be found in figure 12 – both figures depict similar trends. Testing overall differences (averaging over `path_var_ref` and using Tukey's test), `z` significantly outperformed `raw`:

```
##  contrast          odds.ratio         SE  df z.ratio p.value
##  raw / ber_p        0.8158250 0.09847996 Inf  -1.686  0.5409
##  raw / mc           0.8623415 0.10037696 Inf  -1.272  0.8002
##  raw / z            0.6781553 0.08086040 Inf  -3.257  0.0143
##  raw / pagerank     2.3974532 0.27356640 Inf   7.663  <.0001
```

| | raw | ber_p | mc | z | pagerank | random |
|---|---|---|---|---|---|---|
| raw | | -0.038(-0.06,-0.02) | -0.043(-0.077,-0.016) | -0.084(-0.13,-0.045) | 0.16(0.13,0.19) | 0.16(0.1,0.21) |
| ber_p | 3.49e-07 | | -0.0011(-0.01,0.0058) | -0.026(-0.046,-0.011) | 0.22(0.18,0.26) | 0.22(0.17,0.26) |
| mc | 4.74e-04 | 7.73e-01 | | -0.035(-0.053,-0.02) | 0.23(0.19,0.28) | 0.22(0.18,0.26) |
| z | 5.39e-09 | 5.16e-04 | 1.01e-05 | | 0.29(0.24,0.33) | 0.26(0.22,0.3) |
| pagerank | 1.72e-17 | 7.14e-19 | 3.91e-17 | 7.14e-19 | | -0.026(-0.083,0.034) |
| random | 6.66e-07 | 1.96e-11 | 8.36e-13 | 1.71e-17 | 4.40e-01 | |

Table 2: Paired two-sided Wilcoxon test between AUROCs, corrected by FDR. Above diagonal: differences with 0.95 confidence interval. Below diagonal: FDR.

```
##  raw / random       2.2891461 0.24679547 Inf   7.682  <.0001
##  ber_p / mc         1.0570178 0.12225956 Inf   0.479  0.9969
##  ber_p / z          0.8312510 0.09851785 Inf  -1.559  0.6254
##  ber_p / pagerank   2.9386857 0.33311868 Inf   9.510  <.0001
##  ber_p / random     2.8059279 0.30027989 Inf   9.641  <.0001
##  mc / z             0.7864116 0.08974768 Inf  -2.105  0.2843
##  mc / pagerank      2.7801668 0.30235276 Inf   9.402  <.0001
##  mc / random        2.6545702 0.27110599 Inf   9.559  <.0001
##  z / pagerank       3.5352568 0.39518139 Inf  11.297  <.0001
##  z / random         3.3755483 0.35560792 Inf  11.548  <.0001
##  pagerank / random  0.9548241 0.09501094 Inf  -0.465  0.9973
##
## P value adjustment: tukey method for comparing a family of 6 estimates
## Tests are performed on the log odds ratio scale
```

A paired non-parametric test outside the model yielded stronger evidence of such differences, see table 2.

Note how by its own definition, the `pagerank` centrality baseline was noticeably affected by the bias, in a similar way to the `raw` scores (figure 10). This was expected because node degree, the most basic measure of centrality, showed collinearity with the reference variance (figure 5). Provided that pathway biases were found in both directions, i.e. genes with either more or less variance than most genes (figure 8), `pagerank` had a close-to-random AUROC (figure 12). On the other hand, the `random` baseline behaved as expected, with an AUROC close to 0.5 and independent of the reference pathway variance (figure 10).

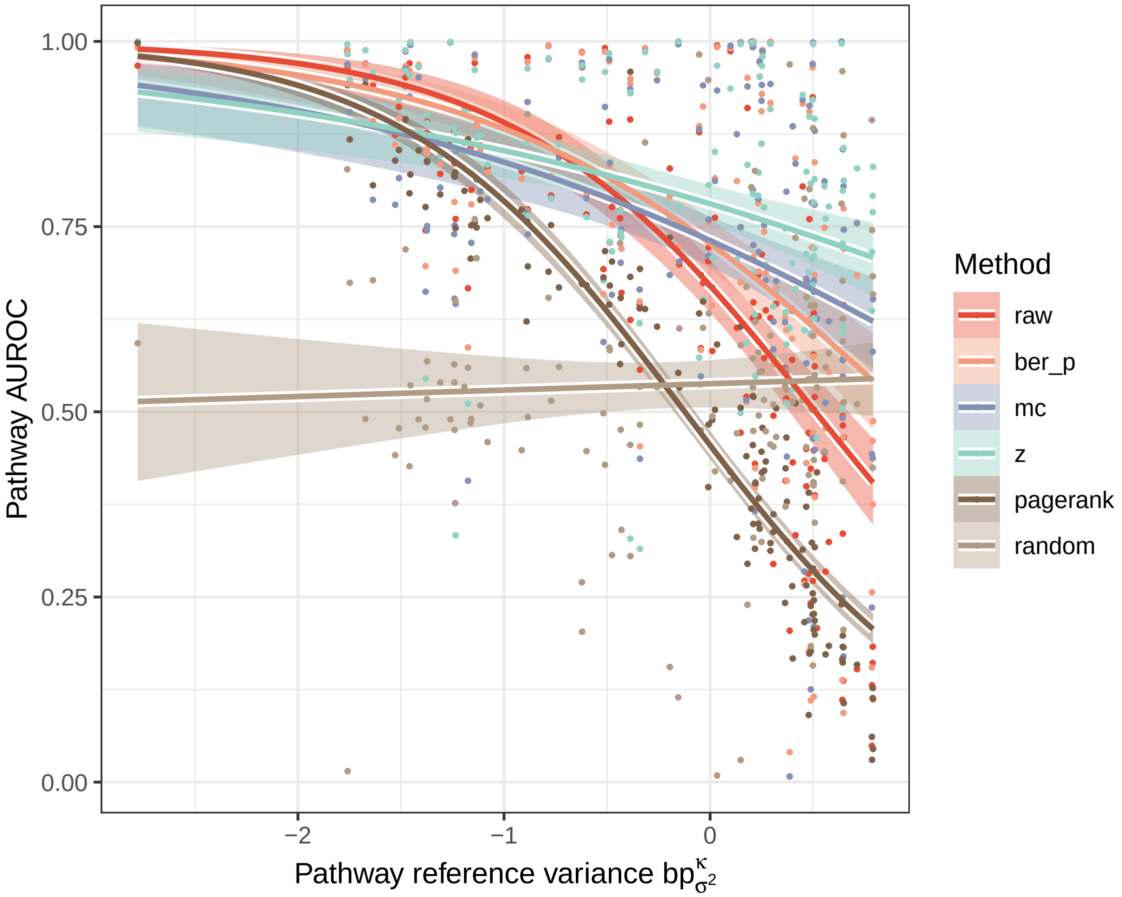

Figure 10: Prediction of the AUROC model by method along the reference pathway variance, represented by path_var_ref. Shaded are the 0.95 confidence intervals for the predicted mean AUROC.

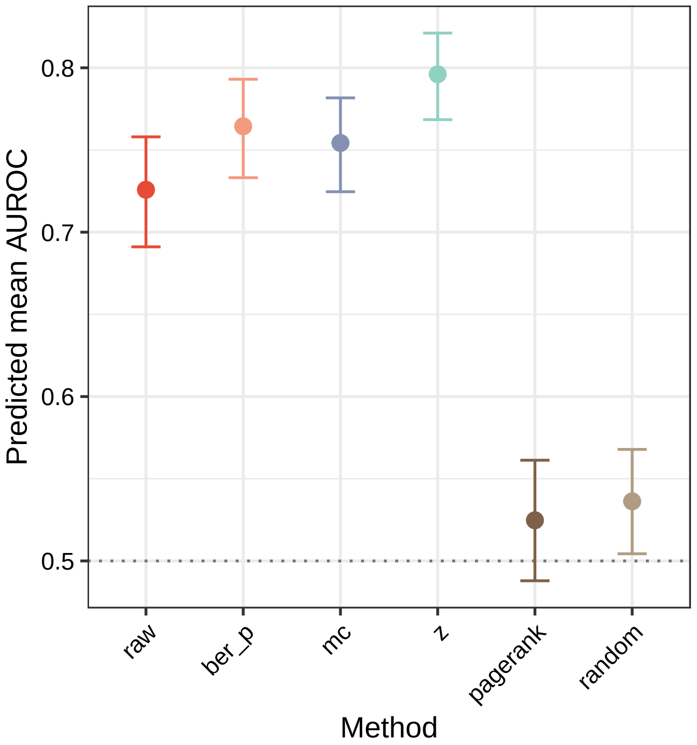

Figure 11: Predictions using the AUROC model (0.95 confidence intervals). Predictions were averaged over the path_var_ref covariate.

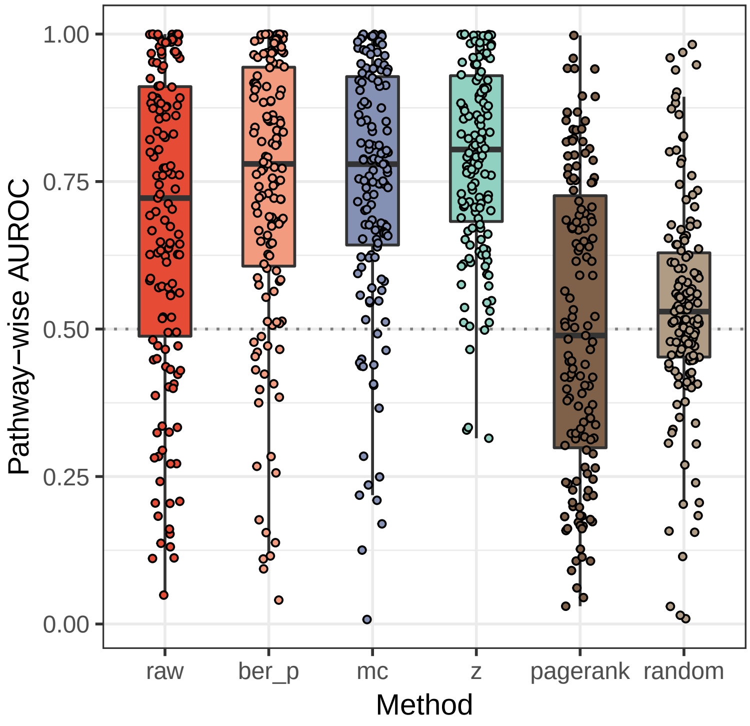

Figure 12: AUROC for all the pathways, by method.

#### 3.3 AUPRC

The model on AUPRC pointed out that early retrieval was challenging in this prospective study. Even though proper methods did outperform the baselines, performances were low, also due to the heavy class imbalance.

Table 3 describes the `quasilogistic` model – differences were minimal between methods, lacking statistical support. Furthermore, figure 14 proves how the 0.95 confidence intervals on the mean value of `path_var_ref` are overlapping.

Besides, AUPRC is affected by the class imbalance, meaning that pathways with few new genes were expected to yield low values of AUPRC.

Due to the two reasons above, AUPRC was not useful to describe differences between methods, but to highlight the difficult nature of this prospective analysis. The fact that an old network was used rules out possible circularities, i.e. the new genes being included in the pathways and in new interactions, based on the same data source.

Table 3: Quasibinomial model for AUPRC

| | |
|---|---|
| methodber_p | 0.021 (−0.488, 0.531) |
| methodmc | $-1.054^{***}$ (−1.783, −0.326) |
| methodz | $-0.554^{*}$ (−1.169, 0.062) |
| methodpagerank | $-3.246^{***}$ (−5.171, −1.320) |
| methodrandom | $-3.656^{***}$ (−5.917, −1.395) |
| methodraw:path_var_ref | 0.002 (−0.450, 0.454) |
| methodber_p:path_var_ref | 0.008 (−0.441, 0.456) |
| methodmc:path_var_ref | $-0.651^{**}$ (−1.239, −0.063) |
| methodz:path_var_ref | $-0.673^{***}$ (−1.137, −0.209) |
| methodpagerank:path_var_ref | −1.062 (−2.507, 0.383) |
| methodrandom:path_var_ref | −0.332 (−2.709, 2.045) |
| Constant | $-3.239^{***}$ (−3.601, −2.877) |
| Observations | 834 |
| *Note:* | $^{*}$p<0.1; $^{**}$p<0.05; $^{***}$p<0.01 |

Performing a paired Wilcoxon test yielded no evidence that `raw` and `z` had different AUPRCs (Table 4). The fact that `mc` was actually performing slighly worse than the rest of methods was deemed uninformative, given the actual magnitude of the effect and the overall low AUPRCs.

For completeness, we checked for significant differences between `raw` and `z` in the interaction term, because table 3 may suggest that `z` could be more affected than `raw` by the reference variances. In line with the other results, this counterintuitive claim could not be proven after a contrast on the interaction term `method:path_var_ref`:

```
##  contrast              estimate        SE  df z.ratio p.value
##  raw - ber_p       -0.005667778 0.3248737 Inf  -0.017  1.0000
##  raw - mc           0.652724459 0.3784352 Inf   1.725  0.5152
```

| | raw | ber_p | mc | z | pagerank | random |
|---|---|---|---|---|---|---|
| raw | | -8e-05(-0.00025,1.1e-05) | 0.0011(4e-04,0.0029) | 8.2e-05(-0.00023,0.00072) | 0.0066(0.0035,0.012) | 0.008(0.0047,0.017) |
| ber_p | 4.65e-02 | | 0.0013(0.00052,0.0028) | 0.00017(-0.00011,0.00075) | 0.0067(0.0035,0.013) | 0.0083(0.0048,0.018) |
| mc | 1.59e-03 | 1.45e-05 | | -8e-04(-0.0026,-0.00014) | 0.0046(0.0021,0.0065) | 0.0055(0.0032,0.0089) |
| z | 7.01e-01 | 3.06e-01 | 3.21e-03 | | 0.0064(0.0035,0.01) | 0.0069(0.0039,0.012) |
| pagerank | 2.50e-17 | 8.34e-18 | 3.79e-15 | 2.56e-19 | | 5.3e-05(-4.1e-05,4e-04) |
| random | 1.08e-17 | 8.82e-19 | 2.81e-18 | 7.87e-19 | 2.34e-01 | |

Table 4: Paired two-sided Wilcoxon test between AUPRCs, corrected by FDR. Above diagonal: differences with 0.95 confidence interval. Below diagonal: FDR.

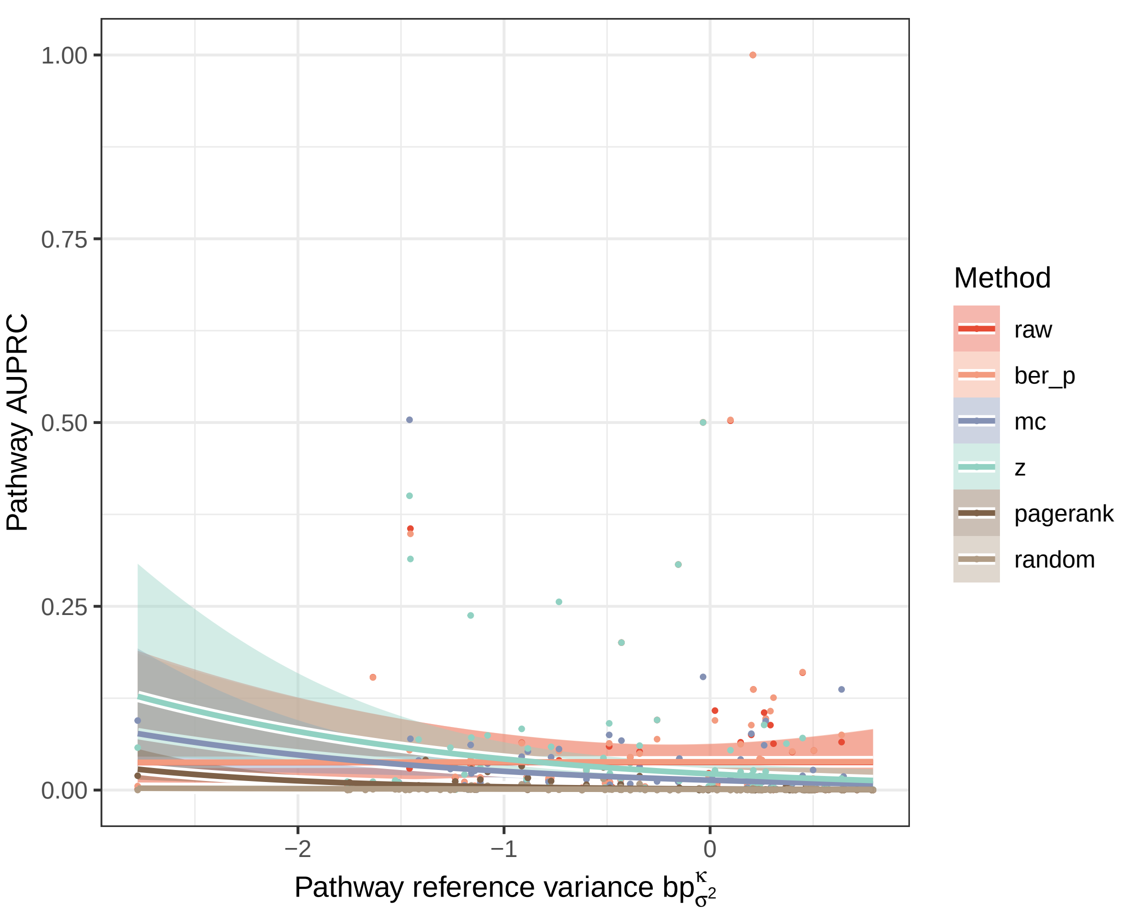

Figure 13: Prediction of the AUPRC model by method along the reference pathway variance, represented by path_var_ref. Shaded are the 0.95 confidence intervals for the predicted mean AUPRC.

```
##  raw - z            0.674948860 0.3303838 Inf   2.043  0.3179
##  raw - pagerank     1.064137045 0.7723649 Inf   1.378  0.7405
##  raw - random       0.333723339 1.2343521 Inf   0.270  0.9998
##  ber_p - mc         0.658392237 0.3773436 Inf   1.745  0.5019
##  ber_p - z          0.680616638 0.3291329 Inf   2.068  0.3042
##  ber_p - pagerank   1.069804823 0.7718306 Inf   1.386  0.7355
##  ber_p - random     0.339391117 1.2340179 Inf   0.275  0.9998
##  mc - z             0.022224401 0.3820977 Inf   0.058  1.0000
##  mc - pagerank      0.411412586 0.7958597 Inf   0.517  0.9955
##  mc - random       -0.319001120 1.2491879 Inf  -0.255  0.9999
##  z - pagerank       0.389188185 0.7741660 Inf   0.503  0.9961
##  z - random        -0.341225521 1.2354800 Inf  -0.276  0.9998
##  pagerank - random -0.730413706 1.4190859 Inf  -0.515  0.9956
##
## P value adjustment: tukey method for comparing a family of 6 estimates
```

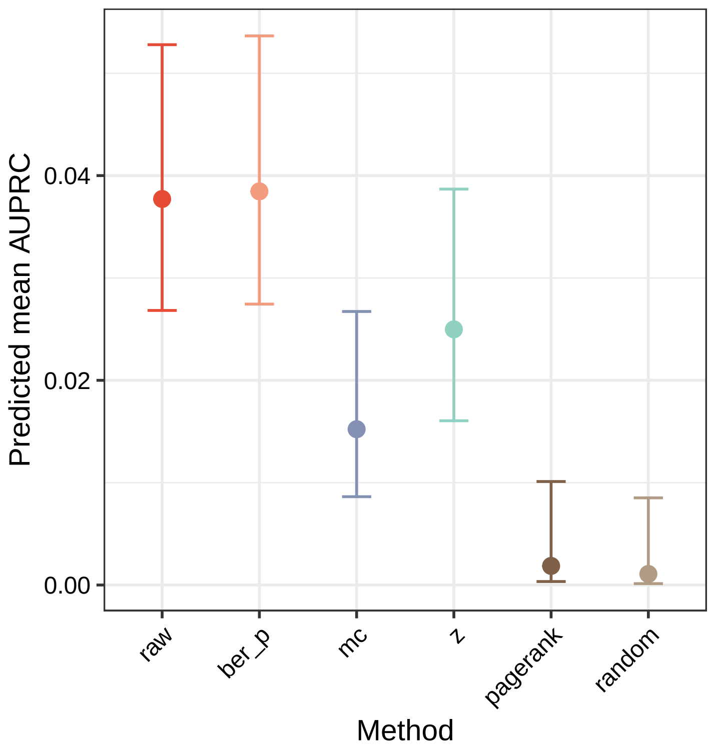

Figure 14: Predictions using the AUPRC model (0.95 confidence intervals). Predictions were averaged over the path_var_ref covariate.

##### 3.4 Other remarks

The present case study served as an illustrative example of variance-related bias in diffusion scores.

- The effect of the bias correction was not as straightforward as the mean value-related bias. We hypothesised that `z` would have more power on low-variance nodes compared to `raw`, but our findings support the opposite. The counterintuitive nature of this bias encourages an additional layer of caution.
- Normalising the diffusion scores led to a more bias-independent power for AUROC, in line with our hypothesis.
- AUROC was more informative than AUPRC and helped identify bias-related trends in predictive power.
- Again, the overall performance, and therefore the decision on normalising, relied on the distribution of the positives with respect to the reference variance. In this particular instance, `z` outperformed `raw`.
- For all the methods, new positives with higher variances were harder to recover, although this was less pronounced in `z`. High variance nodes tended to have a low degree, so we speculate that the network was incomplete when describing their biology, thus limiting the performance in their respective pathways.

### 4 Reproducibility

```
##  [1] "R version 3.5.3 (2019-03-11)"
##  [2] "Platform: x86_64-pc-linux-gnu (64-bit)"
##  [3] "Running under: Ubuntu 16.04.6 LTS"
##  [4] ""
##  [5] "Matrix products: default"
##  [6] "BLAS: /usr/lib/atlas-base/atlas/libblas.so.3.0"
##  [7] "LAPACK: /usr/lib/atlas-base/atlas/liblapack.so.3.0"
##  [8] ""
##  [9] "locale:"
## [10] " [1] LC_CTYPE=en_US.UTF-8       LC_NUMERIC=C              "
## [11] " [3] LC_TIME=en_US.UTF-8        LC_COLLATE=en_US.UTF-8    "
## [12] " [5] LC_MONETARY=en_US.UTF-8    LC_MESSAGES=en_US.UTF-8   "
## [13] " [7] LC_PAPER=en_US.UTF-8       LC_NAME=C                 "
## [14] " [9] LC_ADDRESS=C               LC_TELEPHONE=C            "
## [15] "[11] LC_MEASUREMENT=en_US.UTF-8 LC_IDENTIFICATION=C       "
## [16] ""
## [17] "attached base packages:"
## [18] "[1] grid      stats     graphics  grDevices utils     datasets  methods  "
## [19] "[8] base     "
## [20] ""
## [21] "other attached packages:"
## [22] " [1] bindrcpp_0.2.2   xtable_1.8-3     extrafont_0.17   gtable_0.2.0    "
## [23] " [5] ggsci_2.9        ggplot2_3.1.0    stargazer_5.2.2  emmeans_1.3.0   "
## [24] " [9] magrittr_1.5     tidyr_0.8.2      dplyr_0.7.8      plyr_1.8.4      "
## [25] "[13] reshape2_1.4.3   diffuStats_1.2.0 igraph_1.2.2    "
## [26] ""
## [27] "loaded via a namespace (and not attached):"
## [28] " [1] zoo_1.8-4                 tidyselect_0.2.5         "
## [29] " [3] xfun_0.4                  purrr_0.2.5              "
## [30] " [5] splines_3.5.3             lattice_0.20-38          "
## [31] " [7] colorspace_1.3-2          expm_0.999-3             "
## [32] " [9] htmltools_0.3.6           yaml_2.2.0               "
## [33] "[11] survival_2.43-3           rlang_0.3.0.1            "
## [34] "[13] pillar_1.3.0              withr_2.1.2              "
## [35] "[15] glue_1.3.0                multcomp_1.4-8           "
## [36] "[17] bindr_0.1.1               stringr_1.3.1            "
## [37] "[19] munsell_0.5.0             mvtnorm_1.0-8            "
## [38] "[21] codetools_0.2-16          evaluate_0.12            "
## [39] "[23] labeling_0.3              knitr_1.20               "
## [40] "[25] RcppArmadillo_0.9.200.4.0 Rttf2pt1_1.3.7           "
## [41] "[27] TH.data_1.0-9             Rcpp_1.0.0               "
## [42] "[29] scales_1.0.0              backports_1.1.2          "
## [43] "[31] RcppParallel_4.4.1        precrec_0.9.1            "
## [44] "[33] digest_0.6.18             stringi_1.2.4            "
## [45] "[35] bookdown_0.7              rprojroot_1.3-2          "
## [46] "[37] tools_3.5.3               sandwich_2.5-0           "
## [47] "[39] lazyeval_0.2.1            tibble_1.4.2             "
## [48] "[41] extrafontdb_1.0           crayon_1.3.4             "
## [49] "[43] pkgconfig_2.0.2           MASS_7.3-51.1            "
## [50] "[45] Matrix_1.2-15             data.table_1.11.8        "
## [51] "[47] estimability_1.3          assertthat_0.2.0         "
## [52] "[49] rmarkdown_1.10            R6_2.3.0                 "
```

```
## [53] "[51] compiler_3.5.3            "
```

Carlson, Marc. 2016. *KEGG.db: A Set of Annotation Maps for Kegg.*

Chatr-aryamontri, Andrew, Rose Oughtred, Lorrie Boucher, Jennifer Rust, Christie Chang, Nadine K Kolas, Lara O'Donnell, et al. 2017. "The Biogrid Interaction Database: 2017 Update." *Nucleic Acids Research* 45 (D1). Oxford University Press: D369–D379.

Kanehisa, Minoru, Miho Furumichi, Mao Tanabe, Yoko Sato, and Kanae Morishima. 2017. "KEGG: New Perspectives on Genomes, Pathways, Diseases and Drugs." *Nucleic Acids Research* 45 (D1). Oxford University Press: D353–D361.
